# Comparative multi-omics of the macrophage response to infection with *Mycobacterium tuberculosis* complex bacteria reveals pathogen-driven epigenomic reprogramming

**DOI:** 10.64898/2026.02.15.705989

**Authors:** Thomas J. Hall, Morgane Mitermite, John F. O’Grady, John A. Browne, Gillian P. McHugo, Emily L. Clark, Mazdak Salavati, Stephen V. Gordon, David E. MacHugh

## Abstract

**Background:** Bovine tuberculosis (bTB) is a chronic infectious disease primarily caused by *Mycobacterium bovis*, which inflicts significant economic losses on the global livestock industry worldwide and can also cause tuberculosis (TB) disease in other mammalian species, including humans. Alveolar macrophages are the host cells targeted by the pathogen during the early stages of infection. While they play a crucial role in controlling infection, the exact nature of the host-pathogen interaction and the genetic and epigenetic factors that modulate infection outcome remain poorly understood.

**Results:** Here, we used transcriptomics (RNA-seq), chromatin accessibility (ATAC-seq), and chromatin configuration (ChIP-seq) analyses to examine the effects of intracellular mycobacterial infection on the bovine alveolar macrophage (bAM) transcriptome and epigenome. The primary focus was *M. bovis* infection, but we also conducted parallel comparative analyses using *M. tuberculosis* (the primary cause of human TB—hTB), *M. bovis* BCG (the vaccine strain), and gamma-irradiated (killed) *M. bovis*. Integration of RNA-seq, ChIP-seq, and ATAC-seq data revealed coordinated remodelling of chromatin accessibility and histone modification landscapes underpinning transcriptional activation of key immune and metabolic pathways in response to infection. The identification of candidate genes, including *ERBB4*, *LRCH1*, *MRTFA*, and *RNPC3*, through integrative analysis with a genome-wide association study (GWAS) for *M. bovis* infection susceptibility underscores the functional relevance of these regulatory networks.

**Conclusions:** Our results demonstrate that *M. bovis* drives extensive reprogramming of the bAM epigenome, distinct from the responses elicited by other members of the *M. tuberculosis* complex (MTBC). The results of this multi-omics comparison provide new insights into the function of pivotal response genes and support the hypothesis that pathogen-driven epigenetic reprogramming of the bovine host macrophage is key to *M. bovis* survival. It also identifies molecular targets that may inform genome-enabled breeding strategies to enhance resilience to bTB in cattle.

## Introduction

Bovine tuberculosis (bTB) is a chronic infectious disease that primarily affects livestock, particularly taurine (*Bos taurus*), zebu (*Bos indicus*), and *B. taurus* × *indicus* hybrid cattle populations. Bovine TB has a substantial socioeconomic cost through disrupting food production, threatening stockholder livelihoods, and causing economic damage to global agriculture that runs to billions of dollars annually (Zinsstag et al. 2006; Azami and Zinsstag 2018). The disease is primarily caused by *Mycobacterium bovis*, an intracellular mycobacterial pathogen with 99.95% genome sequence identity to *Mycobacterium tuberculosis*, the primary cause of human tuberculosis (hTB) (Garnier et al. 2003). However, unlike *M. tuberculosis, M. bovis* has a broad host range and can infect several important domestic species besides cattle, including sheep (*Ovis aries*), goats (*Capra hircus*), pigs (*Sus scrofa*), and llamas and alpacas (*Lama* spp.). (Pesciaroli et al. 2014; Malone and Gordon 2017). Importantly, *M. bovis* can also cause zoonotic tuberculosis (zTB), which has serious implications for human health, particularly in the Global South (Olea-Popelka et al. 2017; Vayr et al. 2018; Kock et al. 2021). *Mycobacterium bovis* and *M. tuberculosis* are both classified within a broader taxonomic group, the *Mycobacterium tuberculosis* complex (MTBC), consisting of closely related intracellular mycobacterial species that can propagate and transmit within a spectrum of animal hosts (Malone and Gordon 2017).

Previous research has shown that the pathogenesis of bTB is similar to hTB. There are many shared characteristics between *M. tuberculosis* infection in humans and *M. bovis* infection in cattle, such that bTB can serve as a valuable large animal model for TB disease in humans (Waters et al. 2014; Buddle et al. 2016; Williams and Orme 2016; Gong et al. 2020). Transmission of tubercle bacilli occurs through inhalation of contaminated aerosol droplets, with the primary site of infection establishing in the lungs. Tissue-resident alveolar macrophages (AM) normally phagocytose airborne bacteria and other microorganisms to contain or destroy them (Aegerter et al. 2022; Park et al. 2022). Disease-causing mycobacteria, however, have evolved various mechanisms to interfere with host immune responses and persist and replicate within AM (Cambier et al. 2014; Schorey and Schlesinger 2016; Awuh and Flo 2017; Chandra et al. 2022; Bo et al. 2023). These mechanisms include the manipulation of cell surface receptors on host macrophages, inhibition of phagosome-lysosome fusion, neutralisation of reactive oxygen and nitrogen intermediates (ROI and RNI), exploitation of intracellular nutrient supply and metabolism, suppression of apoptosis and autophagy, interference with antigen presentation, modulation of macrophage signalling pathways, cytosolic escape from the phagosome, and induction of necrosis. These actions contribute to immunopathology, pathogen shedding from the host, and overall disease progression (Ehrt and Schnappinger 2009; Hussain Bhat and Mukhopadhyay 2015; Queval et al. 2017; Chaurasiya 2018; Stutz et al. 2018).

The discovery of substantial reprogramming of AM transcriptional networks in response to *M. bovis* and *M. tuberculosis* infection exemplifies the profound impact of intracellular mycobacteria on host macrophages (Nalpas et al. 2015; Vegh et al. 2015; Lavalett et al. 2017; Jensen et al. 2018; Malone et al. 2018; Papp et al. 2018; Hall et al. 2019; Lavalett et al. 2020a; Pisu et al. 2020; Hall et al. 2021; Hall et al. 2024). These investigations have also revealed that differentially expressed genes (DEGs) and perturbed cellular pathways and networks are functionally associated with macrophage processes that control or eliminate intracellular microbes. It has also been shown that the rewiring of host gene expression induced by intracellular pathogens to promote transmission and persistence can involve reconfiguring and remodelling of chromatin (Hamon and Cossart 2008; Zhang and Cao 2019; Fol et al. 2020). This phenomenon has been observed in infections caused by various bacterial pathogens, including *Chlamydia* spp., *Leishmania donovani*, *Listeria monocytogenes*, and *Yersinia enterocolitica* (Marr et al. 2014; Bierne and Hamon 2020; Bekere et al. 2021; Stein and Thompson 2023).

Regarding intracellular mycobacterial infections and the host epigenome, an early study by Yaseen and colleagues (Yaseen et al. 2015) demonstrated that the Rv1988 protein, secreted by virulent mycobacteria, localises to chromatin during infection and represses host cell genes by methylating histone H3 at a non-canonical arginine residue. Additionally, chromatin immunoprecipitation sequencing (ChIP-seq) analysis revealed that regulatory sequence motifs embedded in specific subtypes of Alu SINE transposable elements play a significant role in the epigenetic machinery that modulates human macrophage gene expression during *M. tuberculosis* infection (Bouttier et al. 2016). More recently, it has been demonstrated that heterologous expression of the mycobacterial PE family protein PE17 gene (*PE17*/*Rv1646*) in a *Mycobacterium smegmatis*-infected macrophage cell line promotes activation of apoptotic signalling pathways via remodelling of host chromatin through reduced H3K9me3 chromatin occupancy (Abo-Kadoum et al. 2021). Correa-Macedo and coworkers have also shown, using a combination of RNA-seq, ATAC-seq, and ChIP-seq, that primary human AM (hAM) obtained using bronchoalveolar lavage (BAL) exhibited altered transcriptional programs and substantially remodelled chromatin when challenged *in vitro* with *M. tuberculosis* (Correa-Macedo et al. 2021). Intriguingly, this work also demonstrated that bAM from HIV-positive subjects and HIV-negative subjects undergoing prophylactic antiretroviral therapy (ART) had a profoundly blunted transcriptional response with little or no changes in chromatin accessibility. In addition, two different studies of THP-1 differentiated macrophages infected with *M. tuberculosis* using transcriptomics with ATAC-seq and single-cell Hi-C, respectively, showed significant reorganisation of chromatin initiated by NF-κB-directed transcriptional regulation (Lin et al. 2022) and impacts on the expression of genes involved in type I interferon (IFN) signalling pathways (Madden et al. 2023).

Host-pathogen interaction at the level of the macrophage epigenome has also been studied for bTB; our group has previously shown that bovine alveolar macrophages (bAM) infected with *M. bovis* undergo extensive transcriptional reprogramming through differential distribution of H3K4me3 and Pol II at key immune genes (Hall et al. 2019). Furthermore, we found evidence suggesting that reconfigured chromatin at some genomic loci may be favourable for intracellular bacterial survival, such as the triple methylation of H3K4 at the transcriptional start site (TSS) of the *TNFAIP3* gene, which encodes a key inhibitor of the innate immune-activating NF-kB pathway. This immunoevasion strategy has also been observed for *Leishmania amazonensis*, where the same modification is seen at *TNFAIP3* in parasite-infected macrophages, but not in macrophages stimulated with LPS (Lecoeur et al. 2020).

The present study aims to extend our understanding of macrophage reprogramming in response to MTBC infection through a comparative multi-omics approach. To achieve this, we used RNA-seq, ChIP-seq, and ATAC-seq to examine gene expression changes and chromatin remodelling in bAM in response to four separate infections: (i) virulent *M. bovis*, the primary agent of bTB and the bovine-adapted MTBC species; (ii) *M. tuberculosis*, also virulent and closely related to *M. bovis* but the human-adapted MTBC species; (iii) *M. bovis* Bacillus Calmette-Guérin (BCG), the attenuated vaccine strain which lacks the RD1 locus, a major virulence determinant required for ESX-1 secretion and pathogenicity in both *M. tuberculosis* and *M. bovis*; and (iv) γ-irradiated *M. bovis*, dead bacteria that retain an intact cell wall and associated surface ligands for engagement of macrophage receptors but that lack secretion of virulence effectors. Taken together, these strains provided a framework for comparing the bovine alveolar macrophage transcriptional and chromatin responses to infection with virulent, attenuated/dead, and host-adapted mycobacteria. Furthermore, we integrated outputs from these analyses with large genome-wide association study (GWAS) datasets focused on an *M. bovis* infection-susceptibility trait to prioritise cattle genomic regions and SNPs associated with resilience to bTB.

## Results

### *In vitro* challenge of bovine alveolar macrophages with four types of MTBC mycobacteria

We investigated reprogramming of the bAM regulatory genome after *a* 24 h *in vitro* challenge with four different types of MTBC bacteria (*M. bovis* ‒ MBO; *M. tuberculosis* ‒ MTU; *M. bovis* BCG ‒ BCG; gamma-irradiated (killed) *M. bovis* ‒ IRR). We used a combined multi-omics approach, including RNA-seq, ChIP-seq, and ATAC-seq analyses, comparing each challenged bAM group to a control non-challenged bAM group (CON) cultured for 24 h post-infection (hpi) (Additional file 2: Table S1). A diagrammatic summary overview of the experimental design and analytical workflow is provided (Fig. 1).

**Fig. 1.**
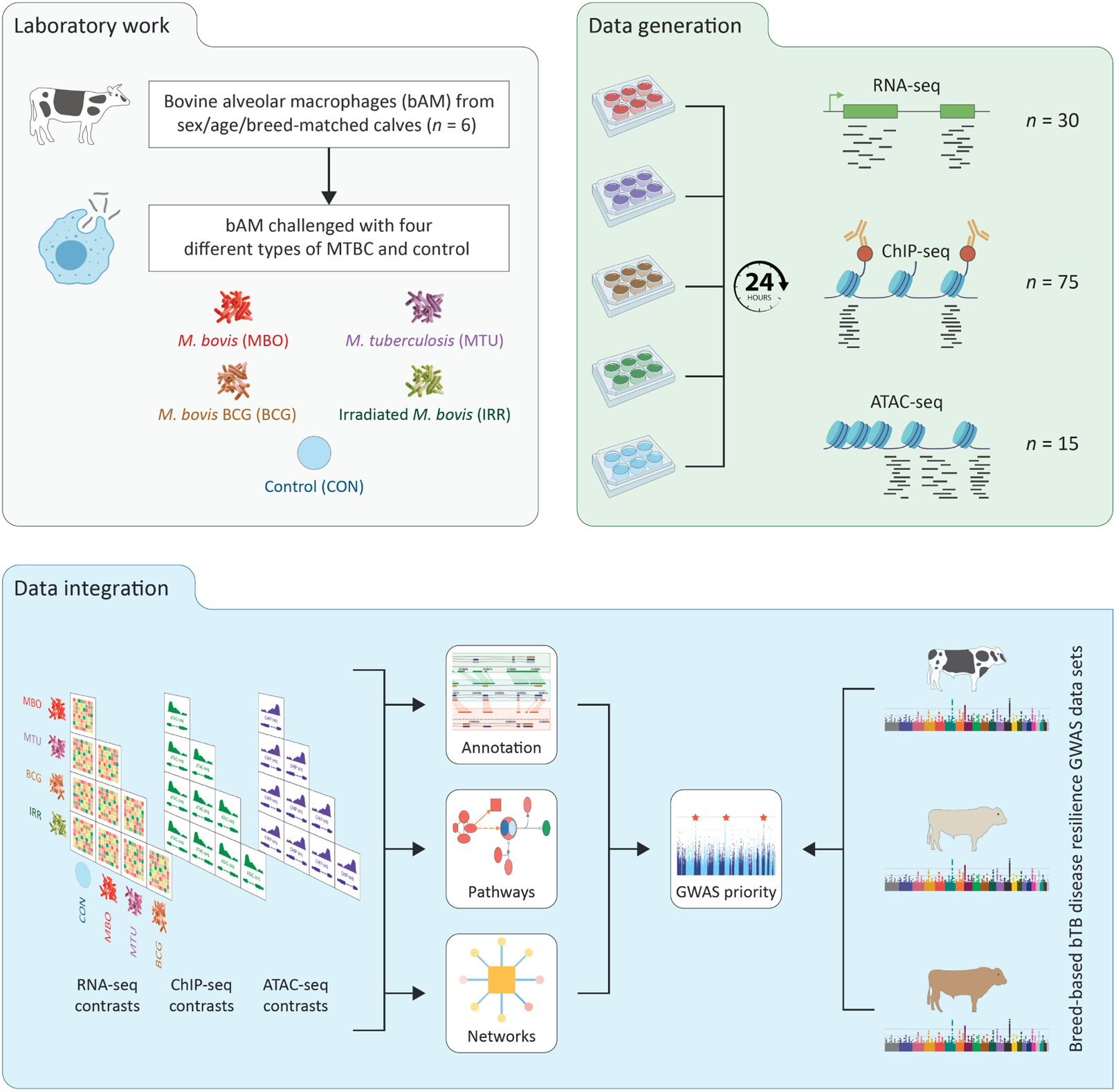
Experimental and computational workflow. Laboratory bovine alveolar macrophage (bAM) challenge, data generation, and data integration steps are shown.

### Bacterial uptake by cultured bovine alveolar macrophages

Fluorescence microscopy was used to generate 10 images of tissue culture plate wells containing the bAM samples for each experimental challenge group at 24 hpi (MBO, MTU, BCG, IRR, and CON). Combined analysis of these images yielded quantitative infection/uptake values, including bAM counts, bacterial cell counts, infected and uninfected cell counts, global multiplicity of infection (MOI) values, and the percentage of infected cells for each experimental challenge group (Additional file 2: Table S2). These results indicated that the relative levels of uninfected and infected cells did not differ significantly among the MBO, MTU, and BCG groups, with a mean 34.5% infection rate for the MBO group (*n* = 6), 33.3% for the MTU group (*n* = 6) and 35.0% for the BCG group (*n* = 6). The exception was the IRR group (*n* = 6), which had a 17.8% uptake rate for these dead bacterial cells. Examples of the images used to generate these quantitative infection/uptake data are provided (Additional file 1: Fig. S1a–e). A *t*-test was performed for each comparison, yielding FDR-*P*_adj._ > 0.05 for the three comparisons involving live bacteria and FDR-*P*_adj._ < 0.05 for the three comparisons with live bacteria versus the IRR group (Additional file 1: Fig. S1f).

### Transcriptional responses of bovine alveolar macrophages to different MTBC mycobacteria challenges

RNA-seq read and mapping statistics and raw gene counts were tabulated (Methods; Additional file 3: Tables S3–S4). The RNA-seq results from the five experimental groups (MBO, MTU, BCG, IRR, and CON) showed distinct transcriptomic responses (Additional file 3: Tables S5‒S8). A single MBO RNA-seq sample (animal ID: 43095) was compromised by contamination and not included in subsequent analyses (Additional file 2: Table S1). Using the PlotPCA function in DESeq2, we generated a plot of the first two principal components (PC1 and PC2) from a principal component analysis (PCA) of the normalised gene expression data for each of the 35 experimental samples (MBO, *n* = 5; MTU, *n* = 6; BCG, *n* = 6; IRR, *n* = 6; and CON, *n* = 6) (Fig. 2a). The values for PC1, which accounted for 39% of the variance, broadly separated the five experimental groups, with the CON group followed sequentially by the IRR, BCG, MTU, and MBO groups, corresponding approximately to the level of bAM transcriptome perturbation caused by each MTBC challenge. The PC2 results, on the other hand, which accounted for 20% of the variance, were largely determined by genetic variation and differentiated the individual biological samples (sample numbers 1‒6) in approximately the same sequential order within each experimental group.

**Fig. 2.**
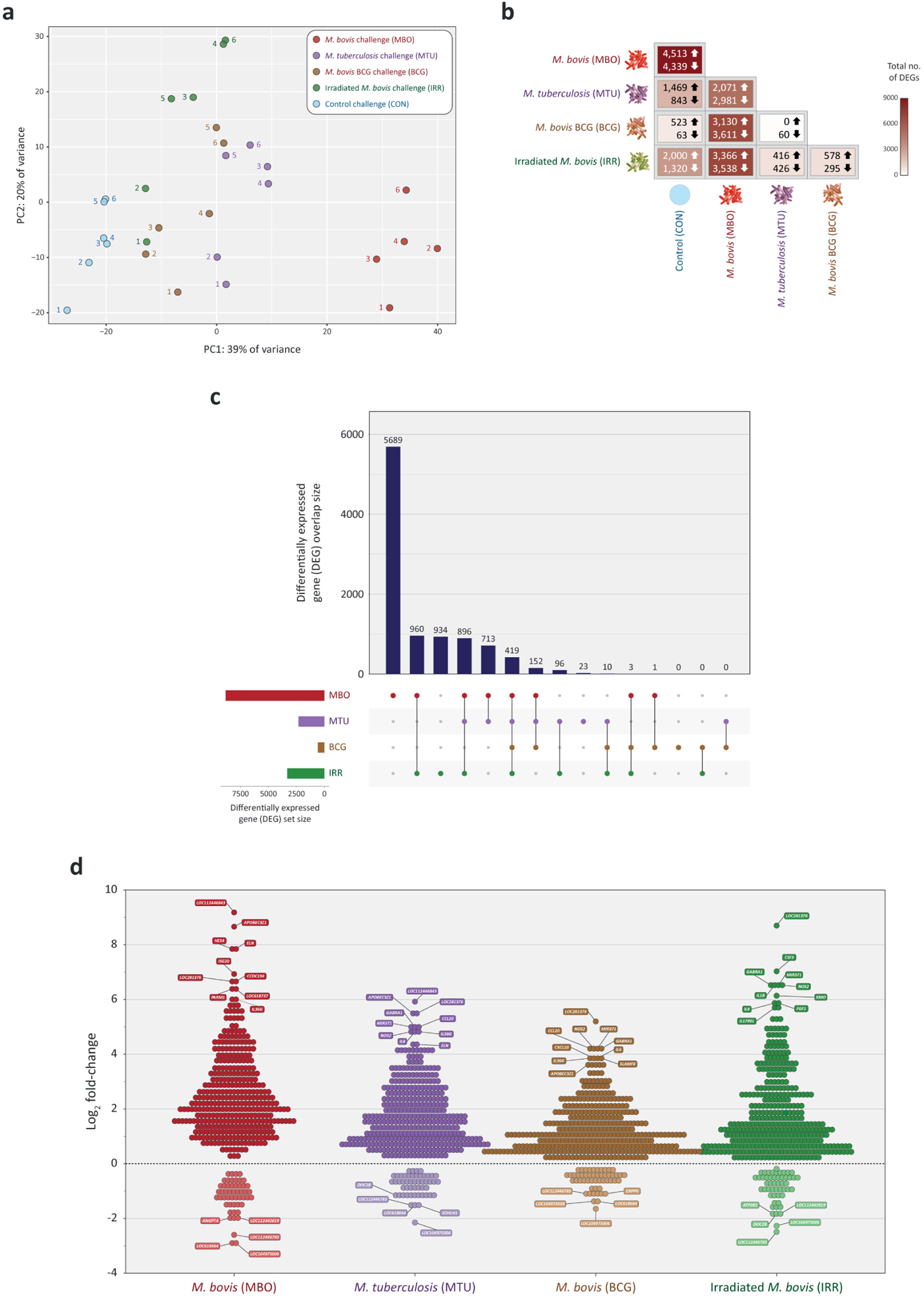
Gene expression and differentially expressed genes (DEGs) in bovine alveolar macrophages (bAM) challenged with *Mycobacterium bovis* (MBO), *M. tuberculosis* (MTU), *M. bovis* BCG (BCG), gamma-irradiated (killed) *M. bovis* (IRR), and a non-challenged control (CON). **a** PCA summary of gene expression data across the five challenge groups. **b** Numbers of DEGs (FDR-*P*_adj._ < 0.05; |log_2_FC| > 0| for all 10 possible contrasts. **c** UpSet plot showing the intersection of shared DEGs across the four primary bAM challenges (MBO/CON, MTU/CON, BCG/CON, and IRR/CON). **d** Binned jitter plot of log_2_FC values for the 419 significant DEGs across the four primary bAM challenges (MBO/CON, MTU/CON, BCG/CON, and IRR/CON).

A matrix heatmap of differentially expressed genes (DEGs) detected using the default criterion for differential expression (FDR-*P*_adj._ < 0.05) and partitioned into up- and downregulated genes for the ten experimental group contrasts across the five experimental groups (MBO, MTU, BCG, IRR, and CON) was generated (Fig. 2b). The numbers of DEGs detected ranged between 60 and almost 9,000 (8,852) for the BCG/MTU and MBO/CON contrasts, respectively. An UpSet plot, with the intersection DEG sets across the ten gene expression contrasts showed the overlaps in DEG sets among the different experimental contrasts (Fig. 2c). For example, of the 8,852 DEGs in the MBO/CON comparison, 5,689 DEGs were specific to bAM infected with *M. bovis* (i.e., only detected in the MBO/CON contrast); 960 DEGs intersected only for bAM challenged with the gamma-irradiated *M. bovis* (i.e., detected solely in the MBO/CON and IRR/CON contrasts); 713 intersected only for bAM infected with *M. tuberculosis* (i.e., detected solely in the MBO/CON and MTU/CON contrasts); and one DEG intersected only for bAM infected with *M. bovis* BCG (i.e., detected solely in the MBO/CON and BCG/CON contrasts). In total, across the experimental contrasts, there were 9,916 unique DEGs, with substantial overlaps among these DEGs across the four challenge-versus-control contrasts (MBO/CON, MTU/CON, BCG/CON, and IRR/CON). For this universal set of 9,916 DEGs, 419 were common to these four experimental groups (Additional file 3: Table S9) and a jitter plot was generated with the log_2_ fold-change (log_2_FC) values for these 419 genes with the top ten upregulated and top five downregulated genes highlighted (Fig. 2d). The top ten upregulated genes shared across the four contrasts included, for example, the *IL36G* gene (MBO/CON, MTU/CON, and BCG/CON), which encodes IL-36γ a cytokine that has been shown to induce bactericidal processes in *M. tuberculosis*-infected macrophages (Ahsan et al. 2016; Gao et al. 2019). Also highlighted were *CCL20* (MTU/CON and BCG/CON), which encodes a key chemokine in the immunoregulatory and inflammatory response to intracellular mycobacterial pathogens (Rivero-Lezcano et al. 2010; Lavalett et al. 2020b); *IL6* (MTU/CON, BCG/CON, and IRR/CON), which encodes the proinflammatory IL-6 cytokine produced by macrophages infected with MTBC pathogens (Martinez et al. 2013); and *NOS2* (MTU/CON, BCG/CON, and IRR/CON), which encodes a nitric oxide (NO) synthase that provides important antimicrobial functions in macrophages (Bogdan 2015).

Focusing on the four contrasts for the MBO, MTU, BCG and IRR bAM challenge groups versus the CON group individually, the following numbers of statistically significant DEGs (FDR-*P*_adj._ < 0.05) were detected: 8,852 (4,513 up, 4,339 down) for the MBO/CON contrast, 2,312 (1,469 up, 843 down) for the MTU/CON contrast, 586 (523 up, 63 down) for the BCG/CON contrast, and 3,320 (2,000 up, 1,320 down) for the IRR/CON contrast (Fig. 2b, Fig. 3a–d; Additional file 3: Tables S5–S8). These results show that the transcriptomic response relative to the CON group was substantially more pronounced in the MBO group, and that most DEGs in the MBO/CON contrast were upregulated. The volcano plots of DEGs (Fig. 3a–d) highlight additional important upregulated genes for each of the four contrasts (MBO/CON, MTU/CON, BCG/CON, and IRR/CON). These include *IFIT3* (MBO/CON and MTU/CON), which is induced by type I interferon (IFN) signalling and has been identified as an important hub gene in host transcriptional networks responding to *M. tuberculosis* infection and as a potential biomarker of active TB disease in humans (Qiu et al. 2022; Wu et al. 2023; Liu and Li 2024). Also highlighted is *IRF7* (MTU/CON and BCG/CON), which encodes a key regulator of type I IFN-dependent immune responses that promote increased mycobacterial intracellular survival (Leisching et al. 2017; Cheng and Schorey 2018). In addition, upregulation of *SP110* was evident (MTU/CON and BCG/CON), a gene that promotes apoptotic cell death and hence containment of infection (Wu et al. 2016) and is a positive regulator of inflammatory responses in *M. tuberculosis*-infected macrophages (Nakamura et al. 2024).

**Fig. 3.**
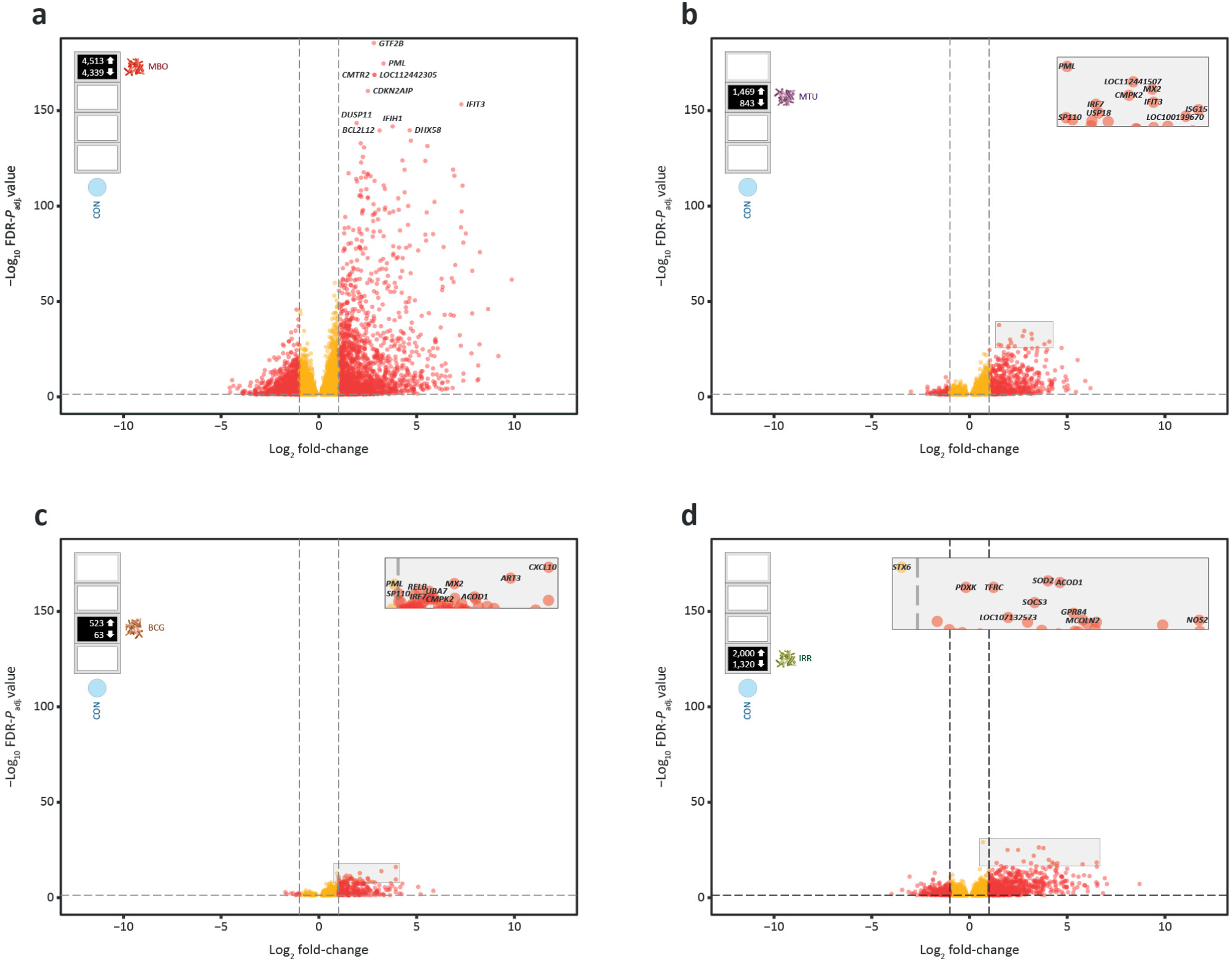
Volcano plots showing statistically significant (FDR-*P*_adj._ < 0.05) differentially expressed genes (DEGs) for the four primary MTBC bAM challenges. Orange data points represent genes with |log_2_FC values| < 1, and red data points denote genes with |log_2_FC values| ≥ 1. The top ten genes ranked by −log_10_ FDR-*P*_adj._ are also shown for each contrast. **a** *Mycobacterium bovis* vs. control (MBO/CON). **b** *M. tuberculosis* vs. control (MTU/CON). **c** *M. bovis* BCG vs. control (BCG/CON). **d** gamma-irradiated (killed) *M. bovis* vs. control (IRR/CON).

The DEG gene sets for the ten possible experimental group contrasts were also analysed using IPA^®^, and the top five pathways (by *P*-value) were examined (Methods; Additional file 3: Table S10). In addition, the top ten pathways for the four primary challenge contrasts (MBO/CON, MTU/CON, BCG/CON, and IRR/CON) were collated and visualised by activation state (Additional file 1: Fig. S2a‒d). For each pathway analysis, among the top five pathways, both unique pathways and overlaps across the ten contrasts were observed. The enriched pathways that were common to two or more groups included *Interferon signalling* and *Activation of IRF by cytosolic pattern recognition receptors* (MBO/CON, MTU/CON, BCG/CON, MBO/BCG, MBO/IRR, MBO/MTU, MTU/BCG, and MTU/IRR); *Role of hypercytokinemia/hyperchemokinemia in the pathogenesis of influenza* (MTU/CON, BCG/CON, BCG/IRR, MBO/IRR, MTU/BCG, and MTU/IRR); *Macrophage classical activation signalling pathway* (MTU/CON and BCG/CON); and *Role of RIG1-like receptors in antiviral innate immunity* (MBO/BCG, MBO/IRR, MBO/MTU, and MBO/BCG), which is in agreement with previous work highlighting the importance of RIG-I-like receptor signalling in the specific responses of *M. bovis*-infected bAM (Malone et al. 2018; Hall et al. 2019; Hall et al. 2021; Hall et al. 2024). Although unique pathways specific to one contrast were less common, two of the most notable examples were *Death receptor signalling* and *Protein ubiquitination pathway* (MBO/CON). Finally, all the top five pathways in the IRR/CON contrast were unique to that group: *IL-6 signalling, PPAR signalling, IL-10 signalling, Acute phase response signalling,* and *CD40 signalling*.

### Transcriptional reprogramming of bovine alveolar macrophages challenged with different MTBC mycobacteria is modulated by histone modifications and chromatin accessibility

Previous work has shown that bAM undergo marked transcriptional perturbation and reprogramming after *in vitro* challenge with *M. bovis* 24 hpi, with a substantial proportion of the detectable bAM transcriptome exhibiting significant differential expression compared to non-infected control bAM (Nalpas et al. 2015; Malone et al. 2018; Hall et al. 2021). The extent of these gene expression changes is comparable to those observed in studies that have also incorporated additional functional genomics assays to investigate chromatin remodelling during bacterial and viral infections. In these cases, the levels of trimethylation of lysine 4 of histone 3 (H3K4me3) and acetylation of lysine 27 of histone 3 (H3K27ac) were associated with active transcription, and monomethylation of lysine 4 of histone 3 (H3K4me1) and trimethylation of lysine 27 of histone 3 (H3K27me3) were associated with transcriptional repression (Abraham and Kulesza 2013; Bouttier et al. 2016; Arts et al. 2018; Hall et al. 2019; Del Rosario et al. 2022).

To expand our previous work examining chromatin configuration and reprogramming of the bAM transcriptome due to *M. bovis* infection (Hall et al. 2019), we used ChIP-seq and ATAC-seq in parallel to examine histone modifications (H3K4me3, H3K4me1, H3K27ac, and H3K27me3), CTCF binding, and broader regions of open chromatin (ATAC-seq) that occur in bAM 24 h after MBO, MTU, BCG, and IRR challenge (Methods). The bAM samples from three sex- and age-matched Holstein-Friesian cattle used for these experiments were a subset of the six biological replicates used for the RNA-seq gene expression analyses (Additional file 2: Table S1). Firstly, the ChIP-seq read and mapping statistics and detailed peak statistics were tabulated (Additional file 4: Tables S12‒S13), and analysis of the ChIP-seq data revealed a mean of 24,641 peaks for H3K4me3, a mark of active transcription, across the 15 challenge samples (3 MBO, 3 MTU, 3 BCG, 3 IRR, and 3 CON). In addition, a mean of 65,040 peaks was obtained for H3K4me1 (marks active and primed enhancers), 31,844 for H3K27ac (marks active enhancers), 5,044 for H3K27me3 (marks transcriptional repression of proximal genes), and 45,292 for CTCF protein binding (marks repression of enhancer-mediated transcriptional activation) (Additional file 4: Tables S14). For the H3K4me1 peaks, only two control (CON) samples were available, so the mean value across 14 samples was calculated (Additional file 4: Table S14). Secondly, the ATAC-seq read and mapping statistics and detailed peak statistics were tabulated (Additional file 5: Tables S19‒S20), and analysis of the ATAC-seq data revealed a mean of 90,540 peaks across the 15 challenge samples (3 MBO, 3 MTU, 3 BCG, 3 IRR, and 3 CON) (Additional file 5: Table S19). The H3K4me3, H3K4me1, H3K27ac, H3K27me3, and ATAC-seq genome-wide peak distributions for each experimental bAM challenge group (MBO, MTU, BCG, IRR, and CON) were plotted as nested circular signal tracks across the ARS-UCD1.2 bovine reference genome (BTA1–BTAX) (Fig. 4‒e).

**Fig. 4.**
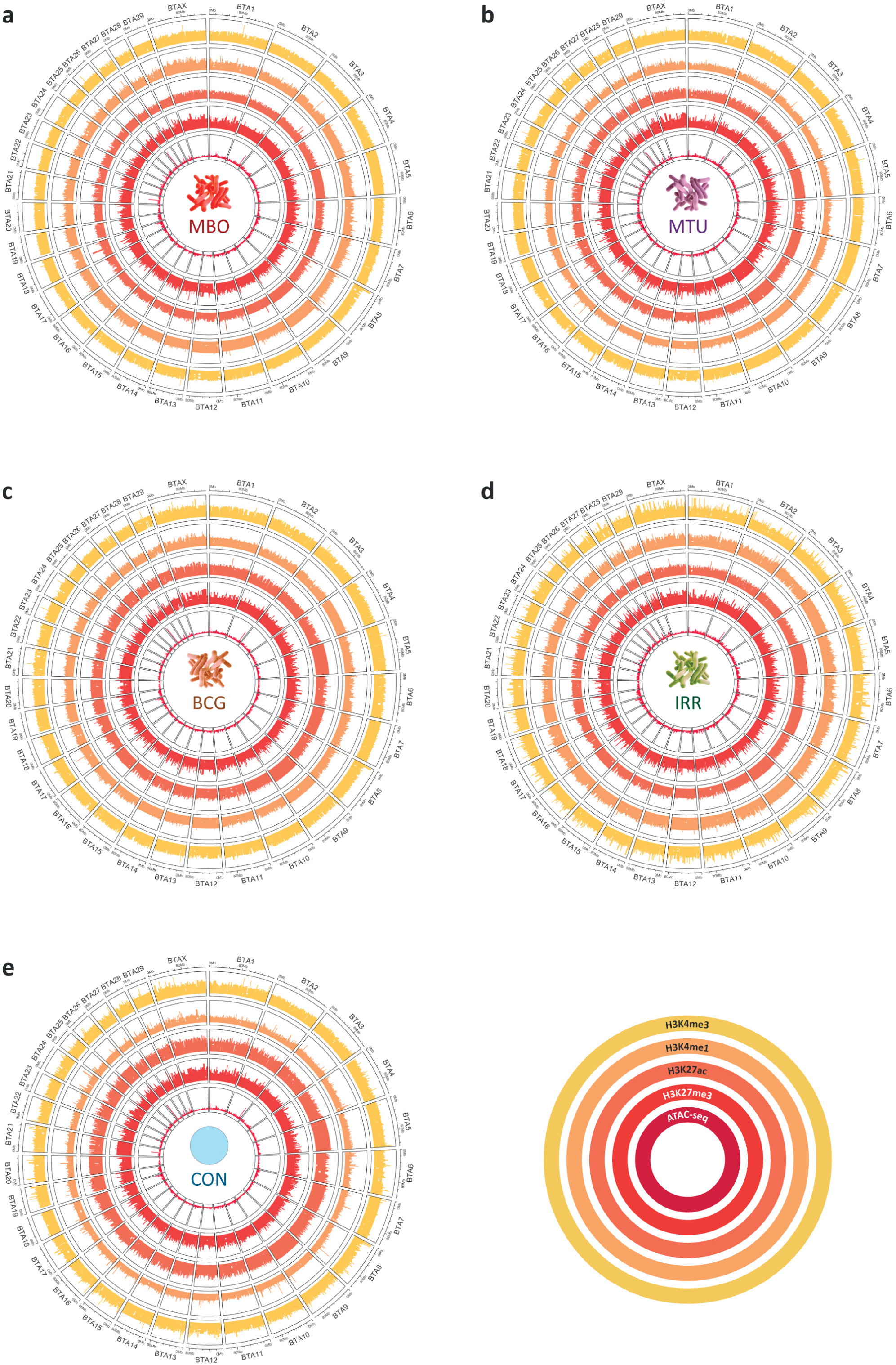
Circular plots of bAM challenge genome-wide peak distributions for ChIP-seq histone modification (H3K4me3, H3K4me3, H3K27ac, and H3K27me3) and ATAC-seq data. The bottom-right radial key shows the concentric circular plots for each epigenomic data type. **a** *Mycobacterium bovis* (MBO). **b** *M. tuberculosis* (MTU). **c** *M. bovis* BCG (BCG). **d** Gamma-irradiated (killed) *M. bovis* (IRR). **e** Control non-challenged (CON).

To identify epigenomic differences for the four contrasts between the MTBC challenged bAM groups (MBO, MTU, BCG, and IRR) and the control group (CON), differential affinity binding site (DABS) and differential open chromatin region (DOCR) results were collated from the ChIP-seq (Methods; Additional file 4: Tables S15‒S18) and ATAC-seq datasets (Additional file 5: Tables S21‒S22), respectively. To obtain a broader perspective on open chromatin regions that differed between MTBC-challenged bAM and control (CON) bAM samples, we focused first on the results from the ATAC-seq analysis, which revealed that the *M. bovis* (MBO/CON) and *M. tuberculosis* (MTU/CON) contrasts were the only two of the four primary challenge contrasts (MBO/CON, MTU/CON, BCG/CON, and IRR/CON) to exhibit statistically significant DOCRs: the MBO/CON contrast generated 226 DOCRs and the MTU/CON contrast produced 1 DOCR; however, there were no DOCRs observed for the BCG/CON and IRR/CON contrasts (Additional file 5: Tables S21‒S22; Additional file 1: Fig. S3a ‒d). The single statistically significant DOCR observed for the MTU/CON is located at the *TMPRSS2* gene, which encodes a transmembrane serine protease that is upregulated in human macrophages infected with *M. tuberculosis*. This DOCR was also found to be significant in the MBO/CON contrast. This *TMPRSS2* upregulation was also shown to contribute to an increased risk of SARS-CoV-2 infection, because TMPRSS2 is a key protease required for spike protein cleavage and ACE2 engagement by the virus (Sheerin et al. 2022). Examination of the transcriptomics results for *TMPRSS2* in the bAM challenged with four types of MTBC bacteria shows that it is substantially upregulated in the MBO/CON and MTU/CON contrasts with log_2_FC values of +4.47 (FDR-*P*_adj._ = 2.43 × 10^-43^) and +2.57 (FDR-*P*_adj._ = 9.48 × 10^-16^), respectively (Additional file 3: Tables S5‒S6). The *TMPRSS2* gene is also moderately upregulated in the BCG/CON contrast (log_2_FC = 1.43; FDR-*P*_adj._ = 1.48 × 10^-4^) but is not DE in the IRR/CON contrast (Additional file 3: Tables S7‒S8).

To quantify and compare the numbers of DOCRs for all 10 possible experimental contrasts, we determined the numbers of statistically significant DOCRs for the remaining six contrasts (MBO/MTU, BCG/MTU, IRR/MTU, MBO/BCG, IRR/BCG, and MBO/IRR) (Methods; Additional file 1: Fig. S3e). This demonstrated that, across all ATAC-seq challenge datasets, *M. bovis* bAM infection was the only challenge with relatively large numbers of DOCRs observed in all MBO contrasts with the control (CON) and other MTBC challenge groups (MTU, BCG, and IRR). In addition to the 226 DOCRs observed for the MBO/CON contrast, there were eight DOCRs observed for the MBO/MTU contrast. Also, in addition to the single statistically significant DOCR observed for the MTU/CON contrast, there was also one DOCR observed for the IRR/MTU contrast; 72 DOCRs observed for the MBO/BCG contrast, and 264 DOCRs observed for the MBO/IRR contrast. The other four contrasts (BCG/CON, IRR/CON, BCG/MTU, and IRR/BCG) did not reveal any statistically significant DOCRs (Additional file 1: Fig. S3e).

In addition to identifying statistically significant DOCRs in the ATAC-seq data, the epigenomic analyses revealed significant differences in ChIP-seq peak counts for histone modifications across the bAM challenge groups (MBO, MTU, BCG, and IRR) compared with the control (CON) group. Specifically, statistically significant differential affinity binding sites (DABSs) for H3K4me1, H3K4me3, and H3K27ac were observed at multiple genomic locations across the four primary challenge contrasts (MBO/CON, MTU/CON, BCG/CON, and IRR/CON) (Additional file 4: Tables S15‒S18). However, no significant H3K27me3 DABS were observed across any of the four primary MTBC/control contrasts. As observed in the ATAC-seq results, the majority of ChIP-seq-derived DABSs were detected in the *M. bovis*-infected samples compared with the control bAM (MBO/CON). Using the default criterion for DABS detection with DiffBind (FDR-*P*_adj._ < 0.05), 878 DABSs were detected for the MBO/CON contrast, and 56, 15, and 138 DABSs were detected for the MTU/CON, BCG/CON, and IRR/CON contrasts, respectively (Additional file 4: Tables S15‒S18). To illustrate this further, the DABs for H3K27ac, which marks higher activation of transcription (active enhancers), were plotted for each of the four primary challenge contrasts (MBO/CON, MTU/CON, BCG/CON, and IRR/CON) Fig. 5a‒d). In a similar fashion to the transcriptomics (RNA-seq) and open chromatin (ATAC-seq) results, differences in the ChIP-seq chromatin accessibility landscape were substantially more pronounced in the *M. bovis*-infected relative to the control bAM (MBO/CON). This is also evident when the transcriptomics results for genes encoding proteins involved in histone methylation and acetylation are examined in isolation. For these functional classes of genes, there are substantially more DEGs detected in the MBO/CON contrast compared to the other three primary contrasts (MTU/CON, BCG/CON, and IRR/CON) (Fig. 6).

**Fig. 5.**
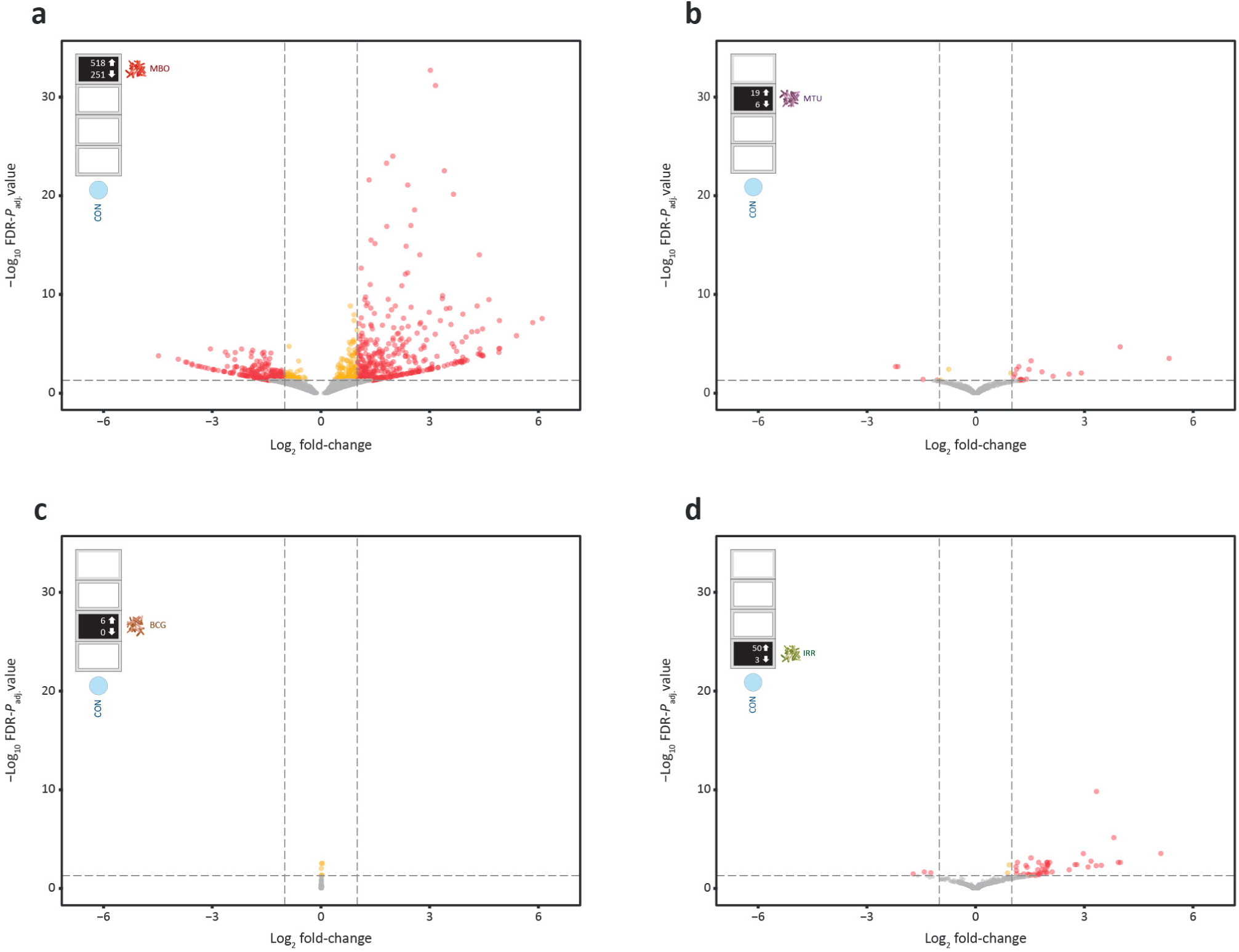
Volcano plots showing ChIP-seq H3K27ac differential affinity binding site (DABS) results for the four primary MTBC bAM challenges. The horizontal dotted line indicates the statistical significance threshold (FDR-*P*_adj._ < 0.05). Orange and red data points indicate significant DAB results (MTBC challenge vs. CON) with |log_2_FC values| < 1 and ≥ 1, respectively. **a** *Mycobacterium bovis* vs. control (MBO/CON). **b** *M. tuberculosis* vs. control (MTU/CON). **c** *M. bovis* BCG vs. control (BCG/CON). **d** gamma-irradiated (killed) *M. bovis* vs. control (IRR/CON).

**Fig. 6.**
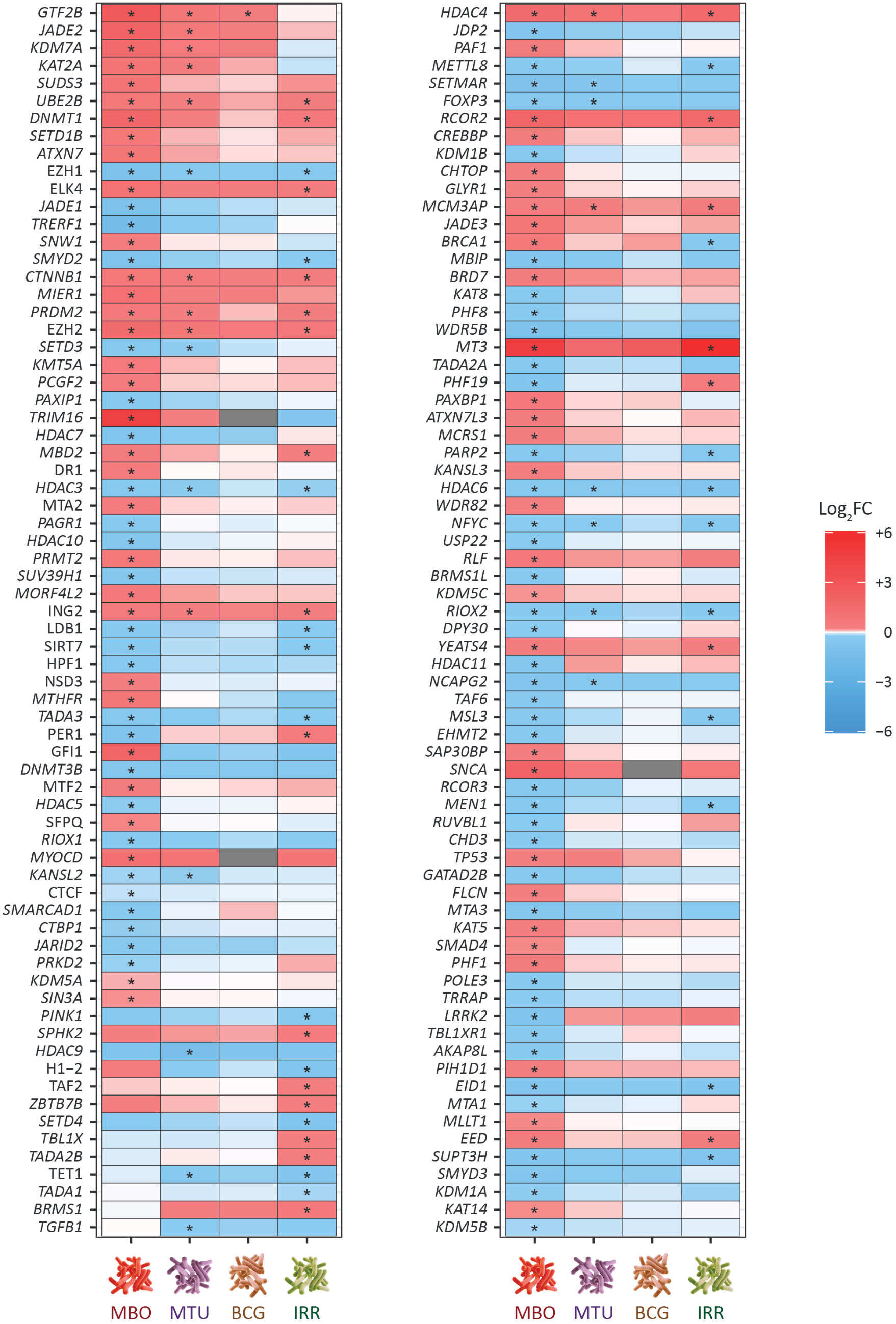
Gene expression results for genes involved in the processes of histone methylation and acetylation. Differentially expressed genes that are statistically significant (FDR-*P*_adj._ < 0.05) are indicated with an asterisk (*), and the heatmap shows log_2_FC values. The four primary bAM challenge contrasts are shown: *Mycobacterium bovis* vs control (MBO); *M. tuberculosis* vs control (MTU); *M. bovis* BCG vs control (BCG); and gamma-irradiated (killed) *M. bovis* vs control (IRR).

When the DOCR and DABS results are compiled and compared across the four primary challenge contrasts (MBO/CON, MTU/CON, BCG/CON, and IRR/CON), a clear pattern of increased chromatin accessibility is observed for bAM challenged with the four different types of MTBC (Table 1). The overall trends for the challenged bAM were increased deposition of the histone marks for proximal gene transcription (H3K4me3) and active enhancers (H3K27ac), coupled with decreased deposition of H3K4me1, which is consistent with previous observations indicating a counterintuitive repressive function for H3K4me1 in macrophages (Cheng et al. 2014). The results for bAM infected with *M. bovis* (MBO/CON) were particularly pronounced, with substantially more open chromatin regions (226 DOCRs) and genome-wide changes to chromatin accessibility determined by histone modifications (878 DABSs).

**Table 1:** Statistically significant differential chromatin modifications for the four MTBC challenge groups versus the control non-challenged group.

| bAM<br>challen<br>ge<br>contras<br>t | ATAC-seq: DOCRs |  | ChIP-seq: H3K4me3<br>DABSs |  | ChIP-seq: H3K4me1<br>DABSs |  | ChIP-seq: H3K27ac<br>DABS |  |
| --- | --- | --- | --- | --- | --- | --- | --- | --- |
|  | Increase<br>d | Decreas<br>ed | Increase<br>d | Decreas<br>ed | Increase<br>d | Decreas<br>ed | Increase<br>d | Decreas<br>ed |
| MBO<br>vs. CON | 226 | 0 | 61 | 9 | 0 | 39 | 518 | 251 |
| MTU vs.<br>CON | 1 | 0 | 0 | 0 | 1 | 30 | 19 | 6 |
| BCG vs.<br>CON | 0 | 0 | 0 | 0 | 0 | 9 | 6 | 0 |
| IRR vs.<br>CON | 0 | 0 | 11 | 0 | 2 | 72 | 50 | 3 |

### Integration of transcriptome and epigenome data reveals profound differences between bovine alveolar macrophages challenged with *Mycobacterium bovis* and other MTBC mycobacteria

The HOMER software suite v.4.11 was used to annotate genes within the differential ATAC-seq and ChIP-seq regions and sites (DOCRs and DABSs, respectively) detected for the four primary bAM challenge contrasts (MBO/CON, MTU/CON, BCG/CON, and IRR/CON). This annotation procedure facilitated the cataloguing of genes across the four groups that are located within the DABS and DOCR regions. For example, it was possible, therefore, to determine if the TSS of a particular gene underwent a differential histone modification caused by a specific MTBC bAM challenge (relative to the control non-challenged bAM) contrast. It was then straightforward to cross-reference the other contrasts to determine if the TSS chromatin environment for that gene was modified by the other MTBC bAM challenges. We could also establish if the histone changes were accompanied by additional DABSs/DOCRs. To visually illustrate the correlation between RNA-seq and ChIP-seq read deposition, we generated heatmaps showing the density of RNA-seq reads at each ChIP-seq histone mark (Additional file 1: Fig. S4).

Using this workflow, we obtained a consolidated set of exactly 1000 genes that had at least one DABS and/or DOCR in at least one of the four primary bAM challenge contrasts (MBO/CON, MTU/CON, BCG/CON, and IRR/CON). In this regard, there were 76 genes that had multiple associated DABSs and/or DOCRs, i.e. there were multiple differential regions of hypo/hyper H3 methylation or acetylation across the sequence of a gene—for example, *STAT1* and most notably *MX2* and *RSAD2* (Additional file 6: Tables S23). In total, there were 928 genes with at least one DABS and/or DOCR identified for the *M. bovis* infected compared to the control bAM (MBO/CON), and 54, 14, and 130 genes with at least one DABS and/or DOCR detected for the MTU/CON, BCG/CON, and IRR/CON bAM challenge contrasts, respectively (Additional file 1: Fig. S5).

A substantial majority of the genes associated with DABSs/DOCRs that were specific to one MTBC challenge were identified in the *M. bovis*-infected bAM (MBO/CON: 835, 83.5%). This was followed by the gamma-irradiated *M. bovis* challenged cells (IRR/CON: 67, 6.7%), the *M. tuberculosis* infected cells (MTU/CON: 9, 0.9%) and the *M. bovis* BCG infected cells (BCG/CON: 1, 0.1%) (Additional file 1: Fig. S5). Of the 1000 genes located at DABSs/DOCRs, only four were common to all four bAM challenge groups; two of which, *CD274* and *TNPO3,* were upregulated in the MTBC-challenged cells. The *TNPO3* gene encodes a protein (transportin 3) that acts as a nuclear import receptor for the trafficking of RNA splicing factor proteins into the cell nucleus (Christie et al. 2016). It has also been shown to have an important function in HIV-1 infection of host cells (Maertens et al. 2014); however, there is no evidence yet of a role in host macrophage responses to intracellular MTBC pathogens. The *CD274* gene, on the other hand, encodes the CD274 ligand, which has an immunoregulatory checkpoint function in restraining T cell activation (Sharpe and Pauken 2018). Increased expression of *CD274* may therefore be advantageous for intracellular MTBC pathogens (Shen et al. 2016; Hu et al. 2020).

We next investigated, at a broader level, the biological pathways and processes functionally associated with the consolidated multi-omic DAB/DOCR 1000-gene set. There were 20 statistically significant overrepresented IPA^®^ canonical pathways (Additional file 1: Fig. S6a). Overrepresented pathways for these 1000 genes included the *Inflammasome*, *Role of RIG1-like receptors in antiviral innate immunity*, *Activation of IRF by cytosolic pattern recognition receptors*, and *Death receptor signalling* pathways. However, the most significantly overrepresented pathway was *Interferon signalling*. The gene set overrepresentation analysis performed using g:Profiler yielded 194 statistically significant (FDR-*P*_adj._ < 0.05) functional entities (Additional file 1: Fig. S6b). The set of 20 top-ranked functional entities (by FDR-*P*_adj._) included GO:CC *Intracellular membrane-bound organelle*, KEGG *C-type lectin receptor signalling pathway*, KEGG *NOD-like receptor signalling pathway*, and the WikiPathways *IL-5 signalling pathway*.

Following this, the RNA-seq data were integrated to systematically determine which genes in the multi-omic DAB/DOCR 1000-gene set were significantly differentially expressed in the four primary bAM challenge contrasts (MBO/CON, MTU/CON, BCG/CON, and IRR/CON). IGV tracks for four selected immune genes (*BCL2*, *CD274*, *OAS2*, and *TMPRSS2*) are shown, illustrating the integration of RNA-seq, ChIP-seq, and ATAC-seq data (Additional file 1: Fig. S7a–d). In addition, the relationships among DABSs and DOCRs with DEGs illustrate that deposition of H3K4me3, H3K27ac, and DOCR locations are associated with an increase in gene expression, while, on the other hand, H3K4me1 marks a decrease in expression (Additional file 1: Fig. S8a–l). To achieve a holistic interpretable overview of the epigenomic regulatory relationships among differential histone modifications (DABSs), open chromatin regions (DOCRs), and transcriptional perturbations (DEGs) for the four primary MTBC-bAM challenge contrasts, the relevant data (Additional file 6: Table S23) were visualised as interaction networks using Cytoscape for the MBO/CON, MTU/CON, BCG/CON, and IRR/CON contrasts (Fig. 7a–d). The interaction network for the MBO/CON contrast is also shown in more detail (Additional file 1: Fig. S9 a–d).

**Fig. 7.**
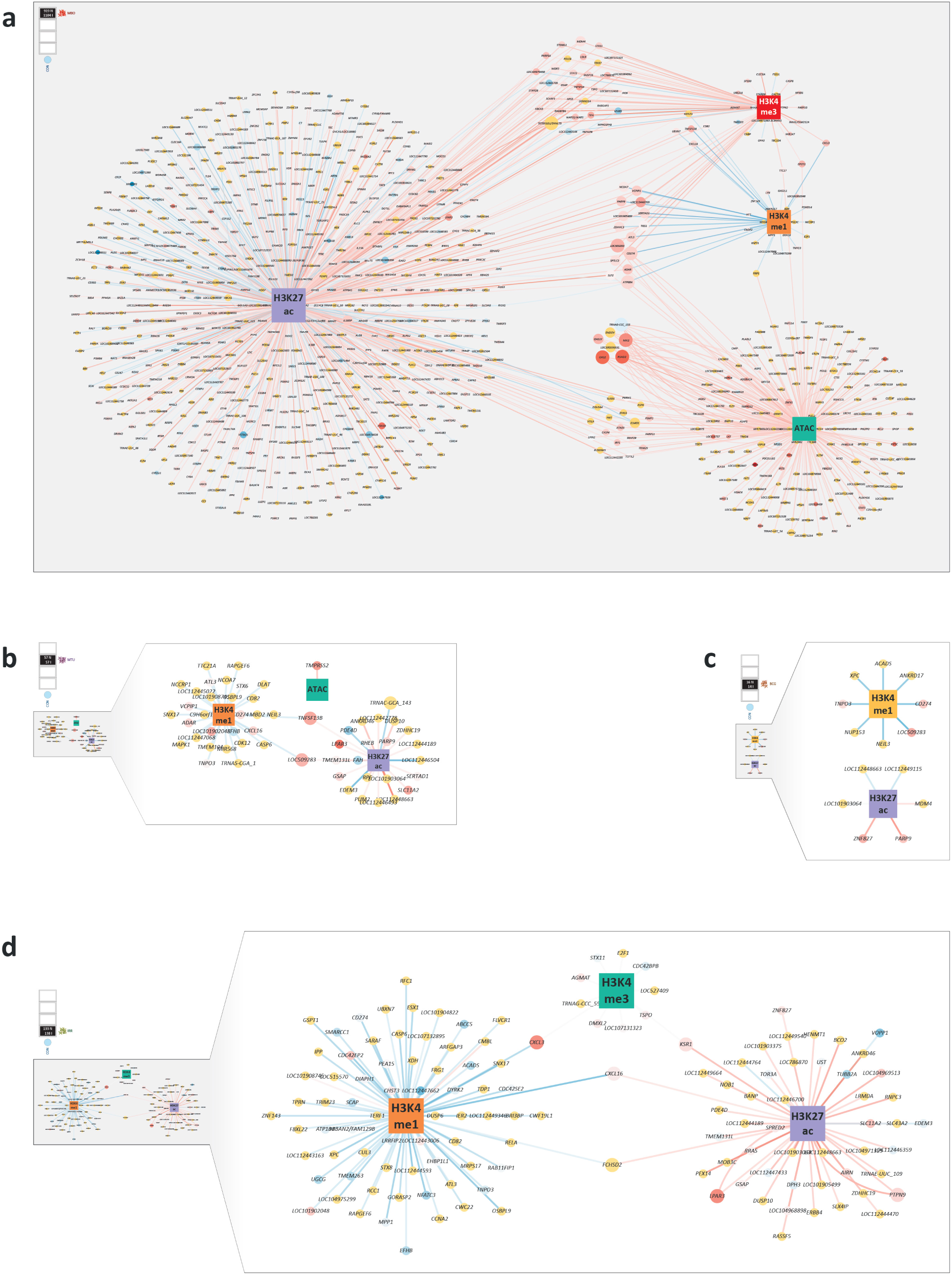
Network representations of the regulatory relationships among genes, histone modifications, and chromatin accessibility for the four primary MTBC bAM challenges. Each circular node in each network corresponds to a gene observed to be co-located with a differential histone modification or chromatin accessibility difference (represented as square anchor hub nodes). The circular node colour shows if a gene was differentially expressed (upregulated: red; downregulated: blue; and yellow: not differentially expressed). The lines (edges) between nodes show if a histone modification difference/chromatin accessibility change was larger (determined by log_2_FC for peak sizes) in the challenged bAM (red) or in the control non-challenged bAM (blue). Circular gene node size is determined by the number of connecting edges (degree). The numbers shown in each of the four keys represent the numbers of nodes (N) and interactions (I). **a** *M. bovis* vs control (MBO/CON). **b** *M. tuberculosis* vs control (MTU/CON). **c** *M. bovis* BCG vs control (BCG/CON). **d** gamma-irradiated (killed) *M. bovis* vs control (IRR/CON).

In the interaction network generated for the MBO/CON contrast, there are 919 nodes (including four anchor hub interaction nodes for H3K4me3, H3K4me1, H3K27ac and ATAC) and 1104 interactions, indicating that there are multiple genes with more than one histone modification (Fig. 7a). Conversely, in the interaction network generated for the MTU/CON contrast, there are only 57 nodes and 57 interactions (Fig. 7b). We also only obtained 16 nodes with 14 interactions for the BCG/CON contrast interaction network (Fig. 7c). The network complexity increased in the interaction network generated for the IRR/CON contrast with 133 nodes and 138 interactions (Fig 7d). Importantly, circular gene nodes can appear in multiple networks because some genes have an associated DABS/DOCR in more than one bAM challenge group (e.g., *CD274*). Taken together, these results highlight substantial epigenomic reconfiguration and transcriptional reprogramming evident in the *M. bovis*-challenged bAM compared with the other MTBC challenges. However, it is also important to note that among the other three bAM challenges, the gamma-irradiated *M. bovis* challenge elicits the most epigenomic reconfiguration and transcriptional perturbation (Fig. 2b, Fig. 3d, and Fig. 7d).

### Integration of the multi-omic DAB/DOCR 1000-gene set with a cattle GWAS data set for an *M. bovis* infection susceptibility trait

The consolidated multi-omic DAB/DOCR 1000-gene set was divided into eight gene subsets, which corresponded, firstly, to all DABS/DOCR genes for each primary bAM challenge contrast, including overlaps across other contrasts (four subsets: 928 MBO/CON overlap genes, 54 MTU/CON overlap genes, 14 BCG/CON overlap genes, and 130 IRR/CON overlap genes), and, secondly, all DABS/DOCR genes specific to a primary bAM challenge contrast, not including overlaps (four subsets: 835 MBO/CON-specific genes, 9 MTU/CON-specific genes, 1 BCG/CON-specific gene, and 67 IRR/CON-specific genes). Using the gwinteR tool, these eight gene subsets were integrated with a Holstein-Friesian breed GWAS data set for an *M. bovis* infection susceptibility trait (Ring et al. 2019). After this workflow was completed, we observed that only the largest MBO/CON overlap gene subset enriched additional statistically significant SNP variants associated with the *M. bovis* infection susceptibility trait (Additional file 6: Table S26). These corresponded to 64 SNPs within, or proximal to, four genes (*ERBB4*, *LRCH1*, *MRTFA*, and *RNPC3*) that had not been previously identified through integrative analyses of this GWAS dataset using either MTBC-challenged bAM or bTB peripheral blood functional genomics datasets (Hall et al. 2019; Hall et al. 2021; Hall et al. 2024; O’Grady et al. 2025). Circular Manhattan plots were generated to show the results of GWAS integration of the MBO/CON overlap gene subset (928 genes) for the *M. bovis* infection susceptibility trait (Fig. 8).

**Fig. 8.**
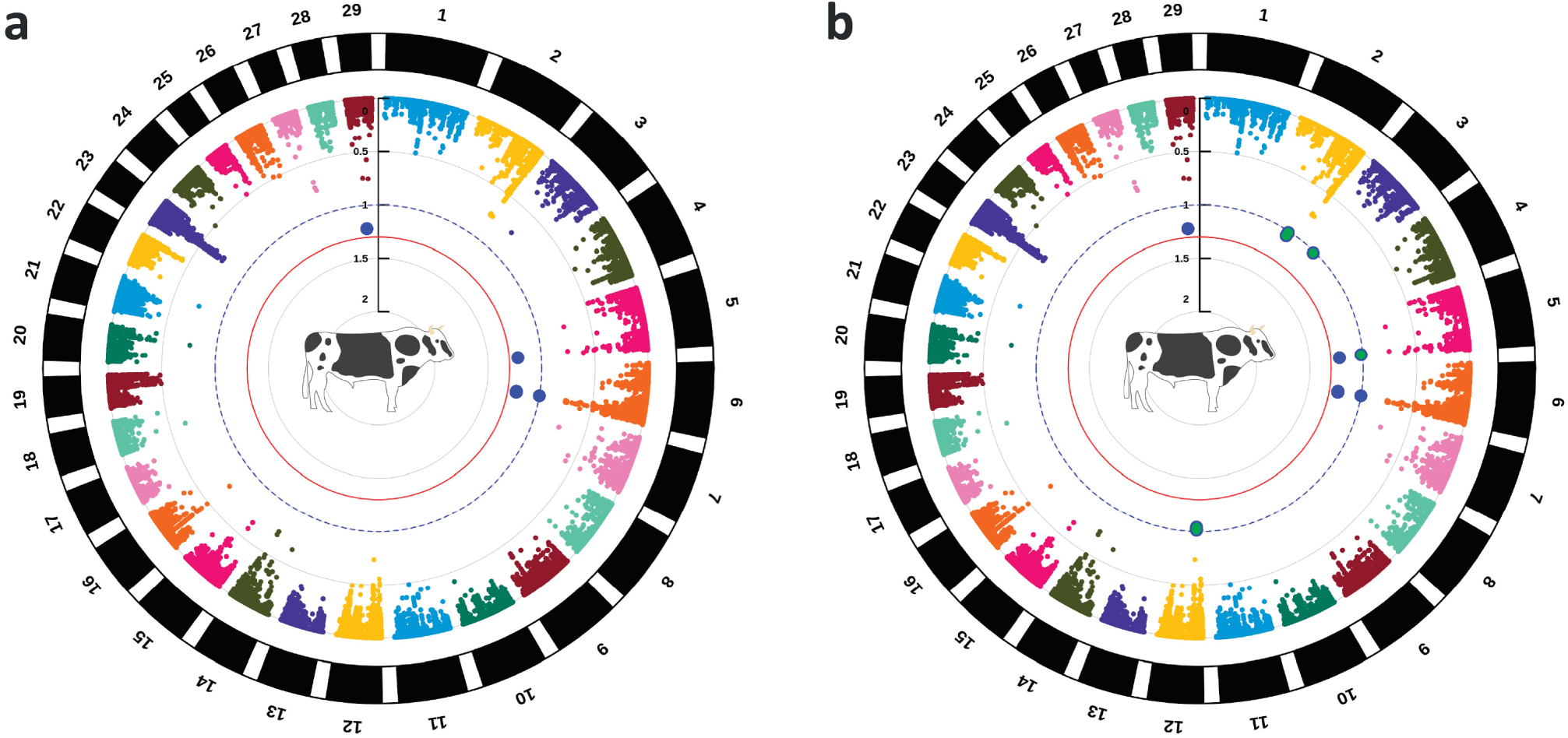
Integration of the MBO/CON overlap gene subset (928 genes) with a Holstein-Friesian *M. bovis* infection susceptibility trait GWAS dataset. **a** Circular Manhattan plot showing GWAS results before integration of the MBO/CON overlap gene subset. **b** Circular Manhattan plot showing GWAS results after integration of the MBO/CON overlap gene subset with the gwinteR tool. Large blue and green data points indicate binned statistically significant SNP clusters (FDR-*P*_adj._ < 0.10) prior to, and post-integration, respectively.

## Discussion

In this study, we applied a comprehensive multi-omics strategy, including RNA-seq, ChIP-seq for multiple histone marks, and ATAC-seq, to characterise transcriptional and epigenomic responses of bovine alveolar macrophages (bAM) challenged with intracellular mycobacteria. To do this, we compared bAM responses to four distinct types of MTBC: bovine-adapted *M. bovis* (MBO); *M. tuberculosis* (MTU), the primary cause of hTB; the attenuated *M. bovis* BCG vaccine strain (BCG); and dead, gamma-irradiated *M. bovis* cells (IRR). From these experiments, we could delineate how host adaptation and bacterial viability influence the reprogramming of alveolar macrophages—phagocytic cells that normally play a crucial role in the immune defence of the respiratory system—to favour the survival and dissemination of intracellular MTBC pathogens that cause tuberculosis in mammals. In addition, we integrated information on key genes identified from our multi-omics analyses with GWAS data to improve our understanding of the genomic architecture of an *M. bovis* infection susceptibility trait in Holstein-Friesian dairy cattle.

### Live, attenuated, and killed MTBC strains elicit distinct transcriptional responses in bovine alveolar macrophages

This study significantly extends previous work by our group showing that *M. bovis* and *M. tuberculosis* separately induce extensive rewiring of the bAM transcriptome, revealing both a core gene expression program induced by the two infections, and importantly, a distinct *M. bovis*-specific bAM transcriptional module governing the interaction between this MTBC strain and the bovine innate immune system (Malone et al. 2018). In the work presented here, using the same FDR-*P*_adj._ threshold of 0.05 but with no fold-change cut-off, we observed substantially larger numbers of DEGs at 24 hpi compared to this previous study, and an almost four-fold difference in the numbers of DEGs detected for the MBO/CON compared to the MTU/CON bAM challenge contrast (8,852 versus 2,312), which is also reflected in the number of DEGs detected for the direct MBO/MTU contrast (5,052) (Fig. 2b). The marked reprogramming of the virulent MBO-challenged bAM is even more striking in comparison to the attenuated *M. bovis* BCG strain. The BCG/CON bAM challenge contrast at 24 hpi only generated 586 DEGs—less than 7% of the total number of DEGs observed under the same conditions for the virulent MBO strain; a difference that is also apparent in the 6,741 DEGs detected in the direct MBO/BCG contrast (Fig. 2b). When considering these results, it is important to keep in mind that there were no statistically significant differences observed in the bacterial uptake by bAM for the MBO, MTU, and BCG challenges (Fig. S1f). It was somewhat surprising, therefore, that the gamma-irradiated (killed) *M. bovis* (IRR), which were phagocytosed two-fold less compared to the live bacterial strains, caused substantially more perturbation of the bAM transcriptome compared to the MTU/CON (3,320 vs. 2,312 DEGs) and BCG/CON (3,320 vs. 586 DEGs) bAM challenge contrasts (Fig. 2b). Given that γ-irradiation preserves bacterial integrity and PAMPs, it was expected that the initial infection response might be similar between live and dead bacteria. Indeed, Andreu and colleagues (Andreu et al. 2017) previously compared the transcriptional response of murine macrophages to gamma-irradiated vs live *M. tuberculosis*. They found that live and irradiated bacilli triggered a similar transcriptional response at 4 and 24 hours of infection, although the response in cells infected with irradiated bacilli diminished after 24 hours. Our findings of a more robust macrophage transcriptional response to *M. bovis*-irradiated cells compared to *M. tuberculosis* or BCG may reflect the inability of dead bacilli to actively suppress the bovine macrophage response to infection, a function that live MTBC are well adapted to.

Examination of the DEGs specific to the bAM challenged with *M. bovis* (the MBO/CON contrast), and the enriched pathways for the MBO/CON and MBO/IRR DEGs (Additional file 1: Fig. S2 and Additional file 3: Table S10), demonstrated that genes related to cytosolic sensing are more apparent in the *M. bovis*-infected bAM compared to the bAM challenged with the dead *M. bovis* cells. In addition, focusing on the results for the IPA^®^ *Role of RIG1-like receptors antiviral innate immunity* canonical pathway, activation of downstream cytosolic targets of the RIG-I-like receptor (RLR) pathway was evident for several bAM challenge contrasts involving the live MTBC strains (MBO/CON, MBO/IRR, MTU/CON, IRR/CON and BCG/CON) (Additional file 1: Fig. S10a–e). *M. bovis* infection induced coordinated perturbation of upstream RNA sensors and downstream signalling and transcriptional effectors (including IRF3/IRF7 and NF-κB), consistent with activation of a canonical type I interferon response. In contrast, *M. bovis* BCG and irradiated *M. bovis* triggered differential expression of selected RLR pathway components without comparable activation of downstream interferon regulatory networks, suggesting partial pathway engagement rather than a robust type I interferon programme (Additional file 1: Fig. S10a–e). The RIG-I–like receptor pathway mediates cytosolic RNA sensing and engages NF-κB and IRF7 signalling axes, coupling intracellular nucleic acid detection to inflammatory responses and, under appropriate conditions, induction of type I interferons (Fallahi-Sichani et al. 2012; Rehwinkel and Gack 2020; Wang et al. 2024).

The RIG-I-like receptor pathway for DEGs obtained from the killed *M. bovis* bAM challenge contrast (IRR/CON) shows activation of the NF-κb pathway but little or no activation of any of the other ligands, receptors, or targets, including IRF7 (Additional file 1: Fig S10e). Moreover, comparison of the RIG-I-like receptor pathway in the *M. bovis* versus gamma-irradiated (killed) *M. bovis* (MBO/IRR) and MBO/CON bAM challenge contrasts revealed highly similar pathway activation profiles (Additional file 1: Fig. S10a and Fig. S10b). This is to be expected as dead *M. bovis* lacks the ability to secrete ESX-1 effectors that facilitate phagosomal escape and induction of cytosolic nucleic acid-sensing pathways and downstream interferon-stimulated genes (ISGs), and also accounts for the differences in the top ten enriched IPA^®^ pathways observed for the MBO/CON and IRR/CON bAM challenges (Additional file 1: Fig. S2a and Fig. S2d). Additionally, RIG-I-like receptor pathway activation was seen in the MTU/CON contrast, but at lower levels of upregulation when compared to MBO/CON, suggesting less cytosolic escape of *M. tuberculosis* vs. *M. bovis* in bovine macrophages, as seen in our previous work (Malone et al. 2018).

Another feature of the transcriptomics results for the primary bAM challenge contrasts (MBO/CON, MTU/CON, BCG/CON, and IRR/CON) was the distinct expression patterns of Toll-like receptor (TLR) genes. The functions of TLRs in MTBC infections have been the subject of intensive research efforts for more than 25 years (Brightbill et al. 1999; Means et al. 1999; Underhill et al. 1999), and as pathogen recognition receptors (PRRs), TLRs play critical roles in MTBC infections (Mortaz et al. 2015; Stamm et al. 2015). Using various techniques to detect DEGs, it has long been established that the *TLR2* and *TLR4* genes are generally upregulated at 6–72 hpi across various human and bovine macrophage types in response to tuberculous mycobacterial infections. For example, using a targeted RNA-seq methodology (Ampliseq), Papp and colleagues (Papp et al. 2018) observed significantly increased *TLR2* expression in human AM (hAM) and monocyte-derived macrophages (hMDM) infected with *M. tuberculosis* at 24 hpi and 72 hpi, and detected significantly increased *TLR4* expression in hMDM at 72 hpi. Also, upstream regulator analysis of gene expression microarray data revealed upregulation of *TLR2* and *TLR4* in response to *M. tuberculosis* challenge in hAM and splenic macrophages (Lavalett et al. 2020a). Regarding bovine macrophages, we previously showed, using RT-qPCR, that *TLR2* and *TLR4* were upregulated in bovine MDM (bMDM) challenged with *M. bovis* 24 hpi (Taraktsoglou et al. 2011). Analysis of RNA-seq data from *M. bovis*-challenged bAM at 6, 24, and 48 hpi showed increased expression of *TLR2* and *TLR4* (Nalpas et al. 2015). Using the same bAM challenge model and RNA-seq, we also showed that *M. tuberculosis* challenge caused upregulation of these two genes (Malone et al. 2018). Finally, in a more recent integrative transcriptomics study of bAM, hAM, and hMDM, we showed that *TLR2* is upregulated 24 hpi after challenge with M. bovis and M. tuberculosis in bovine cells, and in both types of human macrophages 24 hpi with *M. tuberculosis* (Hall et al. 2024).

Surprisingly, therefore, in the present study although we observed that both *TLR2* and *TLR4* were upregulated at 24 hpi in the bAM challenged with live and dead *M. bovis* (MBO/CON and IRR/CON), there was no differential expression of these two genes in bAM challenged with *M. tuberculosis* or *M. bovis* BCG (MTU/CON and BCG/CON) (Additional file 3: Tables S5–S8). These differences are evident through downstream effects on other genes associated with innate immunity. For example, focusing on *TLR2*, signalling via TLR2 has been shown to drive the expression of the *CXCL5* gene, which is critical for innate cell communication and polymorphonuclear leukocyte-driven destructive inflammation during *M. tuberculosis* infection (Nouailles et al. 2014). As expected, *CXCL5* was significantly upregulated in the MBO/CON and IRR/CON bAM challenge contrasts with log_2_FC values of 2.40 (FDR-*P*_adj._ = 0.0077) and 4.65 (FDR-*P*_adj._ = 4.84 × 10^−8^), respectively; however, it did not exhibit any significant differential expression for the MTU/CON or BCG/CON bAM challenge contrasts. With regard to *TLR4*, signalling through TLR4 induces immune cell activation in response to live *M. tuberculosis* bacilli, phosphatidylinositol mannosides (PIM4–6) derived from *M. tuberculosis*, and purified cell wall preparations from *M. bovis* (Heldwein et al. 2003). In this regard, *TLR4*^-/-^ knockout mice show lower expression of *TNF* when exposed to *M. tuberculosis* (Reiling et al. 2002; Branger et al. 2004). In the present study, the *TNF* gene was upregulated across all the four primary bAM challenge contrasts; the highest increased expression was detected for the MBO/CON (log_2_FC = 3.98; FDR-*P*_adj._ = 3.99 × 10^−18^) and IRR/CON (log_2_FC = 2.73; FDR-*P*_adj._ = 3.99 × 10^−18^) contrasts compared to the MTU/CON (log_2_FC = 2.64; FDR-*P*_adj._ = 6.96 × 10^−9^) and BCG/CON (mean log_2_FC = 1.90; FDR-*P*_adj._ = 4.61 × 10^−4^) contrasts. Because human and murine TLR4 activate in the presence of *M. tuberculosis*, it is noteworthy that it remains largely inactivated in bAM unless stimulated by PAMPs derived from *M. bovis* cells (MBO and IRR).

One further difference in TLR activation among the primary bAM challenge contrasts should be highlighted: *TLR3* was upregulated only in *M. bovis*-challenged bAMs (MBO/CON) (Additional file 3: Tables S5–S8). Because TLR3 is an intracellular nucleic acid sensor (Sellge and Kufer 2015), significant activation of TLR3 solely in the MBO/CON bAM challenge contrast further supports a scenario where live *M. bovis* were the only bacilli to achieve substantial phagosomal escape. This suggests that the presence of *M. bovis* in the cytosol partly explains the markedly more extensive reprogramming observed in *M. bovis*-infected bAM compared with the other three MTBC challenges. However, it is important to note that *TLR3* expression may also be induced by host mitochondrial DNA released intracellularly following infection-induced cell damage (Carty et al. 2021). The causes and implications of these differences in TLR signalling activity between the host-adapted (*M. bovis*) and non-host-adapted (*M. tuberculosis*), attenuated (*M. bovis* BCG), and dead (gamma-irradiated *M. bovis*) bacilli remain to be fully elucidated; however, it is possible that TLR activation may contribute to the host tropism differences between *M. bovis* and *M. tuberculosis*.

Notwithstanding the substantial differences observed in the bAM transcriptome that are induced by challenge with different types of MTBC, it is important to highlight DEGs that had similar patterns of expression across the four primary bAM challenge contrasts (MBO/CON, MTU/CON, BCG/CON, and IRR/CON). Firstly, there were several chemokine genes that were significantly upregulated across all the contrasts (*CCL3, CCL4*, *CCL20*, *CXCL2*, *CXCL3*, and *CXCL16*), and secondly, numerous interleukin (*IL1A*, *IL1B*, *IL6*, *IL10*, *IL15*, and *IL27*) and interleukin receptor genes (*IL7R*, *IL10RB*, *IL17REL*, and *IL36G*) (Additional file 3: Tables S5–S8). Not only were these genes upregulated across all four primary bAM challenge contrasts, but many were also among the DEGs showing the greatest increases in expression relative to the control non-challenged bAM, highlighting the critical importance of cell-to-cell signalling during MTBC infection (Domingo-Gonzalez et al. 2016). In addition to chemokine and cytokine genes, other important genes that were upregulated across all bAM challenge contrasts included *NOS2,* which encodes nitric oxide synthase (NOS2), an antimicrobial effector produced by IFN-γ-activated macrophages (Bogdan 2015). The *NOS2* gene was markedly upregulated in the MBO/CON (log_2_FC = 5.84; FDR-*P*_adj._ = 1.32 × 10^−15^), MTU/CON (log_2_FC = 4.91; FDR-*P*_adj._ = 1.81 × 10^−11^), and BCG/CON (log_2_FC = 4.27; FDR-*P*_adj._ = 4.63 × 10^−8^) bAM challenge contrasts; however, it was most significantly upregulated for the IRR/CON contrast (log_2_FC = 6.50; FDR-*P*_adj._ = 3.37 × 10^−19^) (Additional file 3: Tables S5–S8). The strongest *NOS2* upregulation in IRR-challenged macrophages likely reflects unopposed TLR–NF-κB–driven transcription in the absence of immune-evasion mechanisms used by live MTBC strains to suppress antimicrobial effectors such as iNOS. Live bacilli actively dampen IFN-γ and STAT1–NF-κB signalling, whereas killed bacilli engage full proinflammatory transcriptional programs, yielding higher *NOS2* expression (Platanitis and Decker 2018; Chai et al. 2020).

The *SLAMF8* gene, which encodes a member of the CD2 family of cell surface proteins involved in lymphocyte activation, was also observed to be significantly upregulated across all four bAM challenge contrasts: MBO/CON (log_2_FC = 5.29; FDR-*P*_adj._ = 3.16 × 10^−14^), MTU/CON (log_2_FC = 4.35; FDR-*P*_adj._ = 6.28 × 10^−10^), BCG/CON (log_2_FC = 3.76; FDR-*P*_adj._ = 6.05 × 10^−7^), and IRR/CON (log_2_FC = 3.82; FDR-*P*_adj._ = 1.11 × 10^−7^), which we also previously observed at 24 and 48 hpi in independent bAM challenge experiments with *M. bovis* and *M. tuberculosis* (Malone et al. 2018; Hall et al. 2019). Finally, *CSF3,* which encodes the colony stimulating factor 3 protein (CSF3) a member of the IL-6 superfamily of cytokines was significantly upregulated across all four bAM challenge contrasts: MBO/CON (log_2_FC = 4.50; FDR-*P*_adj._ = 8.21 × 10^−4^), MTU/CON (log_2_FC = 4.05; FDR-*P*_adj._ = 6.52 × 10^−3^), BCG/CON (log_2_FC = 3.64; FDR-*P*_adj._ = 4.80 × 10^−2^), with the IRR/CON contrast exhibiting the most significant gene expression change (log_2_FC = 7.03; FDR-*P*_adj._ = 7.23 × 10^−8^).

### Epigenetic reprogramming in response to MTBC challenge in bovine alveolar macrophages is primarily restricted to *M. bovis* infection

During the last decade, it has been established that intracellular mycobacterial infections cause modifications to the host cell epigenome that can have major downstream impacts on host-pathogen interaction through reprogramming of the host transcriptome (Yaseen et al. 2015; Bouttier et al. 2016; Hall et al. 2019; Abo-Kadoum et al. 2021; Correa-Macedo et al. 2021; Lin et al. 2022; Madden et al. 2023)—findings that have been comprehensively reviewed (Gauba et al. 2021; Madden et al. 2022; Sengupta et al. 2023; Mishra et al. 2025). In the present study, where we conducted systematic analyses of the transcriptome and chromatin landscape using RNA-seq, ChIP-seq, and ATAC-seq, we observed that *M. bovis* challenge (MBO/CON) induced substantially more reconfiguration of the bAM epigenome compared to the other three challenges (MTU/CON, BCG/CON, and IRR/CON) (Table 1, Fig. 5a–d, and Additional file 1: Fig. S3a–d). This is also reflected in the transcriptomics results for genes encoding proteins involved with histone methylation and acetylation, processes that ultimately determine the accessibility of many genes to the host cell transcriptional machinery (Fig. 6). In this regard, our ChIP-seq results showed that the MBO/CON bAM challenge contrast generated 878 DABS, more than ten-fold the combined total (71) obtained for the other two live bacilli challenges (MTU/CON and BCG/CON). In addition, the bAM challenge with dead *M. bovis* cells (IRR/CON) produced almost twice as many DABS (138). The extent of chromatin remodelling for the MBO bAM challenge was also reflected in the open chromatin regions detected using ATAC-seq. For the MBO/CON, MBO/MTU, MBO/BCG, and MBO/IRR bAM challenge contrasts, we obtained 226, 8, 72, and 264 DOCR, respectively, which compares to a total of 2 DOCR detected for the other six contrasts (MTU/CON, MTU/BCG, MTU/IRR, BCG/CON, BCG/IRR, and IRR/CON) (Additional file 1: Fig S3e). Our results, therefore, support the hypothesis that epigenomic modifications are a major contributor to the substantial reprogramming of the bAM transcriptome induced by infection with virulent *M. bovis*.

One of the key questions arising from this multi-omics analysis is why *M. bovis* provokes such extensive chromatin remodelling in bAMs compared with *M. tuberculosis* or *M. bovis* BCG. The transcriptional data indicate stronger activation of innate immune signalling in the *M. bovis* group, yet this response is accompanied by distinctive shifts in histone modification and chromatin accessibility. To explore this, we examined differential affinity binding sites (DABs) and differential open chromatin regions (DOCRs) associated with genes central to Toll-like receptor (TLR) signalling—particularly the MyD88–TRAF6–NF-κB axis, which integrates early pathogen recognition with pro-inflammatory transcriptional responses.

Although identifying the precise cause of the observed DABs and DOCRs lies beyond the scope of this study, consistent patterns emerge. The absence of TLR activation in the *M. tuberculosis* (MTU) and *M. bovis* BCG (BCG) groups has downstream effects that are also evident at the epigenetic level. For example, reduced H3K4me1 deposition was detected at the promoter-proximal region of *MYD88*. In macrophages, H3K4me1 can act as a repressive chromatin mark (Hyun et al. 2017), and our data align with this observation—genes showing decreased H3K4me1 enrichment in infected cells were generally upregulated (Fig. 6). Following TLR engagement, MYD88 recruits IRAK1, which interacts with the E3 ubiquitin ligase TRAF6. TRAF6, together with the ubiquitin-conjugating enzyme complex UBC13–UEV1A, stimulates K63-linked polyubiquitination of both TRAF6 and the TAK1 kinase complex (Kawasaki and Kawai 2014). This, in turn, activates NF-κB and MAPK signalling cascades that coordinate pro-inflammatory responses to mycobacterial infection (Xu and Lei 2020). Accordingly, loss of the repressive H3K4me1 mark at *MYD88* following TLR activation is consistent with the chromatin remodelling required for pathway activation. Despite these epigenetic signatures, *MYD88* expression itself was not significantly altered (log₂FC = −0.33; *P*_adj_. = 0.052) and TAK1, encoded by *MAP3K7*, was only modestly upregulated (log₂FC = 0.25; *P*_adj_. = 0.032). This subdued transcriptional activation may reflect increased *TRAF2* expression, a negative regulator of TRAF6-dependent signalling (Kawasaki and Kawai 2014; Xie et al. 2019). The *TRAF2* gene was markedly upregulated in the MBO group and uniquely associated with increased H3K27ac, suggesting that *M. bovis* fine-tunes inflammatory signalling by epigenetically modulating inhibitory feedback loops within the TLR–MyD88 axis.

Finally, an H3K4me3 DAB was observed at the *NFKB1* transcription start site exclusively in the MB group, consistent with selective activation of the canonical NF-κB subunit. Together, these results indicate that *M. bovis* infection drives a unique interplay of activating (H3K4me3, H3K27ac) and repressive (H3K4me1) histone modifications at innate immune loci, reflecting an epigenetically mediated re-wiring of the TLR–MyD88–NF-κB pathway that distinguishes host-adapted *M. bovis* from non-host-adapted or attenuated MTBC strains.

Although there were several H3K4me3, H3K4me1, and H3K27ac histone modification DABS present in the MTU/CON, BCG/CON, and IRR/CON bAM challenge contrasts, significant DOCR results were almost exclusively observed in the MBO/CON primary bAM challenge contrast (Table 1). Examination of these regions highlighted genes involved in the innate immune response (Additional file 6: Table S23). For example, the *TRIM25, ISG15, ADAR*, and *DHX58* genes encode protein products involved with the RLR signalling pathway, which is involved with cytosolic sensing of RNA leading to activation of type I interferons and inflammatory cytokines, such as TNF (Rehwinkel and Gack 2020). Each of these genes was in a DOCR region, and all four were highly upregulated. As described above, based on RNA-seq results, the RLR signalling pathway was activated in bAM challenged with live bacilli (MBO, MTU, and BCG); however, activation was most marked in the MBO-challenged bAM. Localised patterns of open chromatin detected by ATAC-seq exclusively in the MBO contrast are consistent with these enhanced transcriptomic responses.

A cluster of related genes associated with multiple DOCR detected in the MBO-challenged bAM was the 2’-5’-oligoadenylate synthetase (OAS) gene family on BTA17 (*OAS1X*, *OAS1Y*, *OAS1Z*, *OAS2*, and *OASL*) (Additional file 1: Fig. S7c). The *OAS1Y* gene was significantly upregulated in the MBO/CON (log_2_FC = 3.53; FDR-*P*_adj._ = 3.97 × 10^−64^), MTU/CON (log_2_FC = 1.95; FDR-*P*_adj._ = 1.11 × 10^−20^). and BCG/CON (log_2_FC = 1.35; FDR-*P*_adj._ = 3.31 × 10^−9^) bAM challenge contrasts; it was not upregulated in the IRR/CON contrast (Additional file 6: Table S23). It is important to note that *OAS2* was another OAS family DEG we identified that was significantly upregulated in the MBO/CON (log_2_FC =3.57; FDR-*P*_adj._ = 6.47 × 10^−59^), MTU/CON (log_2_FC = 2.15; FDR-*P*_adj._ = 1.08 × 10^−19^) and BCG/CON (log_2_FC = 1.45; FDR-*P*_adj._ = 2.32 × 10^−8^) bAM challenge contrasts; it was not statistically significant for the IRR/CON contrast. The OAS gene family members are interferon-stimulated genes, and, using the human THP-1 macrophage-like cell line, Leisching et al. (2019) have shown that the protein products of the human orthologs (*OAS1* and *OAS2*) act to restrict intracellular *M. tuberculosis* replication and enhance cytokine secretion. Although our OAS DEG results are somewhat equivocal, the presence of the OAS gene cluster DOCR regions only in MBO-challenged bAMs may contribute to the increased relative expression of *OAS1X* in these cells.

Although most of the DABS and DOCR regions for the primary bAM challenge contrasts were detected in the *M. bovis*-infected cells (MBO/CON), there were also significant numbers of DABS regions identified in the IRR/CON bAM challenge contrast; in fact, the highest number of H3K4me1 DABS were detected for this contrast (74), almost twice the number detected for the MBO/CON contrast (39) (Table 1). It is important to note that, for this histone mark, almost all the DABS indicated decreased deposition in MTBC-challenged cells; for example, 39/39 in the MBO/CON and 72/74 in the IRR/CON bAM challenge contrasts. Again, it is also important to emphasise that H3K4me1—normally enriched at active and primed enhancers—has been hypothesised to be a repressive histone mark in macrophages (Cheng et al. 2014). Our results for bAM challenged with live/attenuated/dead MTBC clearly support this hypothesis (Table 1). The reduced H3K4me1 deposition we observed in bAM challenged with live and dead MTBC bacilli suggested that it may partly drive the upregulation of many innate immune and downstream genes (Additional file 6: Table S23). For example, the *CD82* locus had decreased H3K4me1 deposition in *M. bovis*-infected bAM compared to non-challenged control bAM (MBO/CON: log_2_FC −2.25; FDR-*P*_adj._ = 0.0402) and increased expression (log_2_FC 0.50; FDR-*P*_adj._ = 4.37 × 10^−4^). Increased expression of *CD82* has been shown to lead to retardation of phagosome maturation, alteration of the innate immune response, and enhanced survival of virulent *M. tuberculosis* in macrophages, and it was found to be highly expressed in granulomas from hTB patients (Koh et al. 2018). The *CD82* locus also showed decreased H3K4me1 deposition in the IRR/CON bAM contrast (log_2_FC = −3.00; FDR-*P*_adj._ = 0.0163) but was not significantly different in expression. In addition, the *CXCL3* locus also showed substantially decreased H3K4me1 deposition for the MBO/CON bAM contrast (log_2_FC = −4.25; FDR-*P*_adj._ = 7.69 × 10^−5^) and increased expression (log_2_FC = 2.93; FDR-*P*_adj._ = 6.47 × 10^−8^). Expression of the *CXCL3* gene has been shown to be elevated in peripheral blood mononuclear cells (PBMC) from active and latent hTB patients (Yu et al. 2017) and in PBMC from naïve individuals stimulated with *M. tuberculosis* H37Ra antigen (Mtb-Ag) (Wei et al. 2023). The *CXCL3* locus also had decreased H3K4me1 deposition in the IRR/CON bAM contrast (log_2_FC = −2.97; FDR-*P*_adj._ = 2.77 × 10^−4^) and was also substantially increased in expression (log_2_FC = 4.47; FDR-*P*_adj._ = 2.10 × 10^−17^). Finally, the *CD274* locus had markedly decreased H3K4me1 deposition in the MBO/CON bAM contrast (log_2_FC = −5.32; FDR-*P*_adj._ = 1.58 × 10^−4^) and increased expression (log_2_FC = 2.08; FDR-*P*_adj._ = 2.22 × 10^−79^). It also exhibited decreased H3K4me1 deposition and increased expression for the other three bAM challenge contrasts (MTU/CON, BCG/CON, and IRR/CON) (Additional file 6: Table S23). As described in the Results section, increased expression of *CD274* in alveolar macrophages may be advantageous for intracellular MTBC pathogens (Shen et al. 2016; Hu et al. 2020).

### Multi-omics integration focused on the alveolar macrophage response to MTBC challenge sheds new light on the function of key genes

Integration of transcriptomic (RNA-seq), histone modification (ChIP-seq), and open chromatin state (ATAC-seq) datasets has provided comprehensive insights into host-pathogen interactions in bAM challenged with live/attenuated/dead MTBC. A multi-omics interaction network representation of four epigenomic data types (H3K4me1, H3K4me3, H3K27ac, and ATAC open chromatin state), plus transcriptomics (RNA-seq), facilitates exploration and evaluation of bAM gene regulation in response to these MTB challenges (Fig. 7 and Additional file 1: Fig. S9). Systematic examination of these networks requires keeping in mind that H3K4me1 has a repressive function in macrophages (Cheng et al. 2014), that H3K4me3 indicates active transcription of proximal genes, H3K27ac is associated with active enhancers, and that DOCRs detected with ATAC-seq correspond to open chromatin (Zhang and Cao 2019).

The *CD274* gene exemplifies how multi-omics integration can provide new functional insights into infection biology, as shown in the interaction network for the MBO/CON bAM challenge contrast (Fig. 7a and Additional file 1: Fig. S9b). The integrative transducing effect of the active histone marks (increased H3K4me and 3H3K27ac), repressive histone marks (reduced H3K4me1), and open chromatin (increased ATAC signal) on *CD274* transcription (increased RNA-seq expression: log_2_FC = 2.08; FDR-*P*_adj._ = 2.22 × 10^−79^) is evident at this gene node. Epigenomic regulation of *CD274*, although not as dramatic, is also evident with decreased H3K4me1 and increased RNA-seq expression for the MTU/CON (log_2_FC = 0.88; FDR-*P*_adj._ = 4.36 × 10^−15^), BCG/CON (log_2_FC = 0.62; FDR-*P*_adj._ = 4.91 × 10^−7^), and IRR/CON (log_2_FC = 0.42; FDR-*P*_adj._ = 8.23 × 10^−4^) contrasts. These results suggest that epigenomic regulation of *CD274* activity may contribute to *M. bovis* virulence, immunoevasion, and host tropism. This gene encodes the CD274 ligand (also known as PDL1) of the inhibitory programmed cell death protein 1 (PDCD1—also known as PD1) receptor, which is expressed on T cells and has an immunoregulatory checkpoint function in restraining T cell activation (Sharpe and Pauken 2018). Interestingly, we only observed increased expression of *PDCD1* in the MBO/CON bAM challenge group (log_2_FC = 3.55; FDR-*P*_adj._ = 7.13 × 10^−5^); however, there were no DABS or DOCR associated with the *PDCD1* locus, and the gene was not significantly altered in expression for any of the other three primary bAM challenge contrasts (MTU/CON, BCG/CON, or IRR/CON).

Expression of *PDCD1* and *CD274* was previously observed to be significantly elevated in active hTB cases compared to healthy controls (Shen et al. 2016). This study also demonstrated that antibody-directed blockade of the PDCD1/CD274 (PD1/PDL1) pathway in human MDM/CD4^+^ T cell co-cultures inhibited *M. bovis* BCG replication in the hMDM. In addition, it has been shown that murine AM (mAM) infected with *M. tuberculosis* H37Rv and *M. bovis* BCG exhibited elevated *PDCD1* and *CD274* mRNA levels, concomitant with increased abundance of the PD1 and PDL1 proteins (Hu et al. 2020). Increased activation of the PD1/PDL1 pathway may therefore benefit pathogenic MTBC by impairing intracellular mycobacterial killing in macrophages and dampening T-cell activation. Our results support this hypothesis and the heightened activation of the pathway—which may be epigenetically driven—in bAM infected with virulent *M. bovis* indicates immunomodulation by the pathogen. Paradoxically, however, this relatively simple interaction is complicated by accumulating evidence *in vivo* that immune checkpoint inhibitor (ICI) therapy for cancer treatment, which blocks PD1 or PDL1, can increase susceptibility to *M. tuberculosis* infection and disease reactivation, albeit contingent on the immune status of individual patients (Vaddi et al. 2024). This picture is further complicated by recent work using spatial proteomics of granuloma tissue from hTB cases, which showed that increased PDL1 expression was associated with progression to active TB (McCaffrey et al. 2022). These observations and our bAM MTBC challenge results suggest that disentangling the role of the PD1/PDL1 pathway in human and animal MTBC infections will require coordinated *in vitro*, *ex vivo*, and *in vivo* research efforts that leverage all omics technologies that can be used to study genome regulation and host-pathogen interaction.

The *IRF1* gene is also located in a genomic region that had multiple chromatin modifications and exhibited differential expression in bAM infected with *M. bovis* (Fig. 7, Additional file 1: Fig. S9d). The *IRF1* gene encodes interferon regulatory factor 1 (IRF1), a transcriptional regulator of innate and acquired immunity (Feng et al. 2021). The chromatin alterations evident in the MBO/CON bAM challenge contrast (increased H3K27ac and ATAC signal) corresponded to upregulation of *IRF1* expression (log_2_FC = 2.42; FDR-*P*_adj._ = 1.90 × 10^−34^). The gene was also moderately upregulated in the other three MTBC bAM challenge contrasts (log_2_FC: 0.88–1.38; FDR-*P*_adj._: ranging between 8.96 × 10^−5^ and 5.74 × 10^−12^), but there was no *IRF1* chromatin reconfiguration observed for these contrasts (Additional file 3: Tables S5–S8 and Additional file 6: Table S23). It has been shown that homozygous loss-of-function mutations in *IRF1* produce a severe Mendelian susceptibility to mycobacterial disease (MSMD) in humans (Rosain et al. 2023). This study also convincingly demonstrated that IRF1 governs IFN-γ-dependent macrophage activation and resistance to intracellular pathogens. In addition, recent deployment of a bovine multicellular spheroid model of MTBC infection and follow-on multi-omics analyses highlighted the importance of *IRF1* expression as an early biomarker of MTBC infection (Bhaskar et al. 2025). It is noteworthy, therefore, that our results indicate that a bAM epigenetic program drives the upregulation of *IRF1*, which is exclusive to *M. bovis* infection.

In a similar fashion to *IRF1*, the *LDLR* gene exhibited evidence for epigenomic reconfiguration exclusive to the MBO/CON bAM challenge contrast (Fig. 7, Additional file 1: Fig. S9b). The H3K27ac and ATAC signal observed predicted upregulation of *LDLR* in these bAM, which we observed (log_2_FC = 3.00; FDR-*P*_adj._ = 5.62 × 10^−34^). Again, we also found modest upregulation of *LDLR* expression without chromatin modification in the other three MTBC bAM challenge contrasts (log_2_FC: 1.40–1.65; FDR-*P*_adj._: ranging between 2.00 × 10^−7^ and 5.19 × 10^−11^) (Additional file 3: Tables S5–S8 and Additional file 6: Table S23). The *LDLR* gene encodes the low-density lipoprotein receptor (LDR1) that mediates endocytosis of cholesterol-rich low-density lipoprotein (LDL) (Schmidt et al. 2025). The epigenetic regulation of *LDLR* expression we observe in *M. bovis*-infected bAM may therefore be relevant to recent observations that LDLR plays a key role in the *M. bovis* BCG-induced trained immunity in macrophages that promotes cholesterol uptake and leads to enhanced type I interferon responses (Lai et al. 2025).

The final part of this study was the integration of the consolidated multi-omics DAB/DOCR 1000-gene set with GWAS data from the Holstein-Friesian breed for an *M. bovis* infection susceptibility trait (Ring et al. 2019). The four genes identified via the GWAS integration included the *ERBB4* gene (BTA2: 56 SNPs) that encodes an epidermal growth factor receptor involved in innate immunity signalling, and that binds epiregulin encoded by the *EREG* gene, which has been shown to contain polymorphisms that influence several hTB susceptibility traits in human populations (Thuong et al. 2012; Cao et al. 2019). Also included was *RNPC3* (BTA3: 1 SNP), which encodes the RNA binding region (RNP1, RRM) containing 3 protein, a component of the spliceosome (Beusch and Madhani 2024). It is important to note that, in addition to the MBO-challenged bAM, *ERBB4* and *RNPC3* also exhibited DABS in the IRR-challenged bAM. The other two genes were *MRTFA* (previously known as *MKL1*; BTA5: 1 SNP), which encodes the myocardin related transcription factor A protein—a coactivator of serum response factor (SRF)—that exhibits a loss-of-function mutation in humans leading to immunodeficiency and severe susceptibility to bacterial infection (Record et al. 2015); and *LRCH1* (BTA12: 6 SNPs), which encodes the leucine rich repeats and calponin homology domain containing 1 protein that has been shown to be a negative regulator of LAT-mediated TCR signal transduction and CD8^+^ T cell responses against the intracellular bacterial pathogen *Listeria monocytogenes* (Liu et al. 2020). These results will be relevant to genome-enabled breeding programmes aimed at enhancing bTB disease resilience traits in domestic cattle (Banos 2023; Madenci et al. 2025).

## Conclusions

There are limitations inherent in this study that should be considered. For example, a parallel reciprocal experiment involving multi-omics analyses of cultured hAM challenged with the same MTBC types would provide valuable complementary information to deepen understanding of host tropism for tuberculous mycobacteria. In addition, single-cell multi-omics (i.e., scRNA-seq, scChIP-seq, and scATAC-seq) would be a superior approach for understanding how macrophage heterogeneity in the lung can impact MTBC infections (Russell et al. 2025). Notwithstanding these limitations, we have shown in this study that epigenetic reprogramming of the transcriptome differentiates the bovine alveolar macrophage response to bovine-adapted *M. bovis* from that elicited by closely related MTBC strains. We also demonstrate that multiple types of functional genomics data outputs can be integrated with relevant GWAS data and signpost future multi-omics research that can exploit the very large GWAS datasets available for cattle and other livestock species.

## Materials and Methods

### Harvesting bovine alveolar macrophages by lung lavage

Six unrelated and age-matched (6–8 weeks old) male Holstein-Friesian calves were used in this study (Additional file 2: Table S1). All animals were maintained under uniform housing conditions and nutritional regimens at the UCD Lyons Research Farm (Newcastle, County Kildare, Ireland). The subject animals were selected from a tuberculosis-free herd that is screened annually using the single intradermal comparative tuberculin test (SICTT). These cattle were also negative for infection with *Brucella abortus*, *M. avium* subsp. *paratuberculosis*, *Salmonella* Typhimurium, bovine herpesvirus 1 (BHV-1) and bovine viral diarrhoea (BVD) virus. All animal procedures were performed in accordance with Irish law (SI 543/2012, as per EU Directive 2010/63/EU), and ethical approval for the study was obtained from the University College Dublin (UCD) Animal Ethics Committee (protocol number AREC-19-07-Gordon).

Euthanasia was performed by administration into the jugular vein of an overdose of sodium pentobarbital. Lungs and heart were separated from the animal and were visually inspected by a veterinary surgeon for inflammation and pathology before proceeding to pulmonary lavage for harvesting of bovine alveolar macrophages (bAM) as previously described by our group (Magee et al. 2014; Nalpas et al. 2015). Lungs were separated from the heart to prevent blood contamination. The washes were performed using 5–6 L of phosphate-buffered saline (PBS) (Gibco, Thermo Fisher Scientific) infused into the lungs via the trachea (500 ml per infusion), followed by a massage of the infused lungs and then collection of the resulting cell suspension into sterile beakers. Cell suspension (50 ml) from the first infusion was collected, pelleted by centrifugation (200 × g for 10 min at RT), resuspended in 10 ml PBS, and cultured on a range of solid media to screen for microbial contamination; all animals tested negative.

The cell suspension was transferred to 50 ml conical tubes, filtered through 100 µm cell strainers, and centrifuged (300 × g for 10 min at RT). The supernatants were discarded, except for the 5 ml volumes covering the cell pellets, which were pooled. When required, red blood cell lysis was performed by centrifugation (300 × g for 10 min at RT), followed by removal of the supernatant, except for 10 ml remaining on top of the pellet. Then, 30 ml of erythrocyte lysis buffer (Qiagen) was added, the pellet was resuspended and incubated for 3 min, and then the tube was filled to 50 ml with PBS. The tubes were then centrifuged (300 × g for 10 min at RT), the supernatants were discarded entirely, and the pellets were resuspended in 5 ml R medium (RPMI medium with GlutaMAX (Gibco), supplemented with 10% foetal bovine serum (FBS) (Gibco)) and pooled into volumes of 40 ml. The cells were counted using a haemocytometer and 0.4% Trypan Blue Solution (Thermo Fisher Scientific) to estimate viability. The cells were then centrifuged (300 × g for 10 min at RT) and resuspended in freezing solution (10% DMSO (Sigma-Aldrich), 90% FBS) at a density of approximately 2.5 × 10^7^ cells/ml. Cell aliquots (1.5 ml) were then prepared in 2 ml sterile cryovials (Sarstedt) and placed into Mr Frosty^®^ Cryo −1°C/min freezing containers (Nalgene^®^, Thermo Fisher Scientific) containing 100% isopropanol. Cryovials were stored at −80°C for 20 h, after which they were removed from the freezing containers and transferred to liquid nitrogen until required for further use.

### Culture of bovine alveolar macrophages

Stored cells were thawed, transferred to a 50 ml tube with 20 ml complete R+ medium (RPMI medium with GlutaMAX (Gibco) supplemented with 10% FBS (Gibco), 2.5 µg/ml Amphotericin B (Sigma-Aldrich) and 50 µg/ml Cefotaxime (Sigma-Aldrich)) and centrifuged (200 × g for 5 min at RT). The supernatant was discarded, and cells were resuspended in 20 ml of fresh R+ medium. Cell solutions were then placed in T75 vented culture flasks (CELLSTAR^®^, Greiner Bio-One) and incubated at 37°C with 5% CO_2_ for 24 h prior to plating in 24-well plates. After incubation, the medium was removed together with non-adherent cells. Adherent bAM cells were washed with 15 ml PBS and dissociated by adding 10 ml pre-warmed non-enzymatic cell dissociation buffer in PBS (Gibco) to each culture flask, incubating for 10 min at RT, gently tapping the flask after 5 min, and then collecting cells by gentle scraping with a cell scraper. The bAM cells, in cell dissociation buffer, were transferred to a 50 ml tube with 20 ml R medium (RPMI medium with GlutaMAX (Gibco) supplemented with 10% Foetal calf serum (Gibco), no antibiotics) and pelleted (200 × g for 5 min at RT), then resuspended in 5 ml R medium. The number of viable cells was measured using 0.4% Trypan Blue Solution, and cells were counted using a haemocytometer. Cell counts were adjusted to 5 × 10^5^ cells/ml (based on viable cell counts) in R medium, and 5 × 10^5^ cells were seeded per well in individual 24-flat-well tissue culture plates (Sarstedt). A 10 mm-diameter round coverslip was placed at the bottom of the wells used for fluorescence microscopy before cell seeding. The plates were incubated for 24 h at 37°C in 5% CO_2_ until required for mycobacterial challenge.

### Bacterial culture and preparation for *in vitro* challenge assay

Culturing of *M. bovis* AF2122/97, *M. bovis* BCG Denmark, and *M. tuberculosis* H37Rv was performed in a Biosafety Containment Level 3 (BCL3) laboratory, which conformed to the national guidelines on the use of Hazard Group 3 infectious organisms. *M. bovis* BCG Denmark and *M. tuberculosis* H37Rv were cultured in Middlebrook 7H9 medium (Difco^™^, BD) with 0.05% Tween 80, 0.2% glycerol, 0.5% BSA, 0.2% glucose, and 0.085% NaCl. *M. bovis* AF2122/97 was cultured in the same media supplemented with 10 mM sodium pyruvate (Sigma-Aldrich). Gamma-irradiated whole *M. bovis* AF2122/97 cells were obtained from BEI Resources (Catalogue number NR-31210, ATCC). To prepare the gamma-irradiated cells for use and to minimise clumping, 1 ml of the original stock was added to 9 ml of Middlebrook 7H9 medium, sonicated at full power in a Branson 2510 ultrasonic cleaner (Branson Ultrasonics Corporation) for 15 min, and passaged eight times through a 26G needle. The bacterial suspension was aliquoted into 2 ml centrifuge tubes and centrifuged (100 × g for 3 min at RT). The bacterial suspensions were then collected in 15 ml tubes, and the pellets were discarded. The optical density at 600 nm (OD_600_ nm) of the bacterial suspension was measured and adjusted to 0.3–0.5. The bacterial suspension was then aliquoted into 1 ml tubes and stored at −80°C.

To prepare live bacterial strains for the infection experiments, 18 days prior to the bAM challenge, 1 ml of mycobacterial stock was retrieved from the −80°C freezer, thawed, and then added to 4 ml of medium. These starter cultures were incubated for 1 week at 37°C, after which 5 ml of fresh media was added and incubated for a further week without shaking. Cultures were then scaled up to a larger volume by transferring 4 ml of the starter culture into 46 ml of media in 2-L-capacity cell culture roller bottles (Greiner). The bottles were incubated at 37°C with rolling, at approximately 1-2 rotations per minute, for 4 days. The bacterial cultures were monitored until they reached the required optical density of 0.5–0.6 at OD_600_ nm. On the day of bAM challenge, 15 ml aliquots were removed and transferred to 50 ml conical tubes. The cultures were centrifuged (1000 × g for 10 min at RT). Supernatants were discarded, and 10–15 glass beads (3 mm) were added to the tubes, which were then vortexed for 1 min. Complete R medium (6 ml) was then added to resuspend the pellet, and tubes were left to rest for 5 min. The top 5 ml was then removed and transferred to fresh 50 ml conical tubes, which were centrifuged (300 × g for 7 min at RT). The supernatant (4 ml) was then passaged 15 times through a 26G needle. Ready-to-use aliquots of gamma-irradiated whole *M. bovis* AF2122/97 were retrieved from the −80°C freezer and thawed, centrifuged (3800 rpm for 10 min at RT), and the supernatant discarded. The bacterial pellets were then resuspended in 4 ml of R medium, and the bacterial suspension was passaged 15 times through a 26G needle.

The cultures (live bacteria) and prepared gamma-irradiated bacteria were then placed in fresh 15 ml conical tubes and sonicated at full power in a Bransonic 2510 ultrasonic cleaner (Branson Ultrasonics) for 1 min. The OD_600_ nm values of the cultures and the preparation were then determined, and cell numbers were estimated based on an OD_600_ nm of 0.1 corresponding to 1 × 10^7^ bacterial cells. The cell numbers were then adjusted to 5 × 10^5^ bacterial cells/ml in pre-warmed R medium to achieve a multiplicity of infection (MOI) of one bacillus per bAM cell (1:1).

### Bovine alveolar macrophage challenge

All *in vitro* bAM challenges were performed in a BCL3 laboratory. The challenged bAM groups were *M. bovis* (MBO), *M. tuberculosis* (MTU), *M. bovis* BCG (BCG), and gamma-irradiated (killed) *M. bovis* (IRR); a non-challenged bAM control group (CON) was also run in parallel. The R medium from all tissue culture plate wells containing bAM (5 × 10^5^ cells/well) was removed and replaced with 1 ml of the prepared live or gamma-irradiated strains in R media (5 × 10^5^ bacteria/ml). The parallel wells for non-challenged control bAM samples received 1 ml R medium only. Once challenged, the bAM were incubated for bacterial uptake at 37°C, 5% CO_2_ for 3 h. After 3 h of bacterial uptake, the media for all wells used for analyses at 24 h of challenge was removed, the wells washed twice with pre-warmed PBS, and then filled with 1 ml fresh, clean R medium. The plates were then reincubated at 37°C and 5% CO_2_ for 24 h. For the wells required for fluorescence microscopy analysis at 0 h, the media was removed, and cells were directly treated according to sampling requirements without washing.

### Colony-forming unit (CFU) counts

The CFU counts were obtained using the mycobacterial challenge dose at the time of bacterial uptake. All counts were processed by plating serial dilutions on Middlebrook 7H11 agar plates (Difco^™^) with 0.05% Tween 80, 0.2% glycerol, 0.5% BSA, 0.2% glucose, and 0.085% NaCl supplemented with 10 mM sodium pyruvate (Sigma-Aldrich). For all CFU counts, 10-fold serial dilutions were prepared in triplicate in 96-well plates, with 360 μl of Middlebrook 7H9 medium and 40 μl of the live bacteria suspension to be examined added to the first well and mixed by vigorous pipetting. Serial dilutions were then performed on the 96-well plates using the same relative volumes, from 1:10 to 1:100,000. Subsequently, 50 μl of each serial dilution was plated onto 7H11 agar plates. Plates were placed in plastic bags (to prevent drying out) and stored at 37°C for up to 6 weeks, with colonies checked weekly from the second week after plating before final counting.

### Fluorescence microscopy

The bAM cell culture wells containing the coverslips were washed once with pre-warmed PBS, fixed with 1 ml 4% paraformaldehyde (Thermo Fisher Scientific) in PBS for 30 min at RT, and then stored at 4°C. The coverslips were prepared for fluorescence microscopy using the following protocol: the mycobacteria were stained by washing the wells once with PBS, then covering the coverslips with Modified Auramine-O stain (Scientific Device Laboratory) for 2 min at RT. The wells were washed once with PBS, and the coverslips were covered with modified Auramine-O Quencher decolouriser (Scientific Device Laboratory) for 2 min at RT, followed by one PBS wash. Cells and nuclei were then stained simultaneously with a staining mix of 2 µg/ml HCS CellMask^™^ Red stain (Thermo Fisher Scientific) and 10 µg/ml Hoechst stain (Thermo Fisher Scientific) in PBS for 5 min, followed by three washes with PBS. The coverslips were then removed from the wells with tweezers and mounted cell-side down, on a slide with 5 µl of mounting media.

Wide-field fluorescence microscopy was performed on a Zeiss Axio Imager M1 (Carl Zeiss Microscopy) within 2 h of the staining procedure. The excitation light intensity and exposure time settings were maintained across all conditions to facilitate quantitative comparisons. A 40× magnification was used to achieve appropriate resolution and a large field of view, with 10 fields per condition. The microscopy images were analysed using the CellProfiler software v.4.2.4 (Stirling et al. 2021). Identification of the biological objects was performed independently for each channel of the composite microscopy image. For nuclei segmentation, a global thresholding strategy based on the Otsu method was applied to the blue channel of the composite image. Adjacent merged nuclei were separated into individual objects of interest by shape and intensity. The same thresholding strategy and method were used for cell segmentation, with adjacent cells separated by intensity. Image processing was performed on the green channel of the composite image to enhance the bacteria, using the Tubeness method, before measuring Neurite features. The processed image was used for bacteria segmentation, with global thresholding using the Otsu method. Clumped bacteria were separated by intensity, with a single central peak of brightness identified for each organism. Object relationships were then assigned: nuclei were related to parent cell objects, and bacteria were related to parent cell objects. The CellProfiler analysis for each image provides the total number of cells, the number of nuclei per cell, and the number of bacteria per cell.

### Sampling and preparation of RNA for RNA-seq, and DNA for ChIP-seq and ATAC-seq

To extract total RNA from cultured bAM, the media from 5 wells per condition (∼ 2.5 × 10^6^ cells) was removed and replaced with a total of 700 μl of QIAzol lysis reagent (Qiagen), which was then transferred sequentially across the 5 wells with pipetting to achieve disruption and lysis of the cells. RNA purification and separation of small RNA-enriched fractions (<50 nucleotides) from larger RNAs was performed using a miRNeasy Mini kit (Qiagen) according to the manufacturer’s instructions. RNA quantity and quality were assessed using a NanoDrop™ 1000 spectrophotometer (Thermo Fisher Scientific) and an Agilent 2100 Bioanalyzer using an RNA 6000 Nano LabChip kit (Agilent Technologies). Purified RNA samples for mRNA sequencing were stored at −80°C before shipping to a commercial provider for RNA-seq library preparation and sequencing (Azenta Life Sciences, Leipzig, Germany).

For ChIP-seq sample preparation, bAM cells were fixed by adding 1/10 volume of freshly prepared 11% formaldehyde to the fixation buffer (0.1 M NaCl, 1.0 mM EDTA, 0.5 mM EGTA, 50 mM HEPES, pH 7.6) directly to the cell culture media in each well. The bAM cells were incubated with gentle shaking at RT for 8 min. Fixation was stopped by adding 1/10 volume of 1.25 M glycine, and the cells were then incubated with gentle shaking for a further 5 min at RT. Using a cell scraper, the bAM cells were collected, centrifuged (1000 × g for 10 min at 4°C), and washed once with ice-cold PBS. The cell pellets were then frozen immediately at −80°C before shipment to a commercial service provider (Diagenode SA, Liège, Belgium) for downstream laboratory processing. Chromatin for the ChIP-seq analyses was prepared by Diagenode from bAM cell pellets (∼ 1 × 10^7^ cells each using the Bioruptor^®^ Pico sonication device (Diagenode) and the iDeal ChIP-seq kit for Transcription Factors (Diagenode) according to the manufacturer’s instructions. The ChIP assays were performed using 200 ng of chromatin for each immunoprecipitation with antibodies for H3K4me1, H3K4me3, H3K27ac, H3K27me3, and CTCF according to the Functional Annotation of Animal Genomes (FAANG; www.faang.org) recommendations (Clark et al. 2020). Purified DNA samples were stored at −20°C prior to ChIP-seq library preparation and sequencing (Diagenode).

Cell preparation and tagmentation for ATAC-seq were performed with bAM cells, which were washed once with PBS containing 0.1% BSA. The bAM cells were then dissociated by adding pre-warmed non- enzymatic cell dissociation buffer in PBS (Thermo Fisher Scientific) to the wells and incubating at RT for 10 min. Gentle scraping of the cells with a cell scraper was performed after 5 min, followed by the addition of R media after 10 min. The bAM cells were collected and pelleted (500 × g for 10 min at 4°C), then resuspended in 500 μl R medium. The number of viable cells was determined using 0.4% Trypan Blue Solution, and cells were counted using a haemocytometer. As determined by cell counting, 50,000 cells were collected and centrifuged (500 × g for 10 min at 4°C); the supernatant was discarded, and the cell pellets were maintained on ice. Permeabilisation and transposition reactions were performed simultaneously on whole cells by resuspending gently in 50 μl of the permeabilisation/transposition mix (Diagenode), then incubating for 30 min at 37°C with mixing at 500 rpm. The samples were then transferred to ice, and the DNA was purified using the Diapure kit (Diagenode) according to the manufacturer’s instructions. Purified DNA samples were stored at −20°C before shipping to a commercial provider for ATAC-seq library preparation and sequencing (Diagenode).

### RNA-seq, ChIP-seq, and ATAC-seq library preparation and sequencing

Illumina-compatible DNA libraries (30 samples: 6 biological replicates for 5 conditions) were prepared with polyA selection for strand-specific RNA-seq and used for paired-end sequencing (2 × 150 bp) on the Illumina NovaSeq 6000 platform (Azenta Life Sciences). Illumina-compatible DNA libraries (75 samples: 3 biological replicates for 5 conditions and 5 histone marks) were also prepared for ChIP-seq and used for paired-end sequencing (2 × 50 bp) on the Illumina NovaSeq 6000 platform (Diagenode). In addition, Illumina-compatible DNA libraries (15 samples: 3 biological replicates per condition) were prepared for ATAC-seq and sequenced in paired-end mode (2 × 50 bp) on the Illumina NovaSeq 6000 platform (Diagenode). The cattle bAM sample IDs and the allocations to the different functional genomics assays are detailed in Additional file 2: Table S1.

### RNA-seq bioinformatics

Computational analyses for all RNA-seq bioinformatic processes were performed on a 72-CPU Linux Ubuntu (version 18.04.4 LTS) server. Following RNA-seq data generation and pre-processing, a mean of 70.43 million (M) paired-end 150 bp reads per individual library were obtained (*n* = 29 libraries) (Additional file 3: Table S3). The RNA-seq datasets were *M. bovis*-challenged (MBO, *n = 5*); *M. tuberculosis*-challenged (MTU, *n* = 6); *M. bovis-*BCG-challenged (BCG, *n* = 6); gamma-irradiated *M. bovis*-challenged (IRR, *n* = 6); and control non-challenged (CON, *n* = 6). A single MBO RNA-seq sample (animal ID: 43095) was compromised by contamination and not included in subsequent analyses (Additional file 2: Table S1). Adapter sequence contamination and low-quality paired-end reads were filtered out using the FastP v.0.23.2 FASTQ file processor (Chen et al. 2018). At each step, read quality was assessed with FastQC v.0.12.0 (Andrews 2023). After quality control and filtering, a mean of 66.16 M reads per library were mapped to the *B. taurus* ARS-UCD1.2 reference genome (Rosen et al. 2020) using the STAR aligner v.2.7.1b (Dobin et al. 2013) with an average read mapping efficiency of 93.1%. Read counts for each gene were generated using featureCounts v.1.6.4 (Liao et al. 2014), set to unambiguously assign uniquely aligned paired-end reads in a stranded manner to gene exon annotations. Using the R statistical programming language v.4.3.1 (R Core Team 2023), gene annotation was obtained from an NCBI GFF annotation file (GCF_002263805.1) with additional descriptions and chromosomal locations annotated using the GO.db v.3.17.0 (Carlson 2019) and biomaRt v.2.56.1 (Durinck et al. 2009) packages.

Differential gene expression analysis was performed with the DESeq2 package v.1.40.2 (Love et al. 2014) for the 10 possible experimental group contrasts with five experimental groups (MBO, MTU, BCG, IRR, and CON). The differential expression analysis used the DESeq2 Wald test, and to obtain reliable variance estimates, shrunken log fold-changes were computed by combining information across all genes; these estimates were then used for subsequent analysis. Multiple testing correction was performed for each gene expression contrast using the Benjamini-Hochberg (B-H) false discovery rate (FDR) method (Benjamini and Hochberg 1995) and the default threshold for differentially expressed genes (DEGs) was an FDR-adjusted *P*-value (FDR-*P*_adj._) < 0.05.

### ChIP-seq bioinformatics

Computational analyses for all ChIP-seq bioinformatic processes were performed on a 72-CPU Linux Ubuntu (v.18.04.4 LTS) server. Following ChIP-seq data generation and pre-processing, a mean of 129.78 M paired-end 50 bp reads were obtained from three biological replicates per experimental group (MBO, MTU, BCG, IRR, and CON) for each histone mark (H3K27ac, H3K4me3, H3K27me3, and H3K4me1), CTCF binding, and input controls (Additional file 4: Table S12). Therefore, for each of the three biological replicates across the five experimental groups, there were input libraries for five ChiP-seq assays, except for the AS41_H3K4me1 control (CON) group, which did not yield sufficient material for sequencing. Consequently, 74 ChIP-seq datasets were produced (Additional file 4: Table S12‒13). At each step of data processing, read quality was assessed using FastQC v.0.12.0. Reads were aligned to the ARS-UCD1.2 reference genome using the BWA software package v.0.7.5a (Li and Durbin 2009) with the BWA-backtrack algorithm, yielding a mean mapping efficiency of 93.3%. Samples were filtered for regions blacklisted by the ENCODE project (Hoffman et al. 2013). PCR duplicates and multimapping reads were removed using Samtools v.1.13 (Danecek et al. 2021). Alignment coordinates were converted to BED format using BEDTools v.2.17 (Quinlan and Hall 2010), and peak calling was performed using epic2 v.0.0.52, a reimplementation of SICER with optimised parameters for each histone mark, as determined by the optimal peak calling protocol (Stovner and Sætrom 2019).

Peaks were called using alignment files to determine where reads aligned to specific regions of the genome, and then those alignments were compared to the input samples as a normalisation step. Peaks in all marks were visually assessed in parallel across the bovine reference genome with the Integrative Genomics Viewer (IGV) (Robinson et al. 2023). Differential peak calling was performed with the DiffBind v.3.10 (Ross-Innes et al. 2012) and DESeq2 v.1.40.2 R packages to identify differential affinity binding sites (DABSs) between the control group (CON) and the four experimental groups (MBO, MTU, BCG, and IRR), and then combined into a DABS score matrix and summarised into a comparative table (Additional file 6: Table S23). Peaks for each biological sample and experimental condition for each mark were cross-referenced with the IGV images and the differential peak caller to generate a fold enrichment value for each observed peak difference between conditions. This procedure required comparing and evaluating peak start and end sites, chromosomal locations, *P*-values, and FDR-*P*_adj._ values for each peak summit, as well as the normalised fold enrichment of each peak relative to the input sample, and the comparison of that enrichment between the control group (CON) and the challenged groups (MBO, MTU, BCG, and IRR).

Differential binding affinity analyses were performed using the dba.analyze function in DiffBind. DiffBind includes functions to support the processing of peak sets, the inclusion of overlapping and merging peak sets, sequence read counting, the overlapping of intervals in peak sets, and the identification of statistically significantly differentially bound sites based on evidence of binding affinity measured by differences in read densities (Additional file 4: Table S15–18). Parallel normalisation using the CTCF datasets was implemented as described in the DiffBind vignette, as no differences were observed among groups in CTCF binding. The default criterion for identification of a DABS was FDR-*P*_adj._ < 0.05. All detected DABS were then annotated using the Hypergeometric Optimisation of Motif EnRichment (HOMER) software suite v.4.11 (Heinz et al. 2010). HOMER provides various tools for motif discovery, annotation, and visualisation.

It leverages the hypergeometric distribution to identify significant motif enrichments and provides extensive options for analysing ChIP-seq, RNA-seq, and ATAC-seq data. The process of annotating peaks or regions consisted of two main steps: an initial step that identifies the closest transcription start site (TSS) and associates the peak with the corresponding gene; a second step then determines the annotation for the genomic interval encompassing the peak or region’s centre. Importantly, HOMER can be used with a custom genome annotation, such as GCF_002263795.1_ARS-UCD1.2_genomic.gtf, making it a suitable tool for analysing bovine data and genome annotation.

### ATAC-seq bioinformatics

Computational analyses for all ATAC-seq bioinformatic processes were performed on a 72-CPU Linux Ubuntu (version 18.04.4 LTS) server. Following ATAC-seq data generation and pre-processing, a mean of 100.45 M paired-end 50 bp sequence reads were obtained for each sample (Additional file 5: Table S19) and read QC was performed using FastQC v.0.12.0. Reads were aligned to the ARS-UCD1.2 reference genome using Bowtie2 v.2.4.5 (Langmead and Salzberg 2012) with a mean mapping efficiency of 95.07%. Samples were filtered for regions blacklisted by the ENCODE project and deduplicated using Picard v.2.26.10 (http://broadinstitute.github.io/picard). Alignment coordinates were converted to BED format using BEDTools v.2.17, and peak calling and fragment-in-peak scores (FriP) were calculated using MACS2 v.2.1.2 (Zhang et al. 2008; Gaspar 2018) with customised parameters for ATAC-seq data. For consistency, ATAC-seq peak analysis closely followed the ChIP-seq analysis pipeline, and peaks were visually assessed across the bovine reference genome using IGV. Differential peak calling was performed with DiffBind v.3.10 and DESeq2 v.1.40.2 to identify differential open chromatin regions (DOCRs) between the control group (CON) and the four experimental groups (MBO, MTU, BCG, and IRR). These results were then combined into a DOCR score matrix and summarised into a comparative table (Additional file 6: Table S23). Peaks from each biological sample for each condition and mark were cross-referenced with the IGV images and the differential peak caller to generate a fold enrichment value for each observed peak difference between conditions. This procedure required cataloguing and evaluating peak start and end sites, chromosomes, *P*-values and FDR-*P*_adj._ values for each peak summit, peak summit locations, normalised fold enrichment of a peak against the input sample, and comparisons of that enrichment between the control group (CON) and each of the challenged groups (MBO, MTU, BCG, and IRR). Differential binding affinity analyses were performed using the dba.analyze function in DiffBind v.3.10. The default criterion for identification of a DOCR was FDR-*P*_adj._ < 0.05 and all DOCR sites were then annotated using HOMER v.4.11 and the GCF_002263795.1_ARS-UCD1.2_genomic.gtf file as described for the ChIP-seq analysis.

### Multi-omics data integration

The DEG, DABS, and DOCR locations for the four primary bAM challenge contrasts (MBO vs. CON, MTU vs. CON, BCG vs. CON, and IRR vs. CON) were integrated based on gene location, HOMER annotation and any ChiP-seq peak, ATAC-seq peak, or RNA-seq expression log_2_ fold change (log_2_FC) with an FDR-*P*_adj._ < 0.05 (Additional file 6: Table S23). Correlation heatmaps were generated using genome-wide H3K4me1, H3K4me3, H3K27me3, and H3K27ac sequencing reads, along with RNA-seq data from each experimental group (MBO, MTU, BCG, IRR, and CON), using EaSeq v.1.05 (Lerdrup et al. 2016). Circle plots for each experimental group containing all peaks of H3K4me1, H3K4me3, H3K27me3, H3K27ac and ATAC-seq reads were generated using the circlize R package v.0.4.15 (Gu et al. 2014). BED files for each of the histone modification experimental contrasts from each sample were concatenated and visualised as stacked signal tracks, where the *x*-axis represents a two-point start and end location for a given read, and the *y*-axis shows the signal score value of that read, a measurement of overall enrichment for the region generated by MACS2. Multi-omic IGV track images for each infection group were generated by converting each binary alignment file for each ChIP-seq, ATAC-seq and RNA-seq sample to bigWig format using the deepTools bamCoverage program v. 3.5.0 (Ramírez et al. 2016), and then combining and merging overlapping peaks in all samples for each dataset. Peak images were generated by visually assessing all bigWigs in tandem across the entire bovine genome using IGV. Interaction networks were generated in Cytoscape v.3.10.0 (Shannon et al. 2003) and node colour was determined by the positive (red) or negative (blue) log_2_FC change for each gene. Lines (edges) between nodes were coloured by the increase in peak signal in the control bAM group (blue) or in the challenged bAM groups (red). Node size was determined by the number of connecting edges to that node from other nodes (degree). A gene/node was connected to the square histone hubs if that gene exhibited either a DABS or a DORC between the control bAM and the challenged bAM groups.

### Gene set enrichment analysis, overrepresentation analysis, and GWAS integration

The Ingenuity^®^ Pathway Analysis (IPA^®^) software tool (Krämer et al. 2014) with the Ingenuity^®^ Knowledge Base (Qiagen; release date December 2022) was used to perform pathway-focused gene set enrichment analysis (GSEA) using each of the ten possible transcriptomics (DEG) experimental group contrast datasets with further investigation of the results of four contrasts (MBO, MTU, BCG, and IRR vs. CON), and overrepresentation analysis (ORA) of a composite set of genes that also had DABS and/or DOCRs (DEG/DABS/DOCR set). This approach enabled the identification of canonical pathways and biological processes important to host-pathogen interactions for bAM challenged with *M. bovis, M. tuberculosis, M. bovis* BCG, and gamma-irradiated *M. bovis*. The statistical threshold used to generate suitable input gene sets for GSEA and ORA was FDR-*P*_adj._ < 0.05, unless a more stringent threshold was required to obtain an input gene set of an appropriate size (< 3000 genes) for IPA^®^ based on the IPA^®^ KnowledgeBase Core Analysis Setup recommendations. The IPA^®^ settings were configured with *Homo sapiens* as the target species, “Macrophage” as the cell type, all node types selected, and with the “Experimentally Observed” and “High Predicted” confidence settings. Following best practice, the background gene set used for GSEA and ORA was the set of detectable genes across all RNA-seq libraries for each time point contrast, not the complete bovine transcriptome (Timmons et al. 2015). The background gene set used for the ORA was the IPA^®^ Knowledge Base. Finally, the g:GOSt partition of the g.Profiler webtool (version release 19/04/2023) (Raudvere et al. 2019) was used to perform a gene ontology (GO) focused ORA using the composite set of DEG/DABS/DOCR genes.

The DEG/DABS/DOCR output gene set obtained from the multi-omics analyses was integrated with a published bTB *M. bovis* infection susceptibility trait GWAS dataset for Holstein-Friesian cattle (Ring et al. 2019). The gwinteR tool was used, which can test the hypothesis that a specific set of genes is enriched for signal in a GWAS dataset relative to the genomic background (Hall et al. 2021). The bTB GWAS dataset consisted of 12,740,315 autosomal SNPs with nominal *P*-values from an analysis of estimated breeding values (EBVs) derived from an *M. bovis* infection phenotype, generated for 1502 Holstein sires and described in detail elsewhere (Ring et al. 2019; Hall et al. 2021).

## Data and computer code availability

The bovine RNA-seq, ChIP-seq, and ATAC-seq datasets were generated by the authors and are available from the NCBI Gene Expression Omnibus (GEO) under the following accessions: RNA-seq data – GSE317895; ChIP-seq data – GSE317896; and ATAC-seq data – GSE317894. The bovine GWAS summary statistics were obtained from a published study that provides the information concerning sequence and genotype data availability (Ring et al. 2019). The code used to integrate the functional genomics outputs and GWAS datasets for this study is available on GitHub (https://github.com/ThomasHall1688/gwinteR).

## Author contributions

TJH, MM, SVG and DEM conceived and designed the study. MM and JAB performed experimental work. TJH, JFO’G, and GPM performed bioinformatics and computational analyses. ELC and MS provided input on bioinformatics and computational genomics. TJH and DEM wrote and prepared the manuscript and figures. MM wrote the laboratory-based methodology. All authors read and approved the final manuscript.

## Acknowledgements

The authors would like to thank Donagh Berry and Siobhán Ring for providing cattle GWAS data.

