## Additional File 1 for "Comparative multi-omics of the macrophage response to infection with *Mycobacterium tuberculosis* complex bacteria reveals pathogen-driven epigenomic reprogramming"

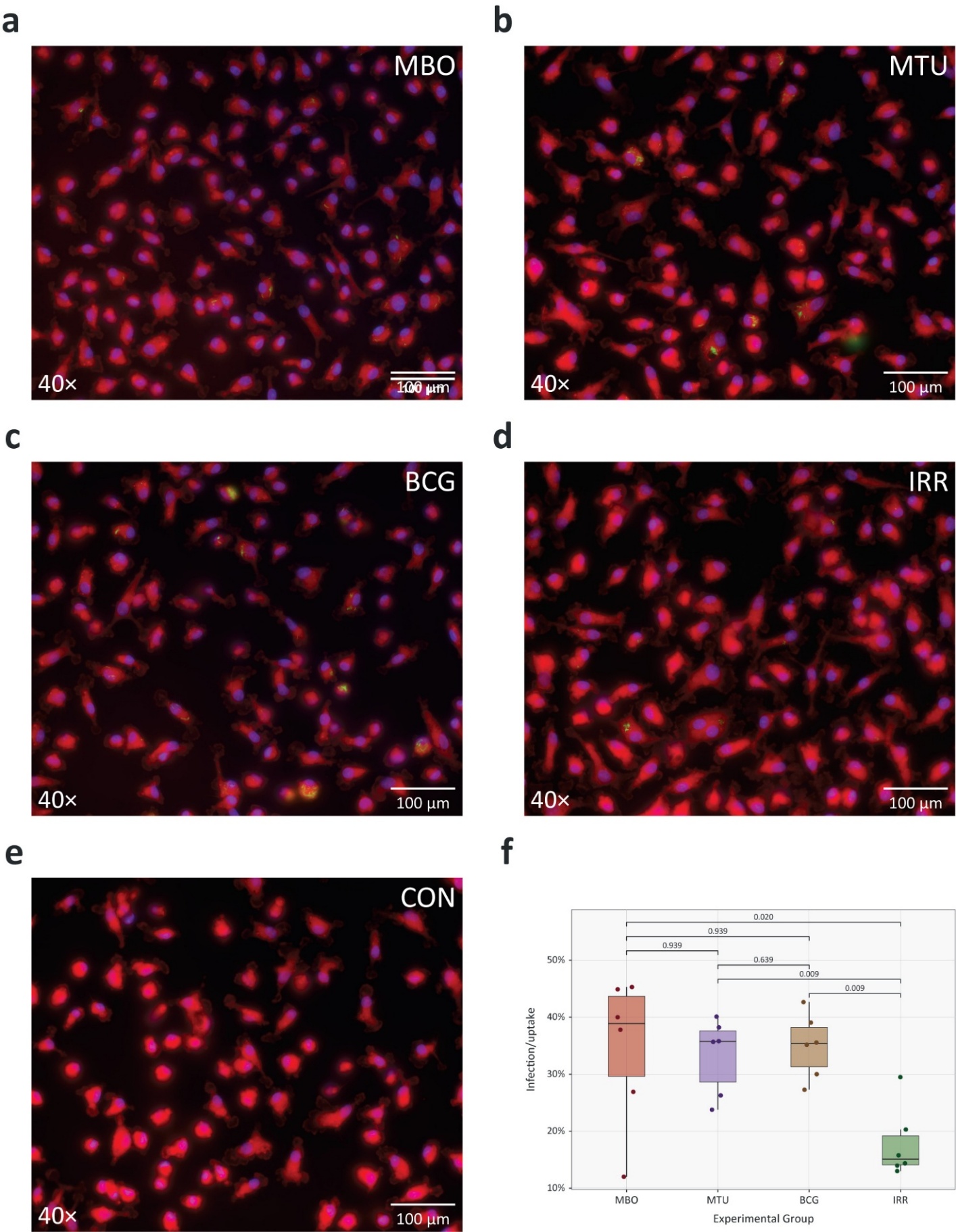

**Fig. S1:** Micrographs and statistical analysis of MTBC bacterial infection/uptake by bovine alveolar macrophages (bAM). The bAM are labelled with HCS CellMask™ Red stain, and MTBC cells are labelled with purple Hoechst stain. **a** *Mycobacterium bovis* (MBO). **b** *M. tuberculosis* (MTU). **c** *M. bovis* BCG (BCG). **d** gamma-irradiated (killed) *M. bovis* (IRR). **e** Control non-infected bAM. **f.** Box plots of infection/uptake data showing results for the four MTBC challenges (from ten image replicates each). The median, the upper/lower quartile and the minimum/maximum ranges are shown. The FDR-*P*_adj._ values for the *t*-test results comparing the four challenge groups are also shown.

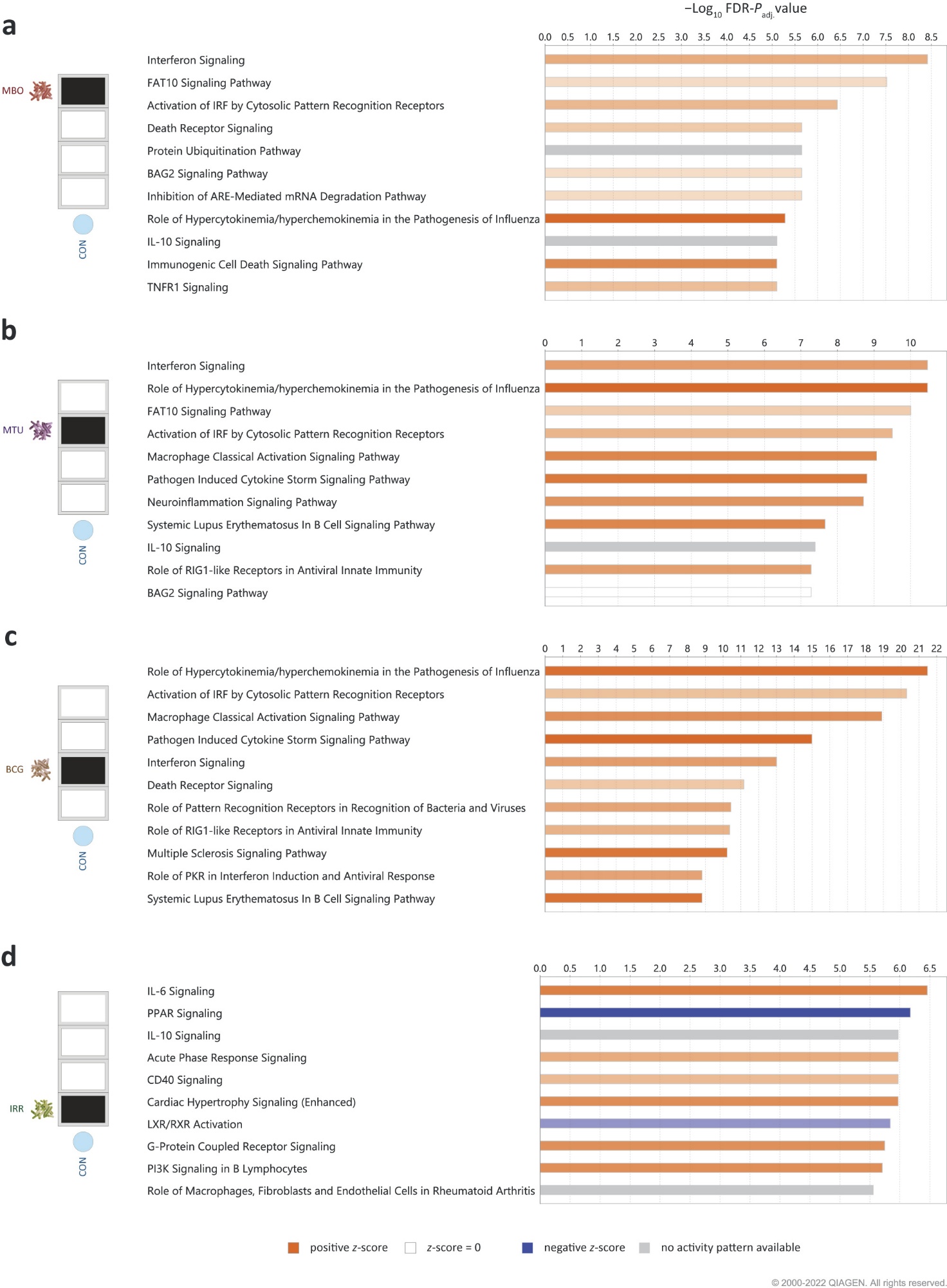

**Fig. S2:** Top ten enriched pathways detected using Ingenuity Pathway Analysis (IPA®) for the four primary MTBC bAM challenges. **a** *Mycobacterium bovis* (MBO). **b** *M. tuberculosis* (MTU). **c** *M. bovis* BCG (BCG). **d** gamma-irradiated (killed) *M. bovis* (IRR).

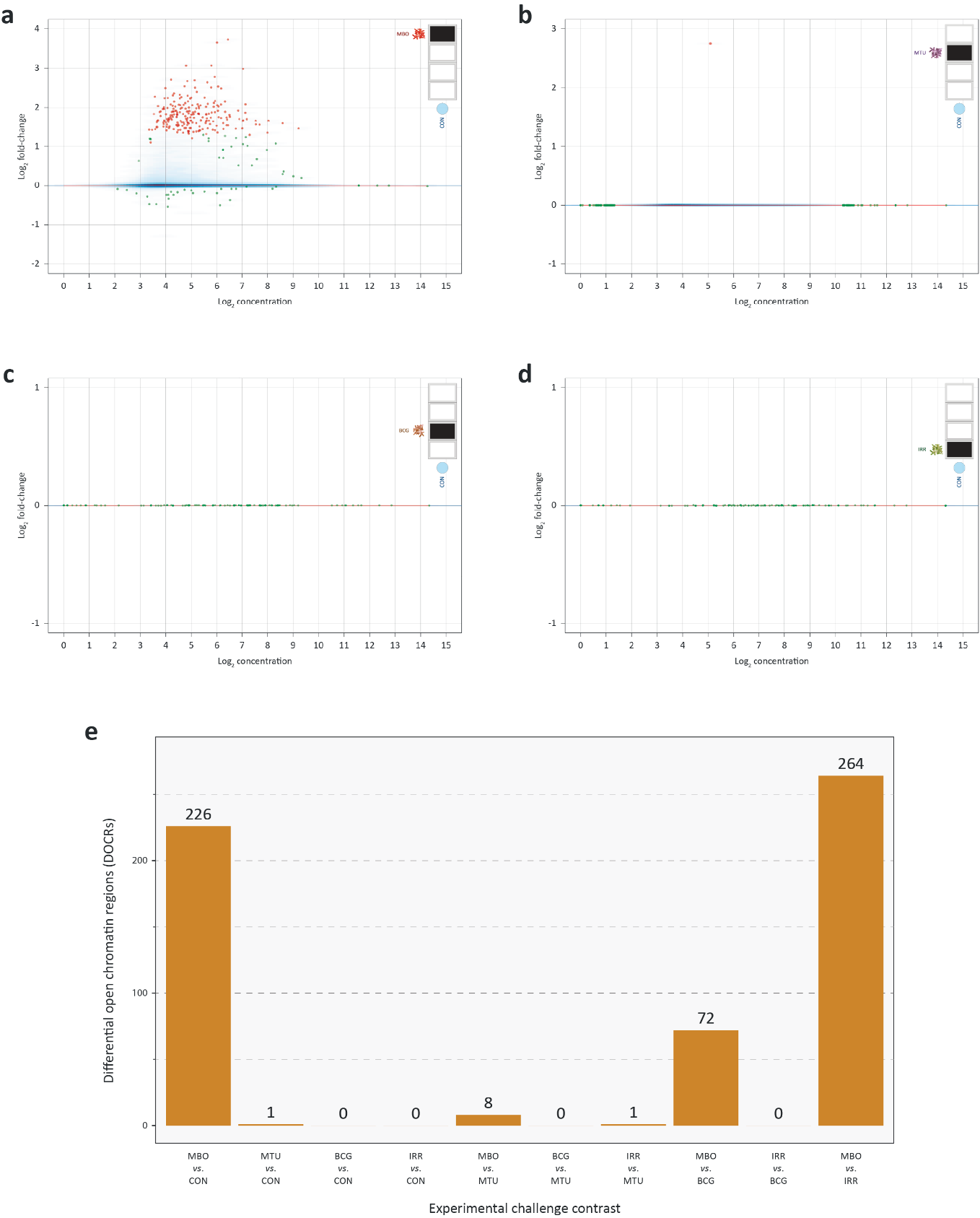

**Fig. S3:** ATAC-seq differential open chromatin region (DOCR) results for bAM challenged with MTBC. For panels a–d, the *x*-axis shows the DOCR log_2_ concentration values for the four primary MTBC bAM challenges with larger values indicating increased accessibility in MTBC-challenged bAM samples compared to non-challenged controls (CON). The *y*-axis shows the log_2_ fold-change values for chromatin accessibility (differential binding) between the MTBC-challenged and CON samples. Red data points indicate regions that show a statistically significant increase in accessibility after the MTBC challenge, while green data points indicate regions that show a statistically significant decrease in accessibility after the MTBC challenge. **a** *Mycobacterium bovis* vs control (MBO/CON). **b** *M. tuberculosis* vs control (MTU/CON). **c** *M. bovis* BCG vs control (BCG/CON). **d** gamma-irradiated (killed) *M. bovis* vs control (IRR/CON). **e** Bar chart showing DOCR results for all ten bAM challenge contrasts.

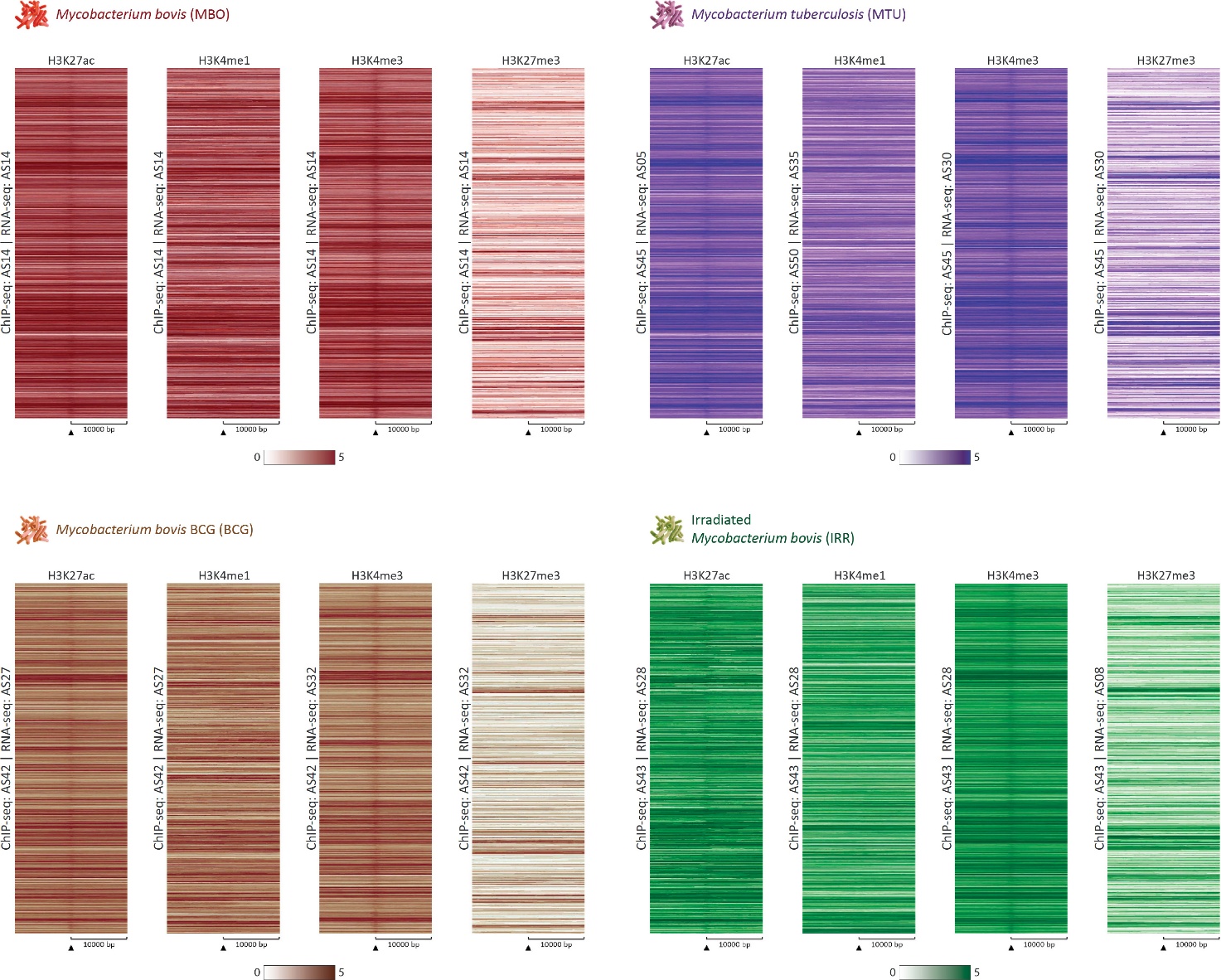

**Fig. S4:** Heat maps recoloured from greyscale showing the RNA-seq reads at the ChIP-seq set of regions for each histone modification and for each of the four MTBC challenges. The laboratory sample codes for each ChIP-seq and RNA-seq sample dataset are shown vertically beside each heat map. The Y-axis represents the individual positions of the regions, and the X-axis represents a surrounding area corresponding to 20000% of each ChIP-seq region, segmented into 200 bins. Regions were sorted according to original order. A scale bar showing the relationship between colour and signal intensity is provided below each group of ChIP-seq mark heat maps.

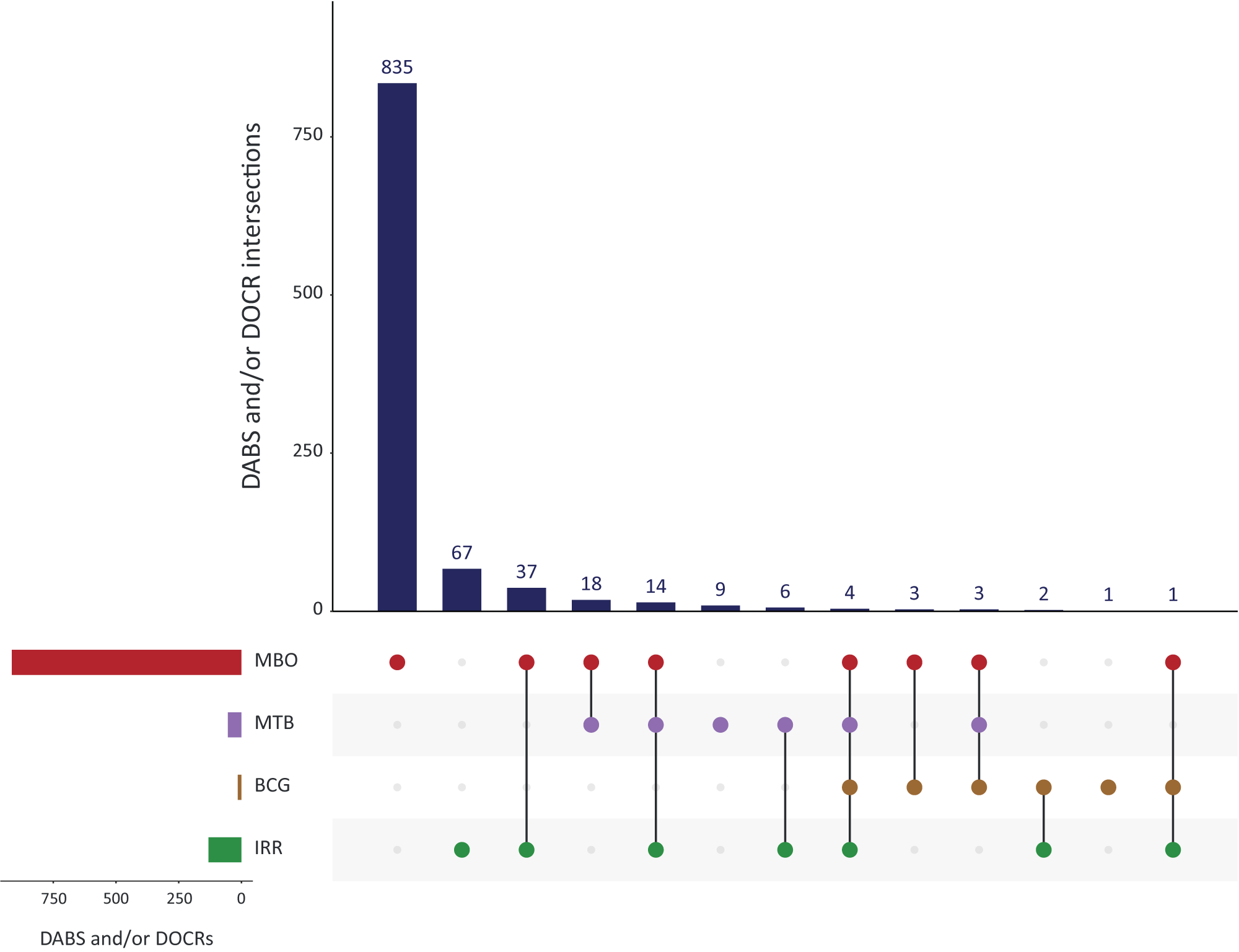

**Fig. S5:** UpSet plot showing the intersection of the consolidated multi-omic DAB/DOCR 1000 gene set.

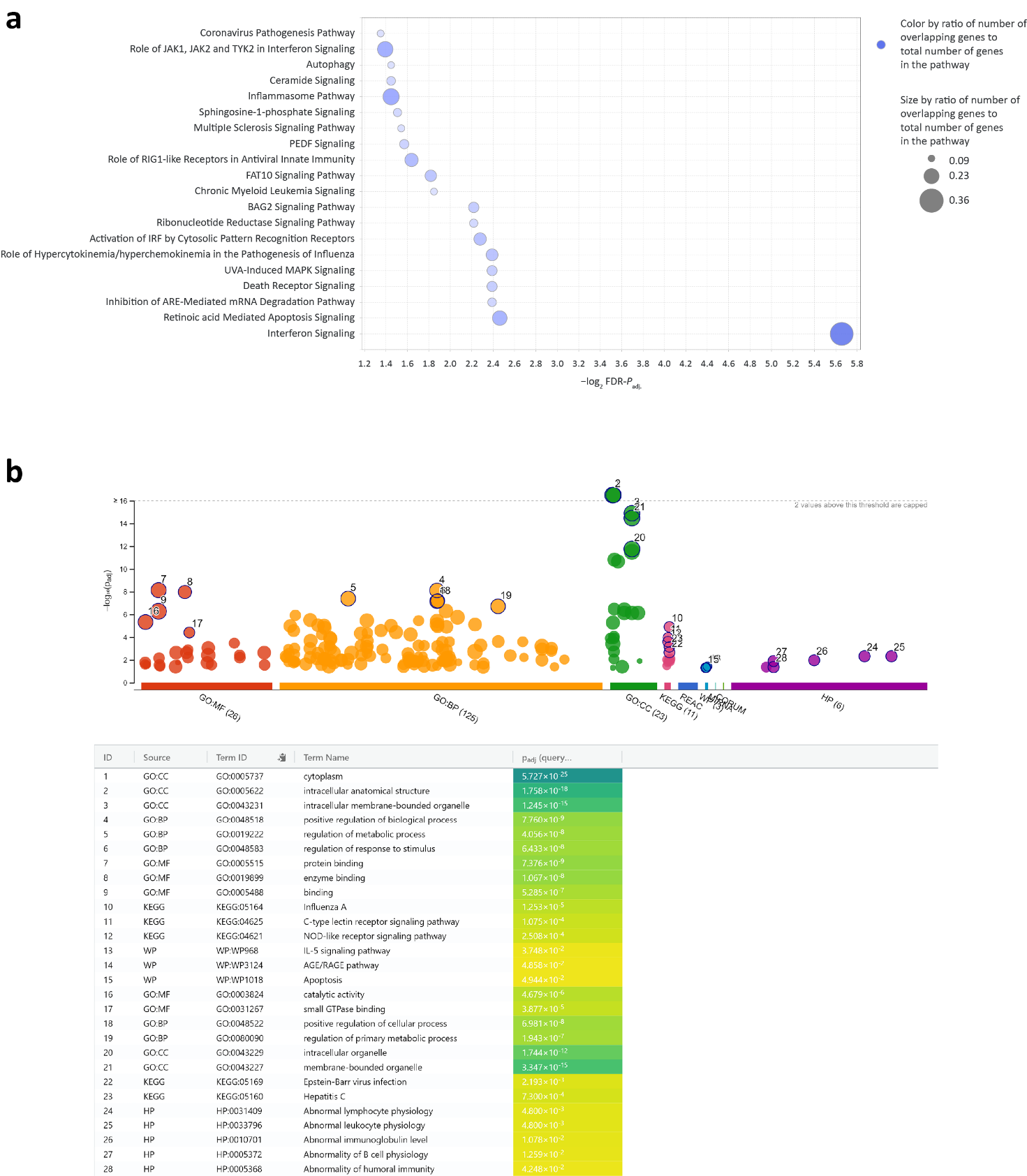

**Fig. S6:** Results of gene set overrepresentation analyses for the consolidated multi-omic DAB/DOCR 1000-gene set. **a** Bubble plot showing the 20 statistically significant overrepresented canonical pathways identified using IPA®. The colour and size of the bubbles correspond to the ratio of overlapping genes in the 1000-gene set and the total number of genes in a canonical pathway. **b** Gene set overrepresentation chart and ranked list of the top 28 functional category entities obtained using g:Profiler. The functional categories included were Gene Ontology (GO) molecular function (GO:MF), GO biological process (GO:BP), GO cellular component (GO:CC), KEGG Pathway (KEGG), Reactome (REAC), WikiPathways (WP), mirTarBase (MIRNA), CORUM protein complexes (CORUM), and Human Phenotype Ontology (HP) [1].

**Fig. S7:** Integrated Genome Viewer (IGV) track visualisations for ChIP-seq (H3K27ac, H3K4me3, H3K4me1), RNA-seq, and ATAC-seq data in the five bAM challenge groups (MBO, MTU, BCG, IRR, and CON) shown on the following four pages. **a** BCL2 apoptosis regulator gene (*BCL2*). **b** CD274 molecule gene (*CD274*). **c** 2'-5'-oligoadenylate synthetase 2 gene cluster (*OAS2*, *OAS1Y*, *OAS1X*, and *OAS1Z*). **d** transmembrane serine protease 2 gene (*TMPRSS2*).

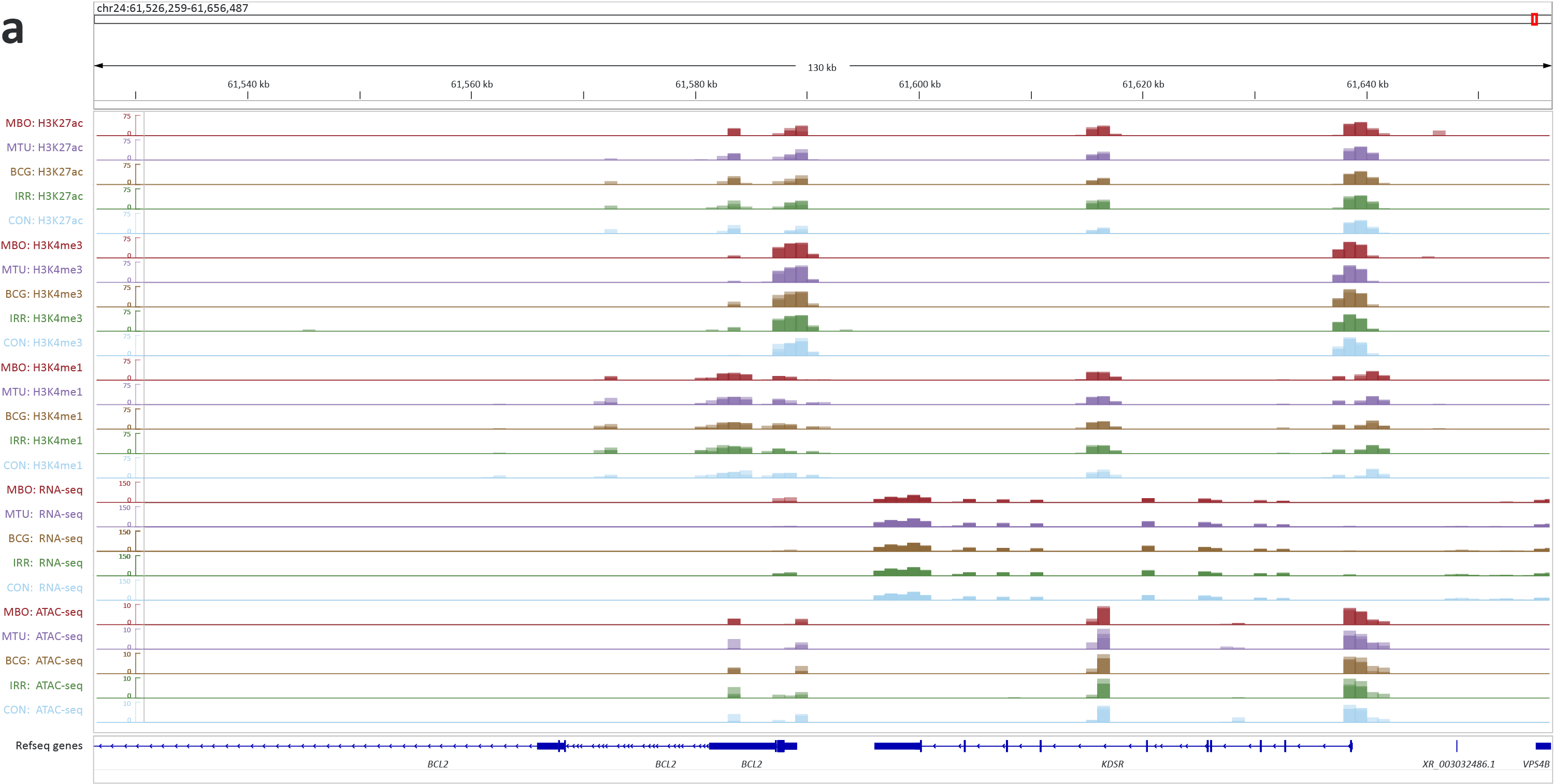

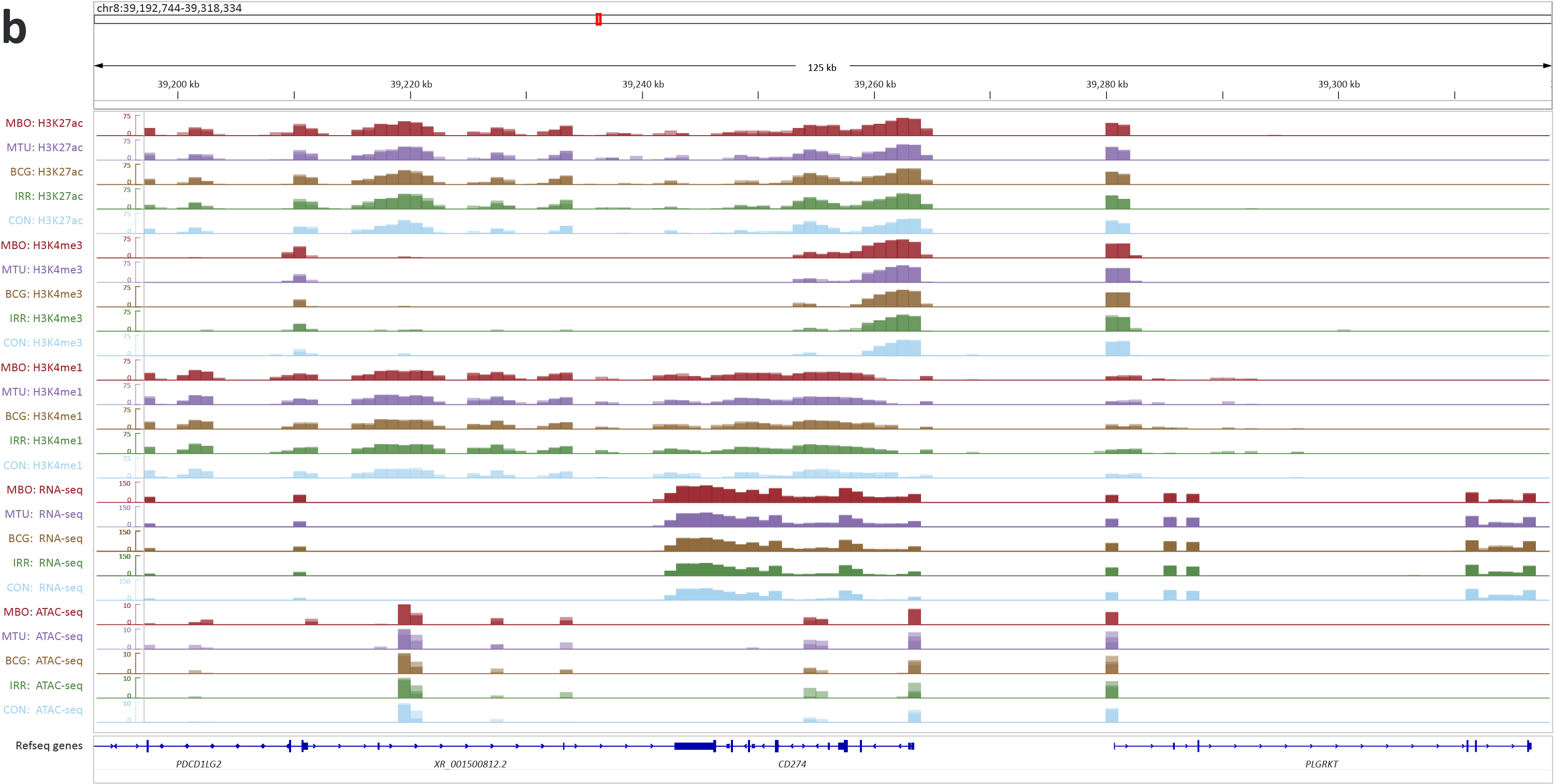

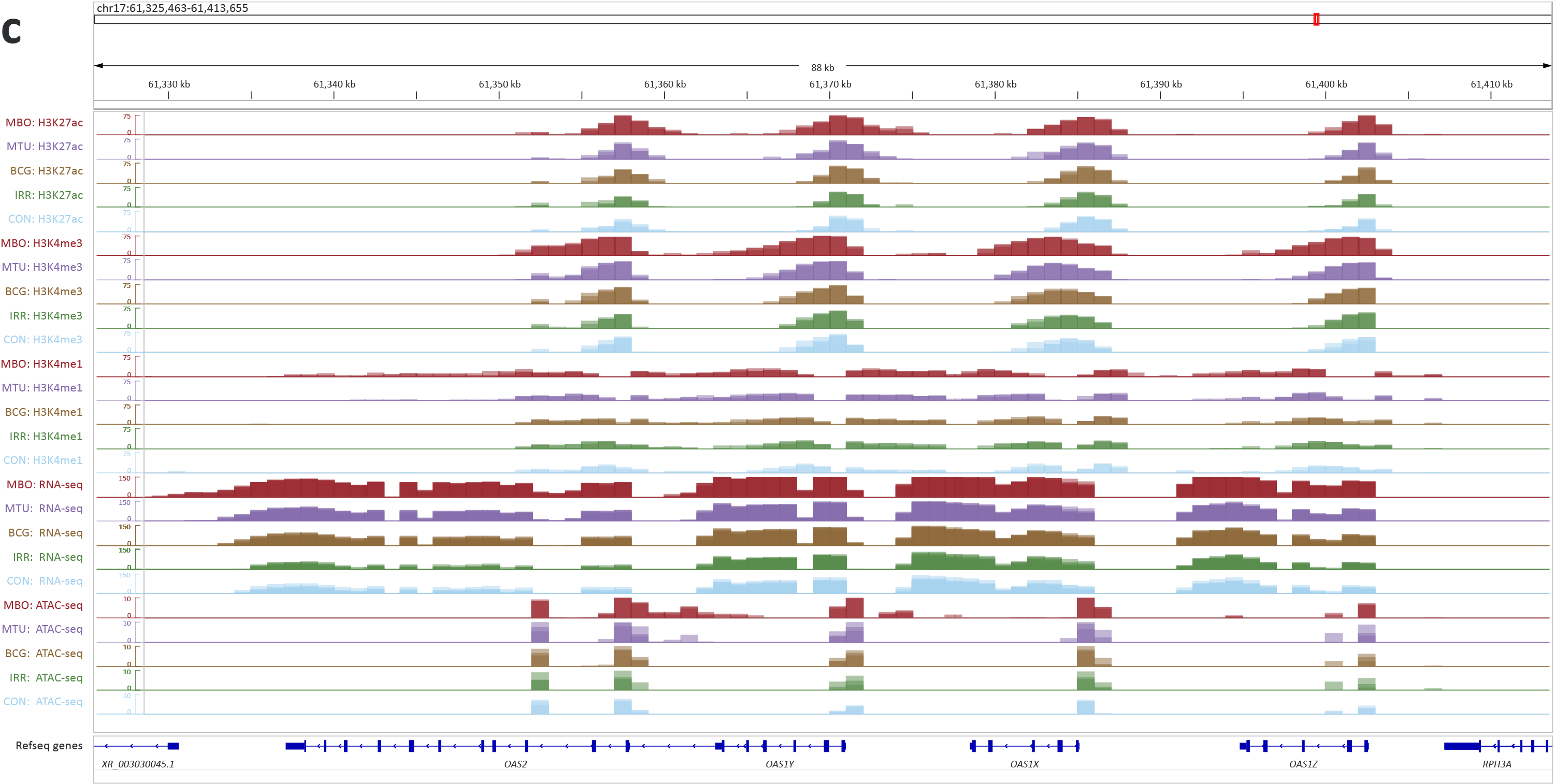

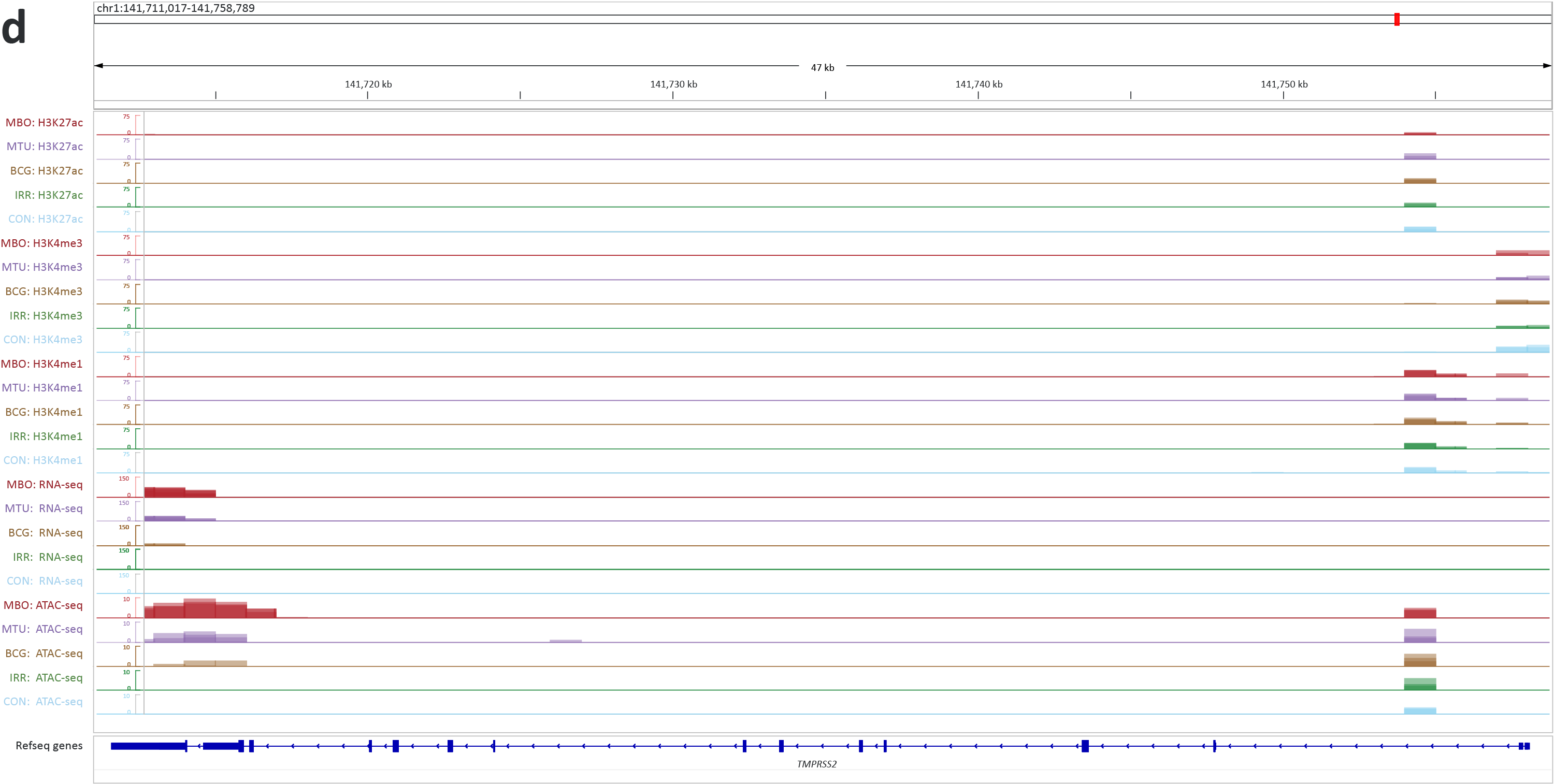

**Fig. S8:** Graphs of DABS and DOCR gene log_2_FC values plotted against RNA-seq log_2_FC values for the four primary MTBC bAM challenges (MBO/CON, MTU/CON, BCG/CON, and IRR/CON) shown on the following six pages. **a** MBO: H3K27ac vs RNA-seq. **b** MBO: H3K4me3 vs RNA-seq. **c** MBO: H3K4me1 vs RNA-seq. **d** MBO: ATAC vs RNA-seq. **e** MTU: H3K27ac vs RNA-seq. **f** MTU: H3K4me1 vs RNA-seq. **g** MTU ATAC vs RNA-seq. **h** BCG: H3K27ac vs RNA-seq. **i** BCG: H3K4me1 vs RNA-seq. **j** IRR: H3K27ac vs RNA-seq. **k** IRR: H3K4me3 vs RNA-seq. **l** IRR: H3K4me1 vs RNA.

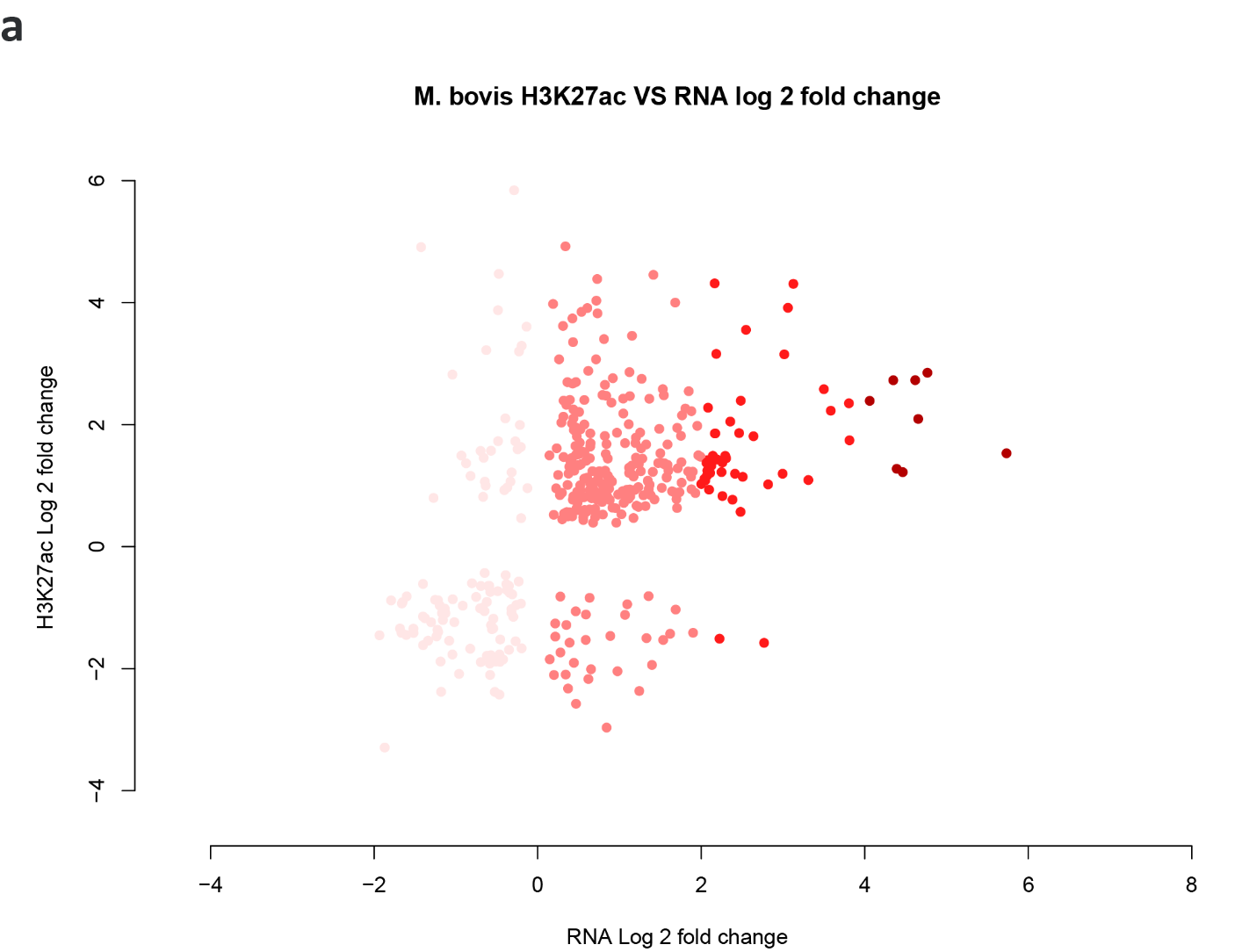

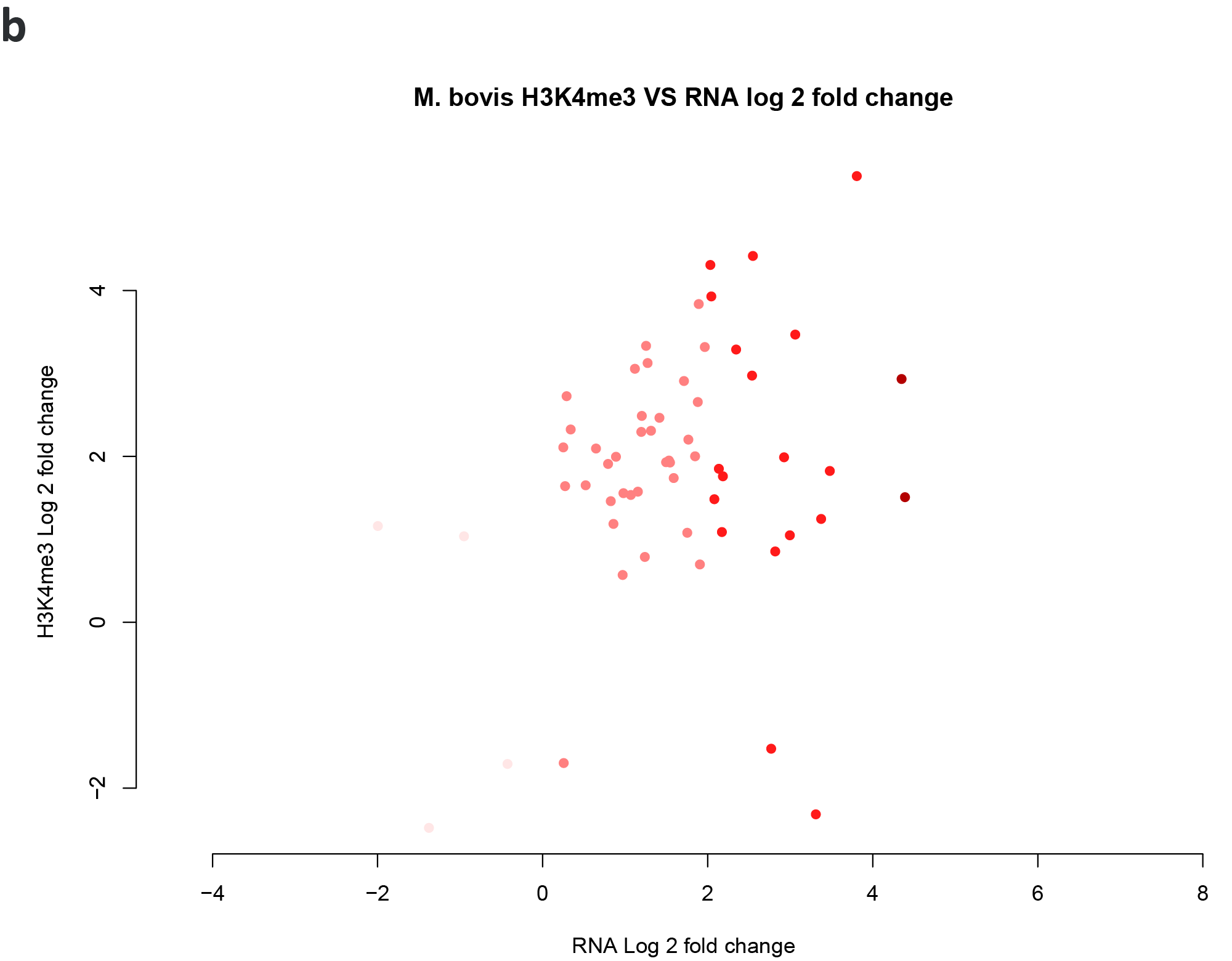

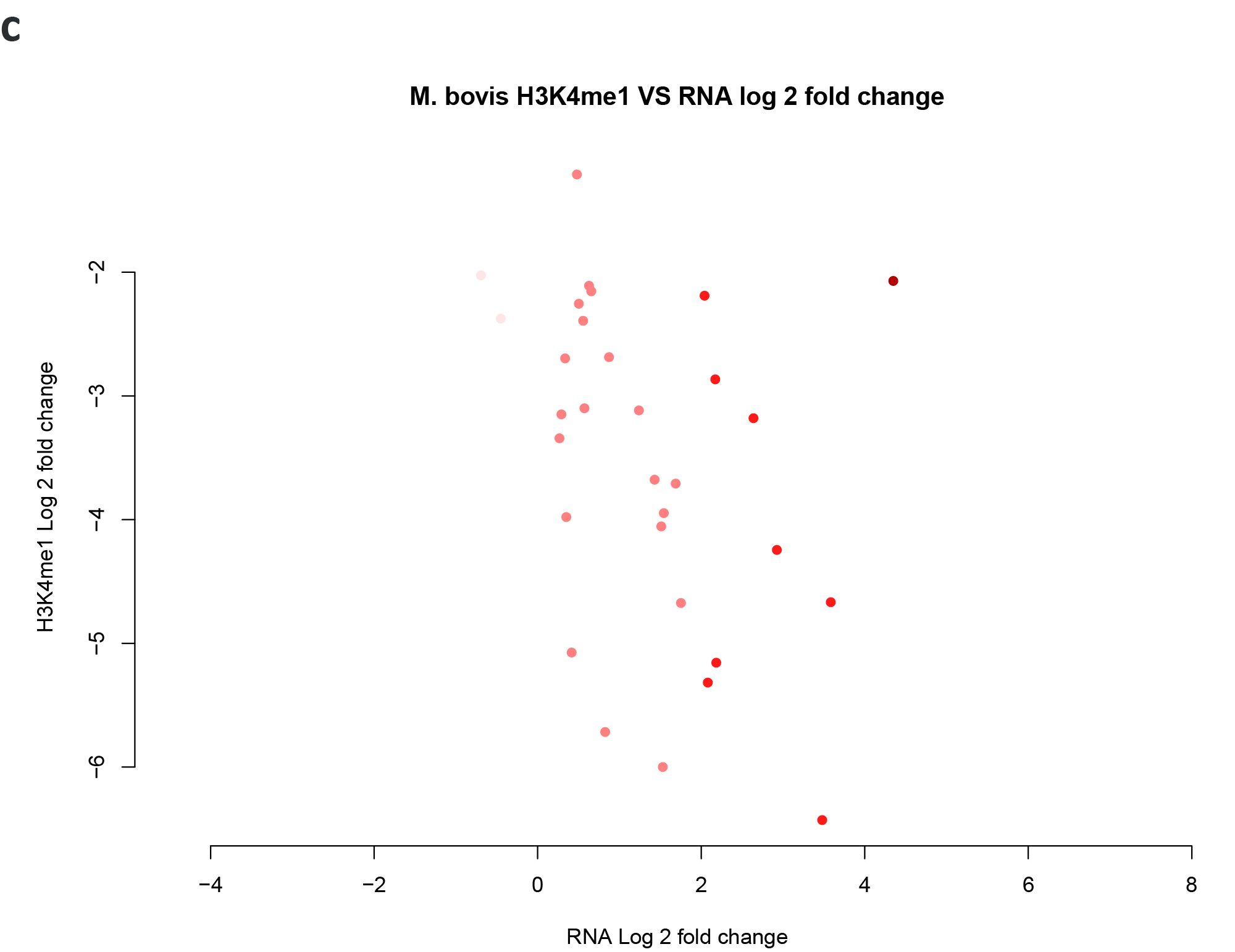

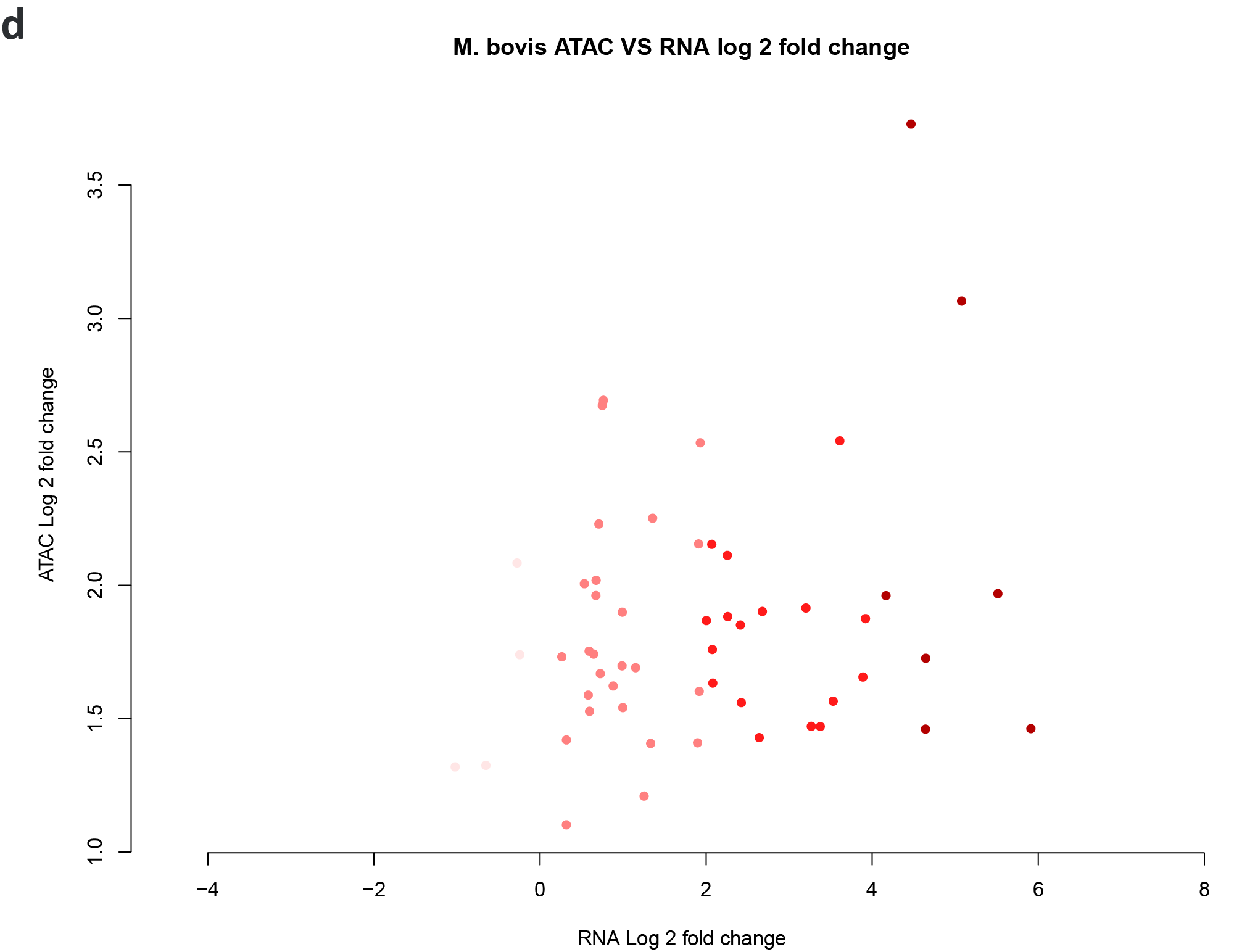

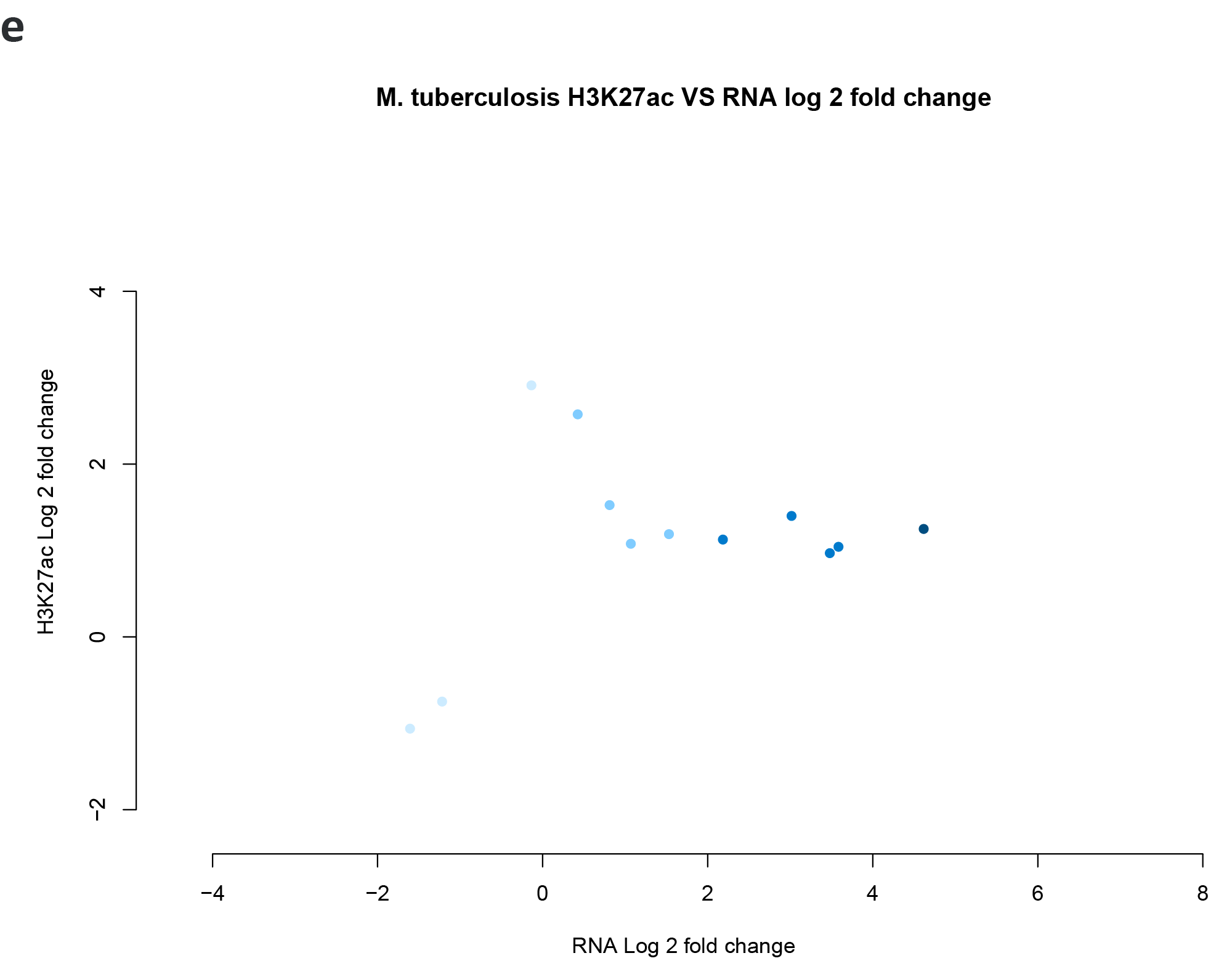

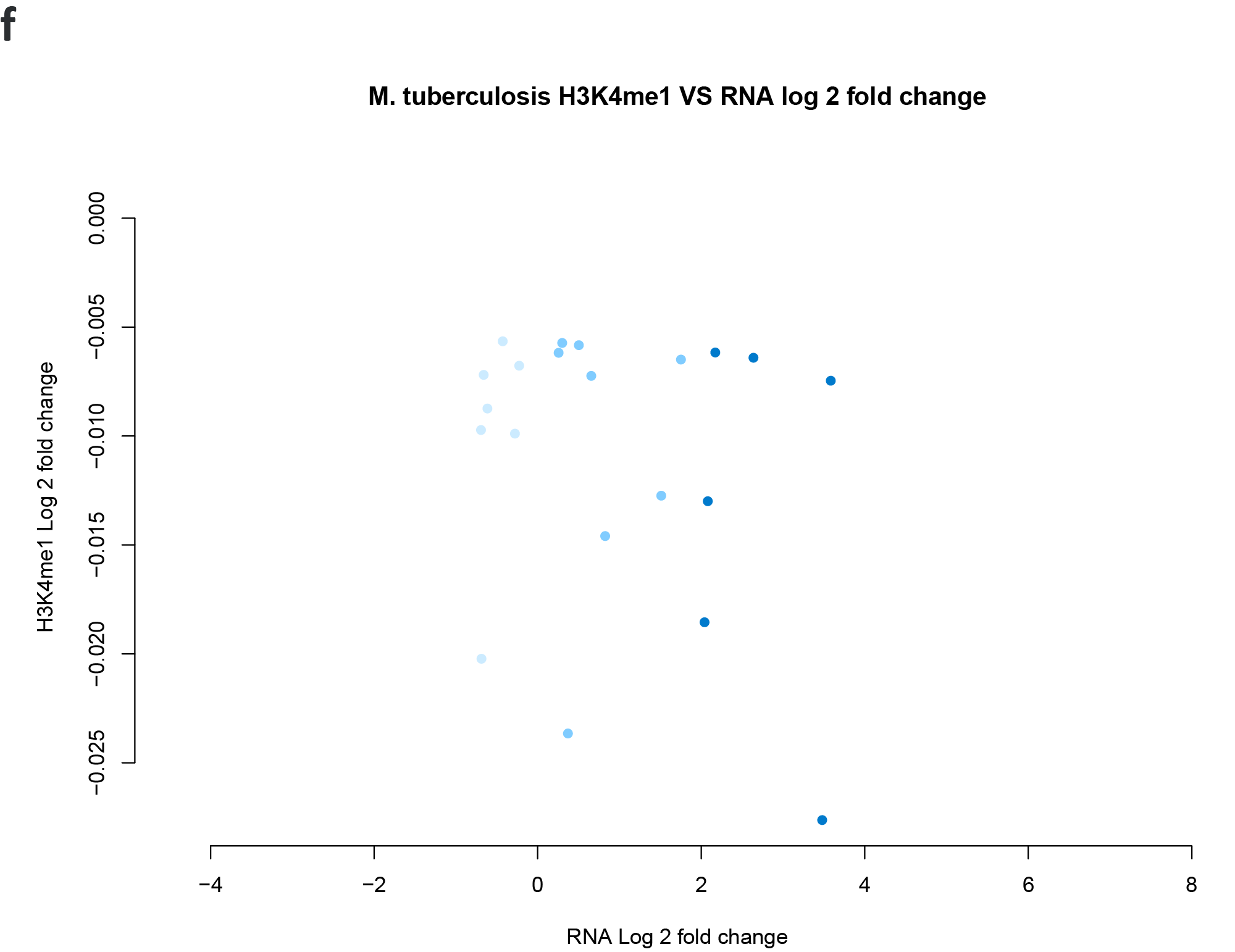

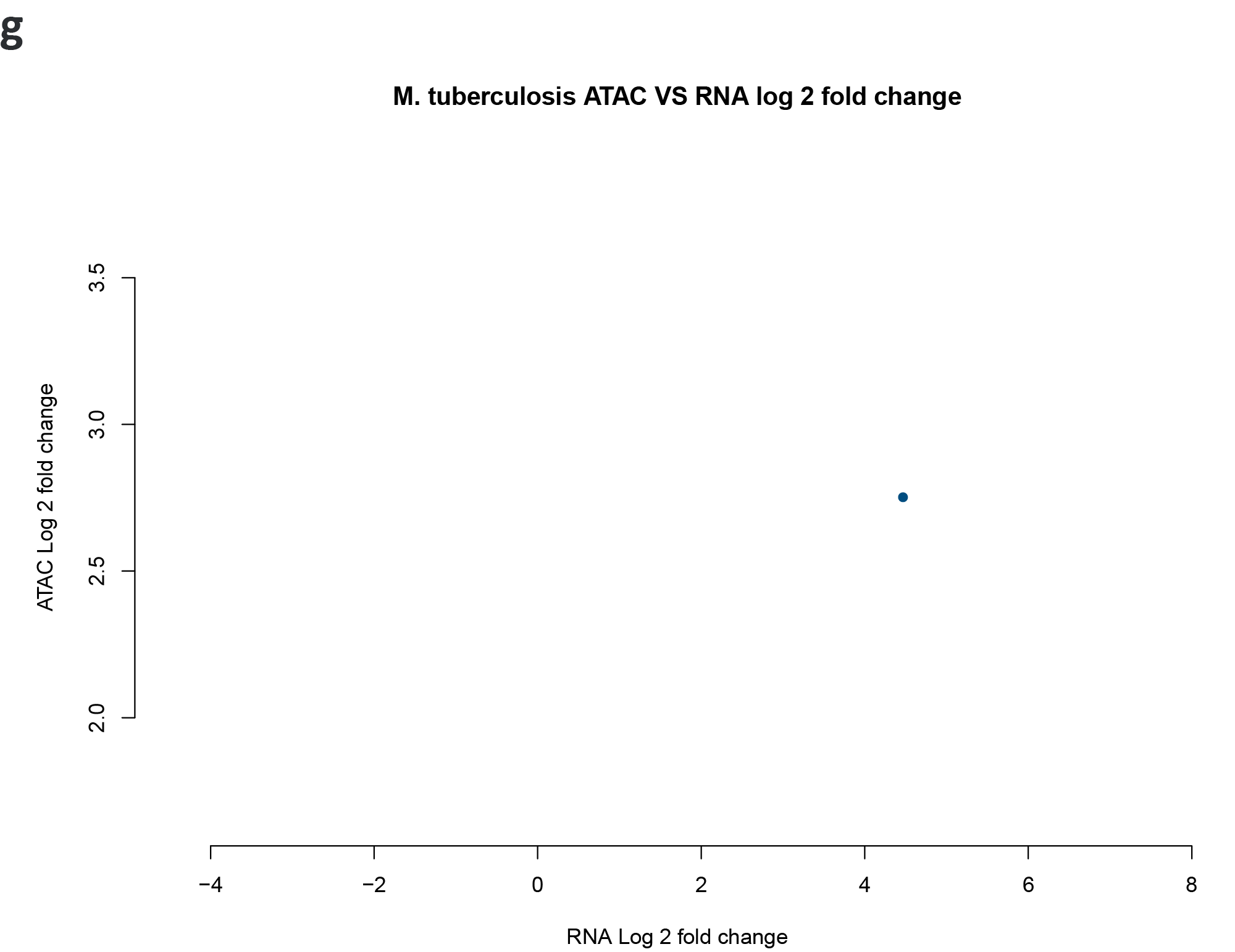

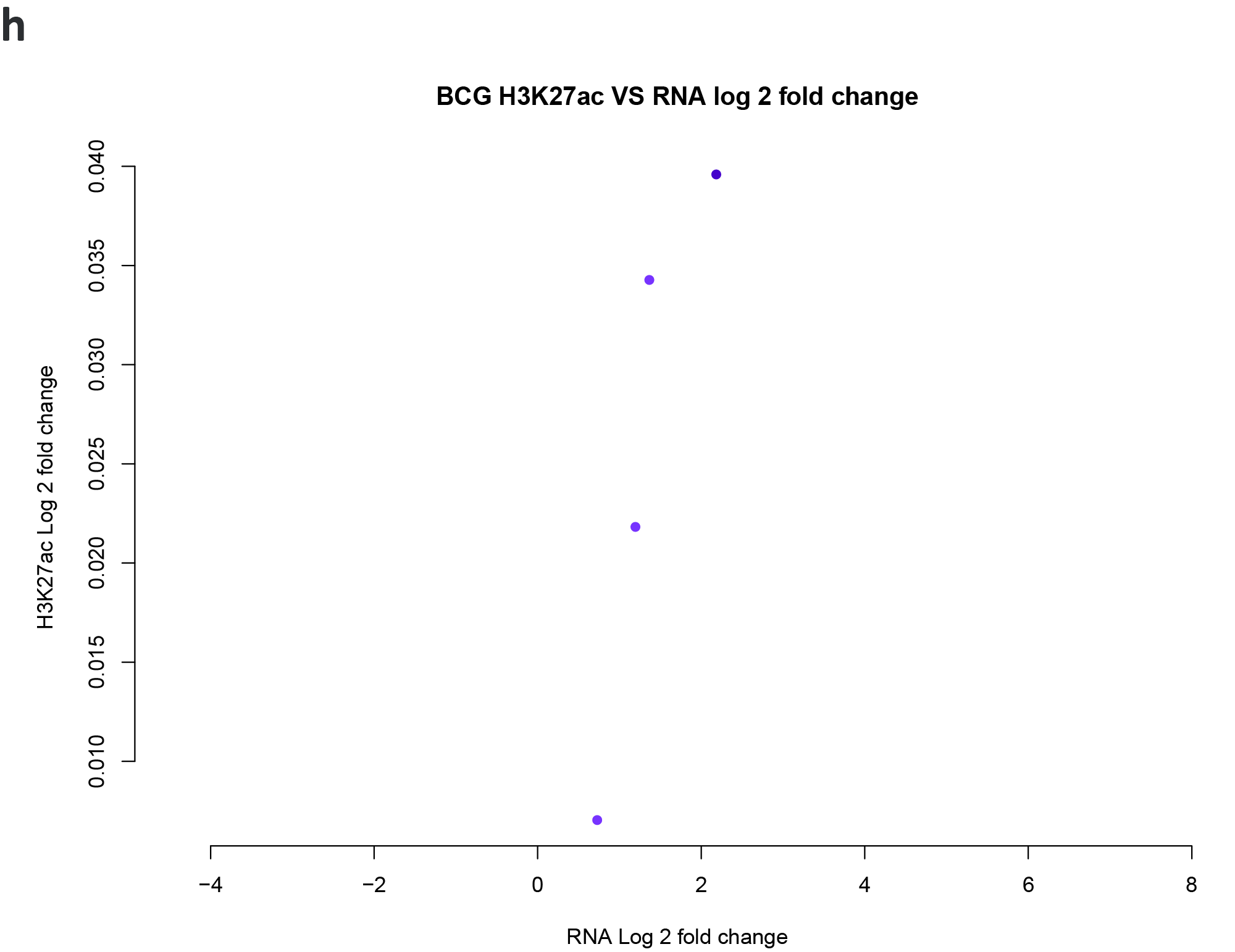

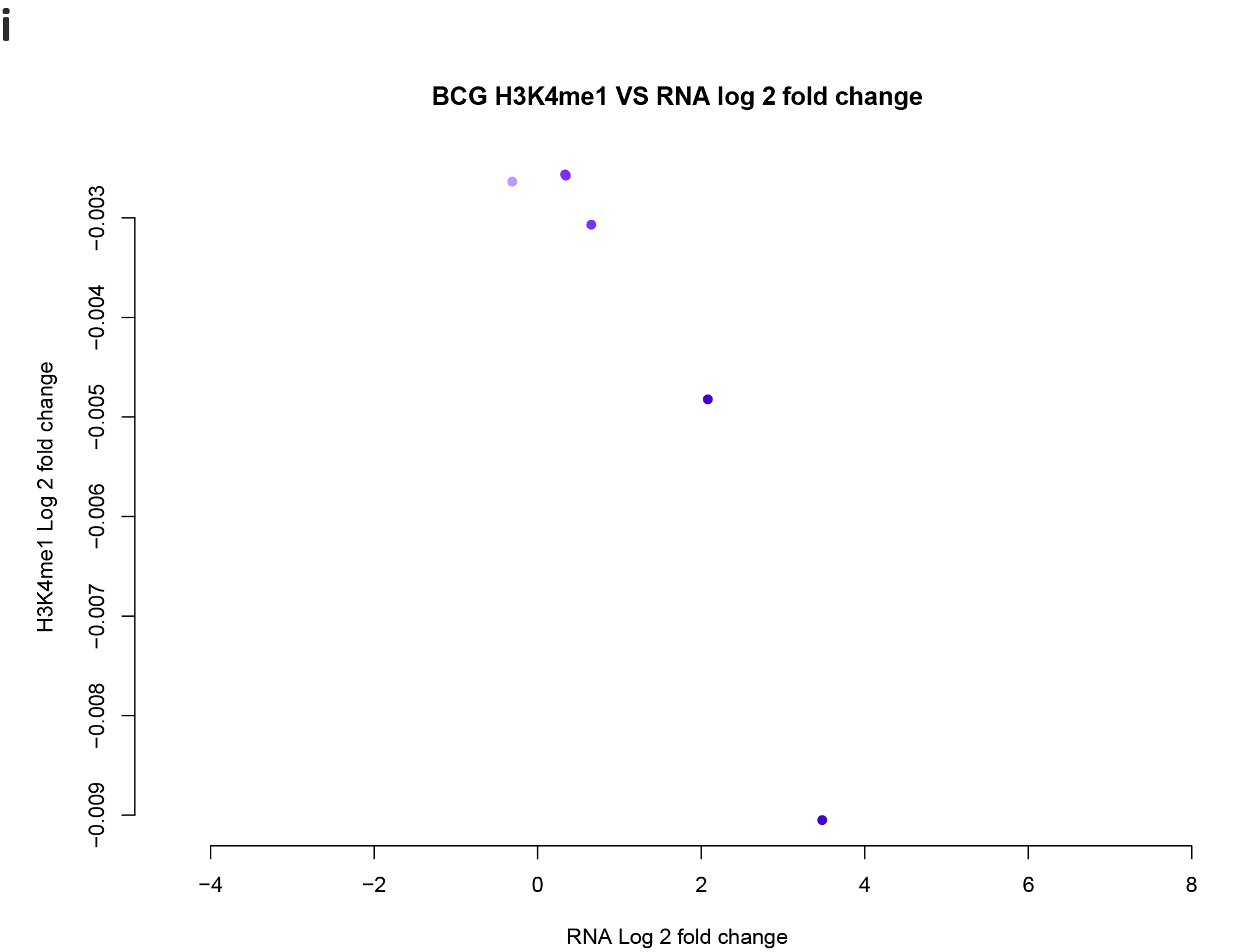

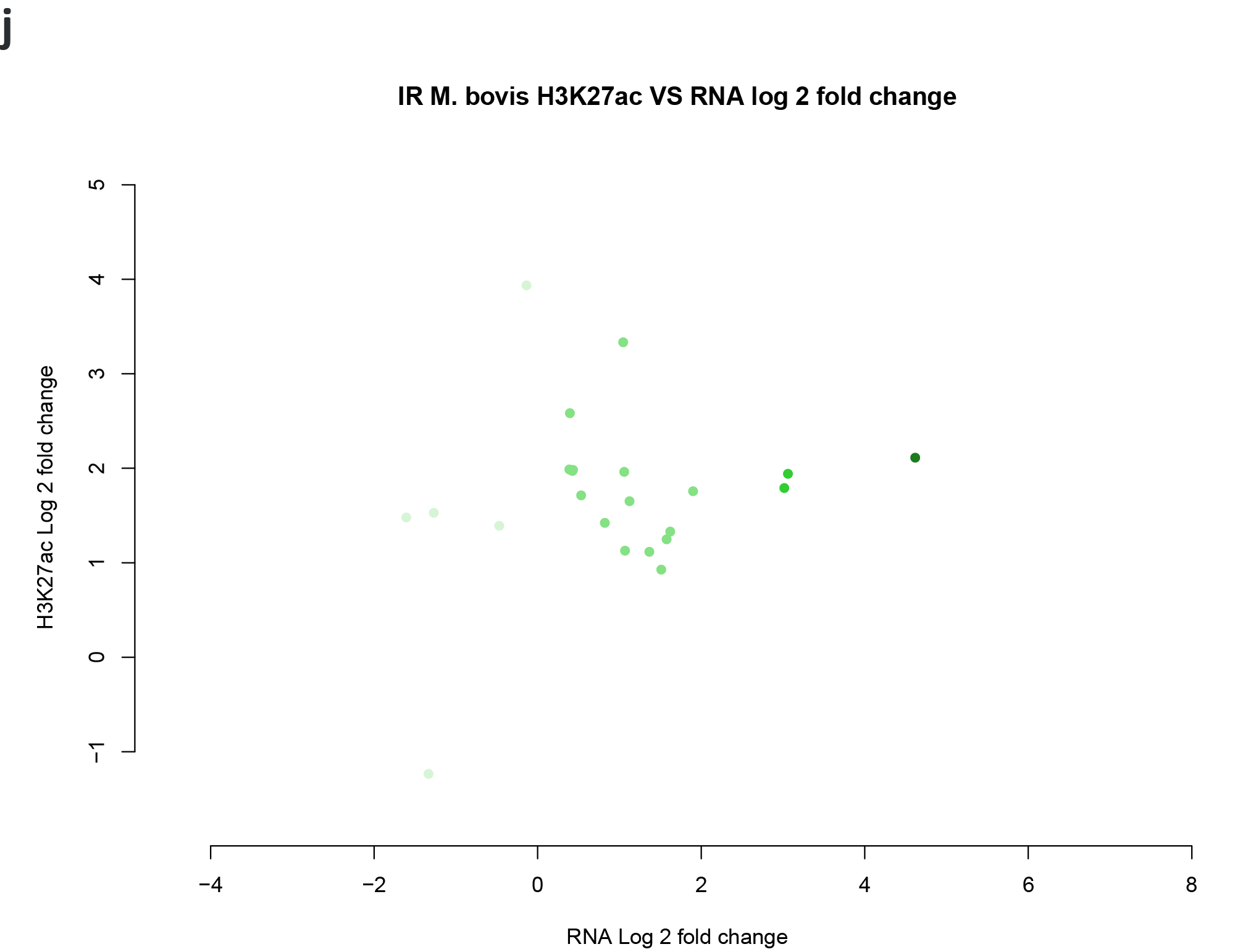

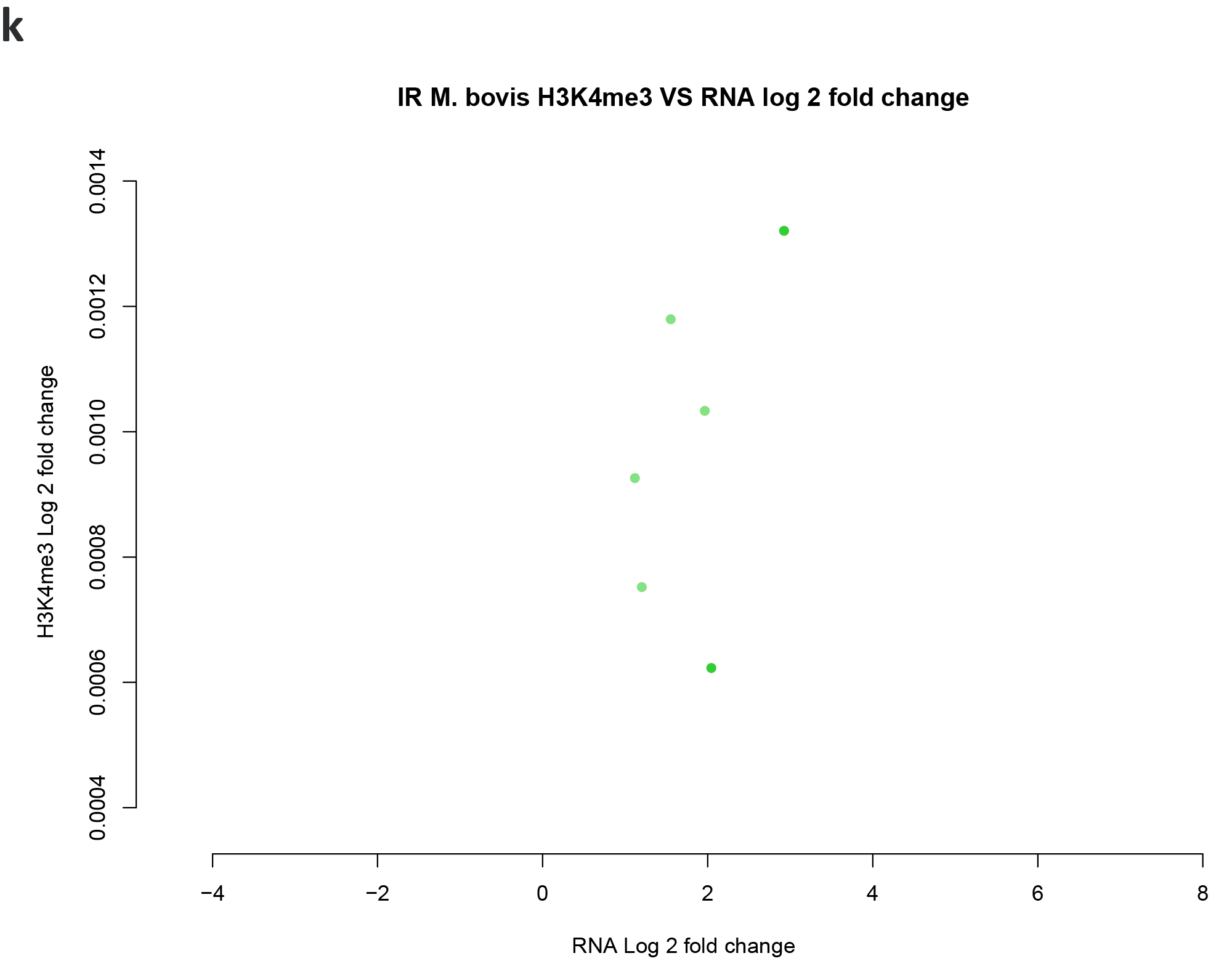

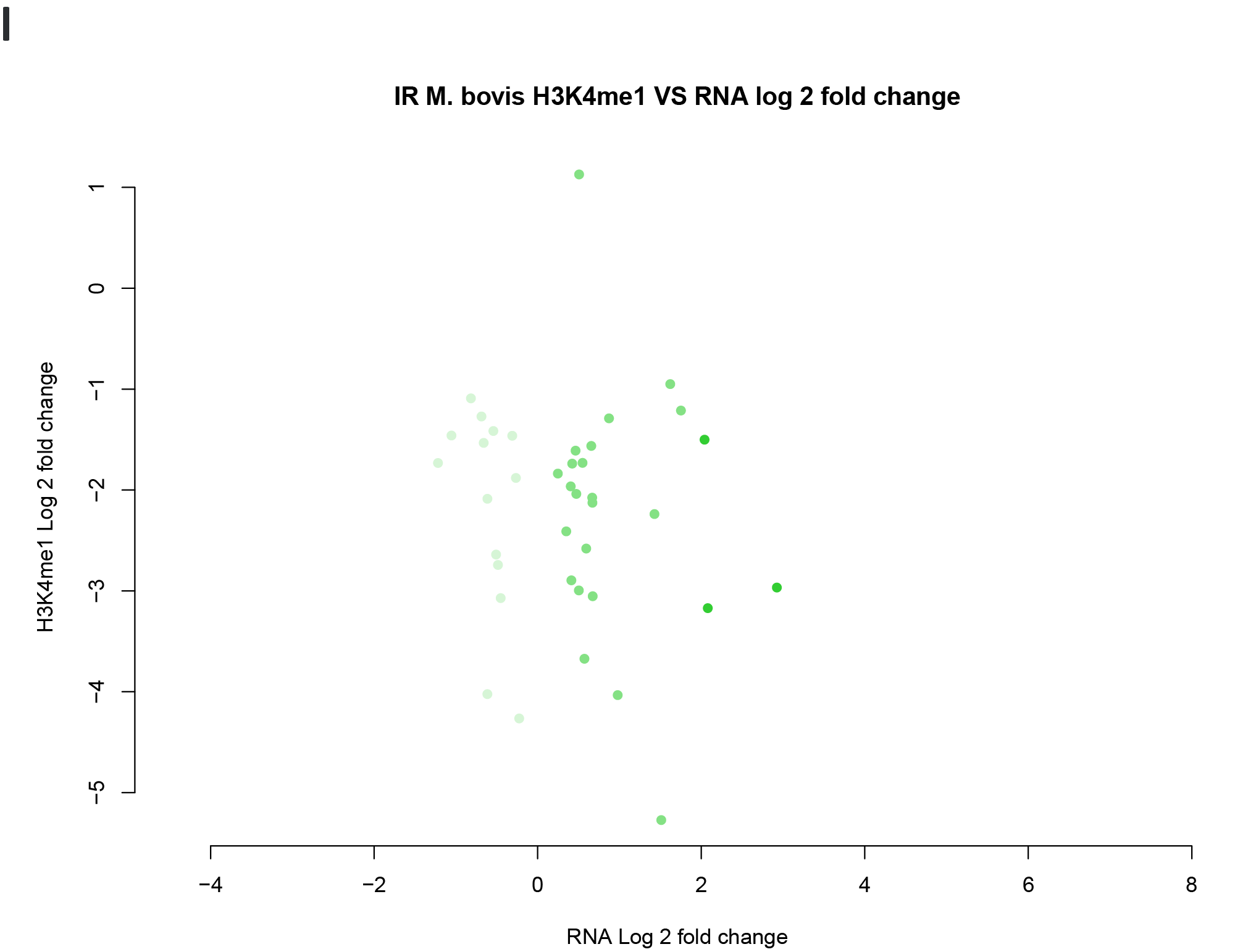

**Fig. S9** Network representation of the regulatory relationships among genes, histone modifications, and chromatin accessibility for the *Mycobacterium bovis* vs. control non-challenged bAM challenge contrast (MBO/CON) shown over the next four pages. Each circular node in the network corresponds to a gene observed to be co-located with a differential histone modification or chromatin accessibility difference (represented as square anchor hub nodes). The circular node colour shows if a gene was differentially expressed (upregulated: red; downregulated: blue; and yellow: not differentially expressed). The lines (edges) between nodes show if a histone modification difference/chromatin accessibility change was larger (determined by log_2_FC for peak sizes) in the infected bAM (red) or in the control non-challenged bAM (blue). Circular gene node size is determined by the number of connecting edges (degree). **a** Top left network quadrant. **B** Top right network quadrant. **c** Bottom left network quadrant. **d** Bottom right network quadrant.

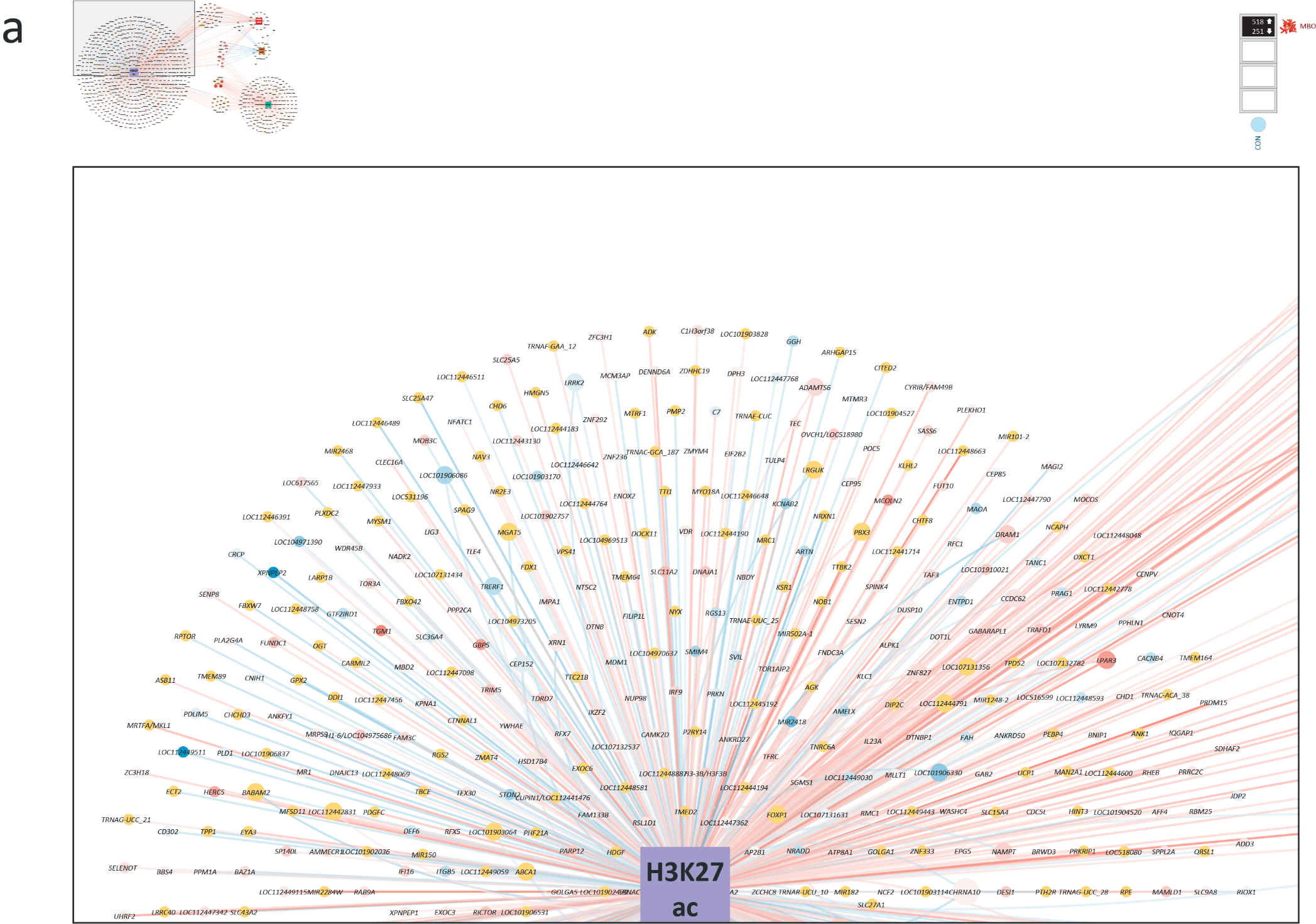

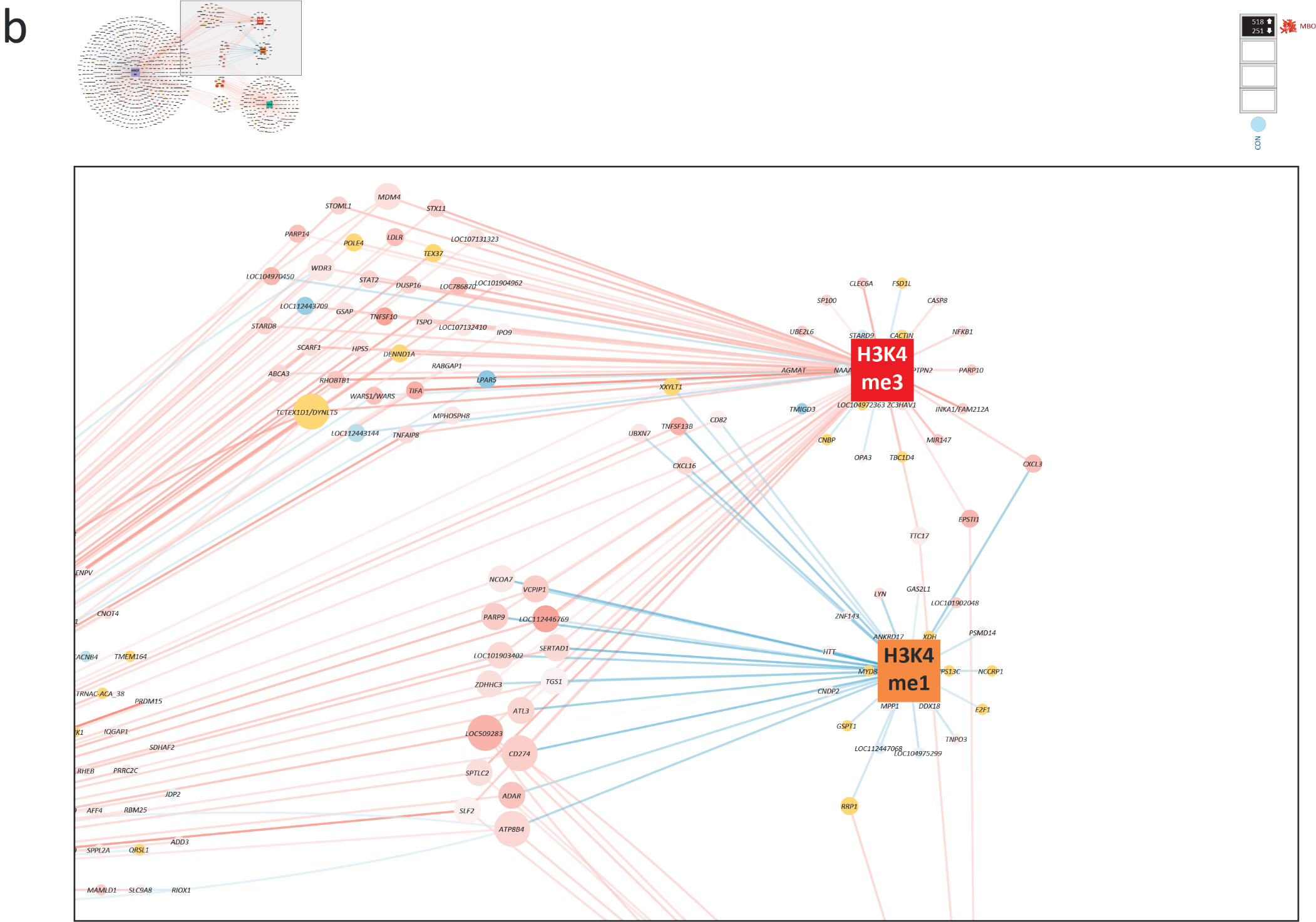

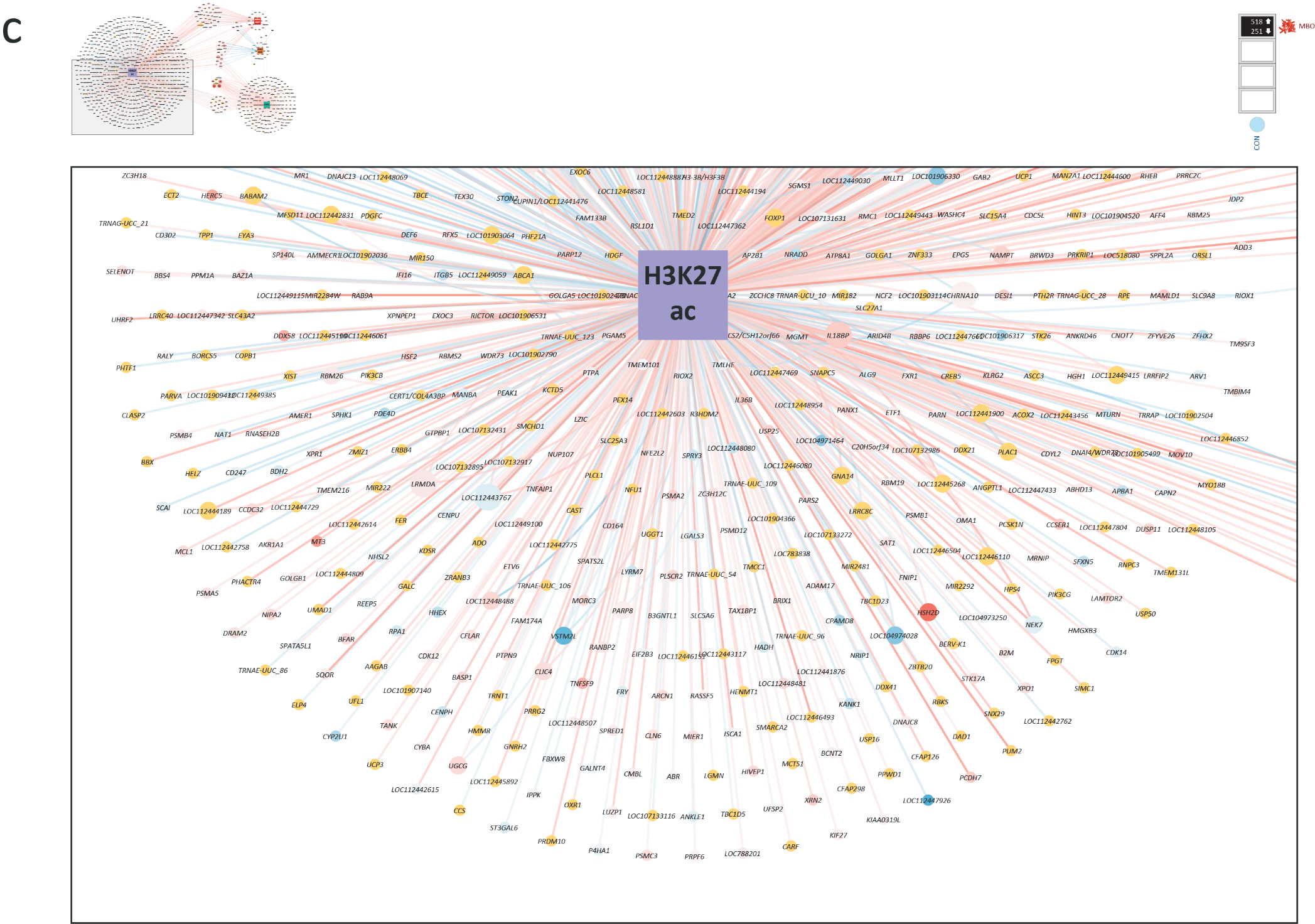

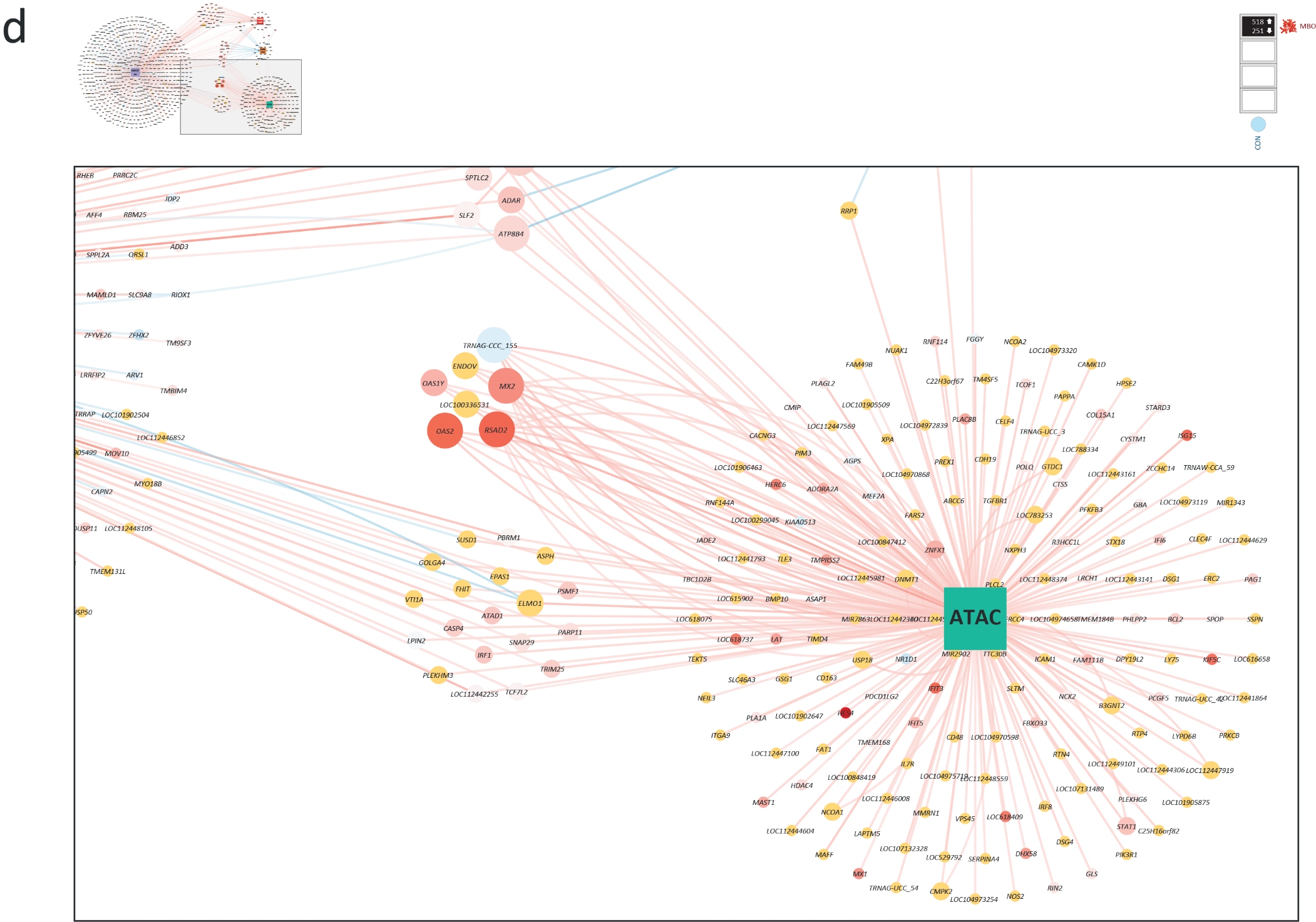

**Fig. S10** Gene expression changes and activation state for the Ingenuity Pathway Analysis (IPA®) *Role of RIG1-like receptors antiviral innate immunity* canonical pathway. Genes represented in pink were identified as differentially expressed genes (DEGs). DEGs and non-DEG genes shown in orange and blue were predicted to be upregulated and downregulated, respectively. **a** Pathway results for the *Mycobacterium bovis* vs control non-challenged bAM contrast (MBO/CON) using DEGs with an FDR-*P*_adj._ threshold of 0.00001. **b** Pathway results for the *M. bovis* vs irradiated (killed) *M. bovis* bAM contrast (MBO/IRR) using DEGs with an FDR-*P*_adj._ threshold of 0.001. **c** Pathway results for the *M. tuberculosis* vs irradiated (killed) *M. bovis* bAM contrast (MTU/IRR) using DEGs with an FDR-*P*_adj._ threshold of 0.05. **d** Pathway results for the *M. bovis* BCG vs control non-challenged bAM contrast (BCG/CON) using DEGs with an FDR-*P*_adj._ threshold of 0.05. **e** Pathway results for the irradiated (killed) *M. bovis* vs control non-challenged bAM contrast (IRR/CON) using DEGs with an FDR-*P*_adj._ threshold of 0.05.

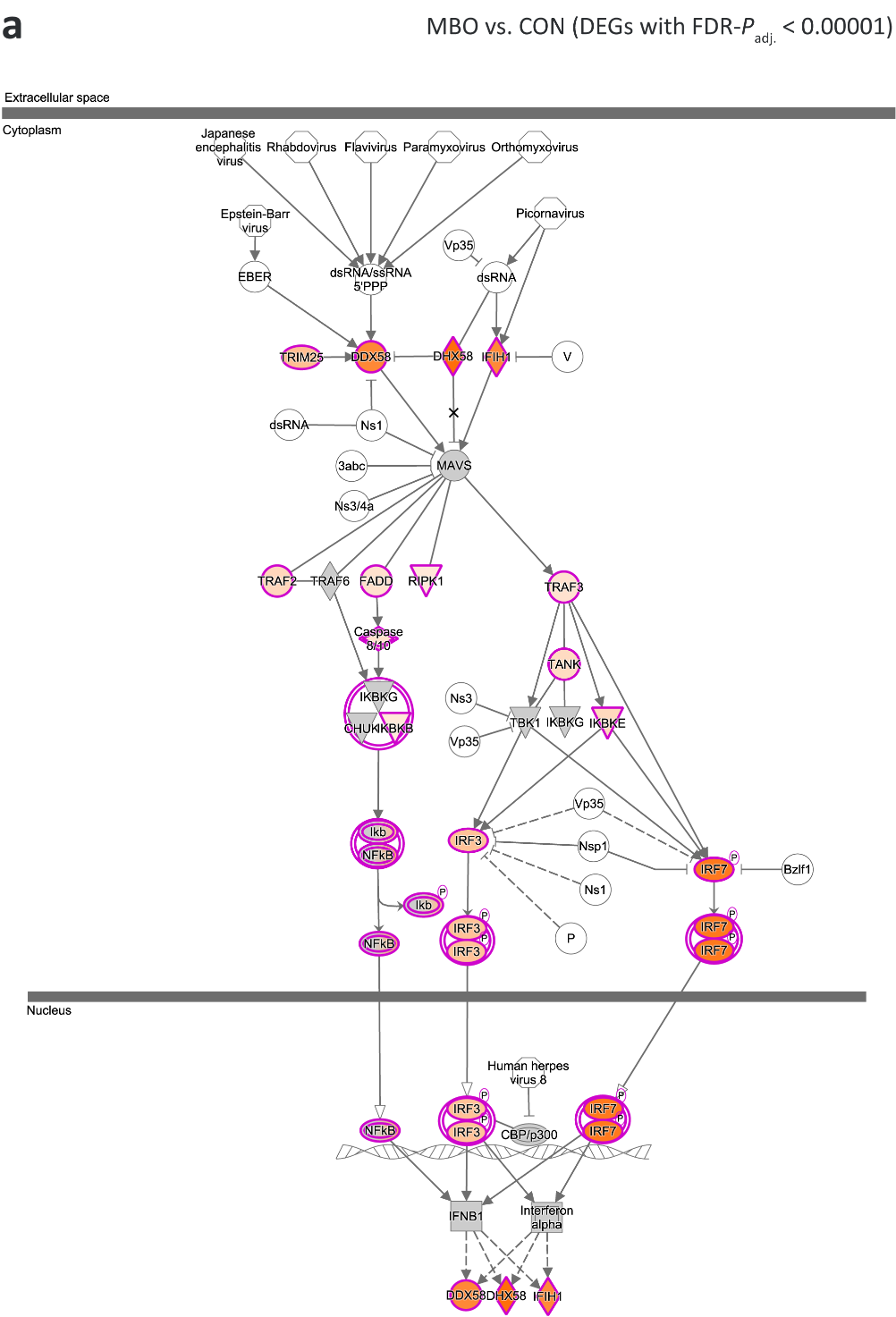

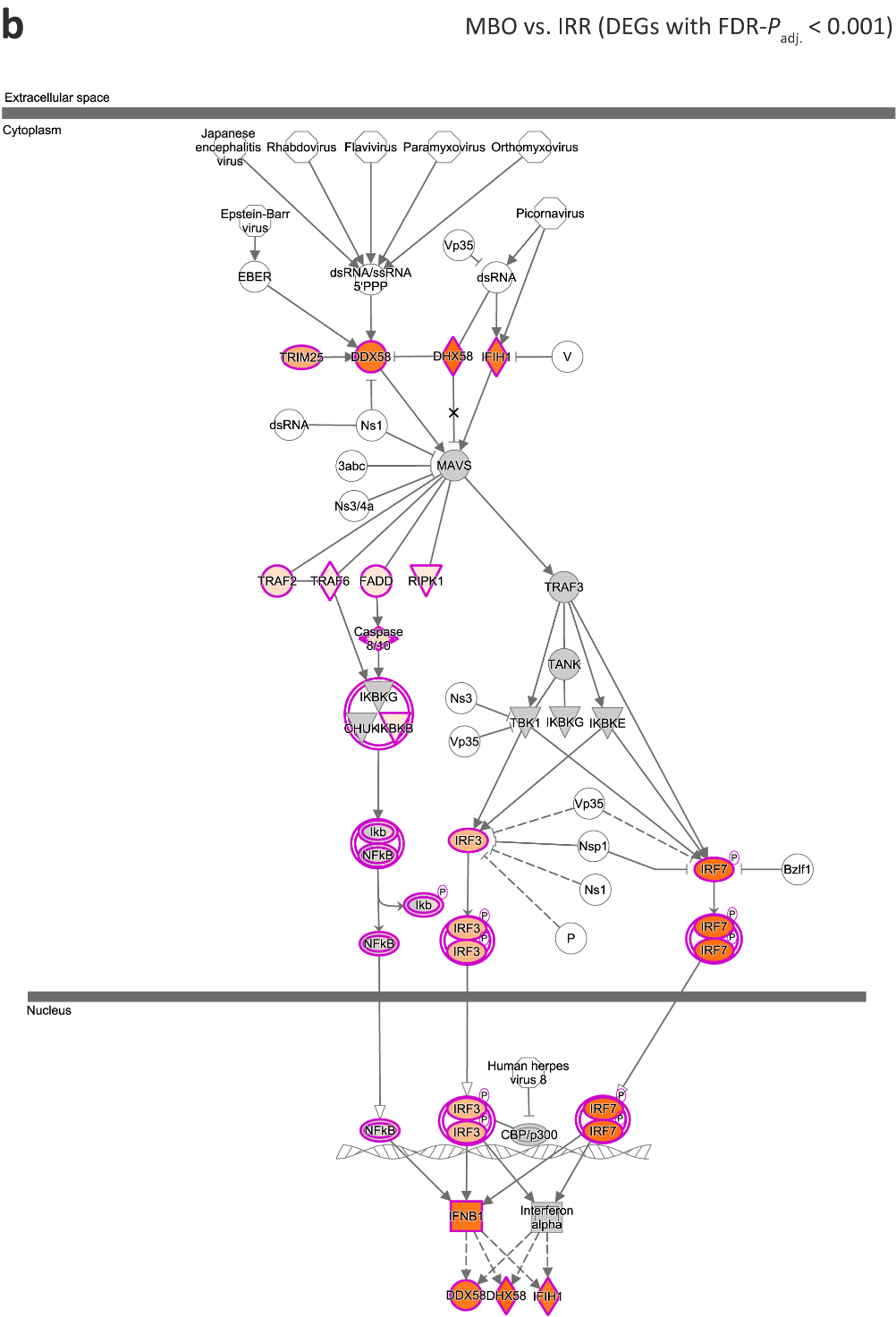

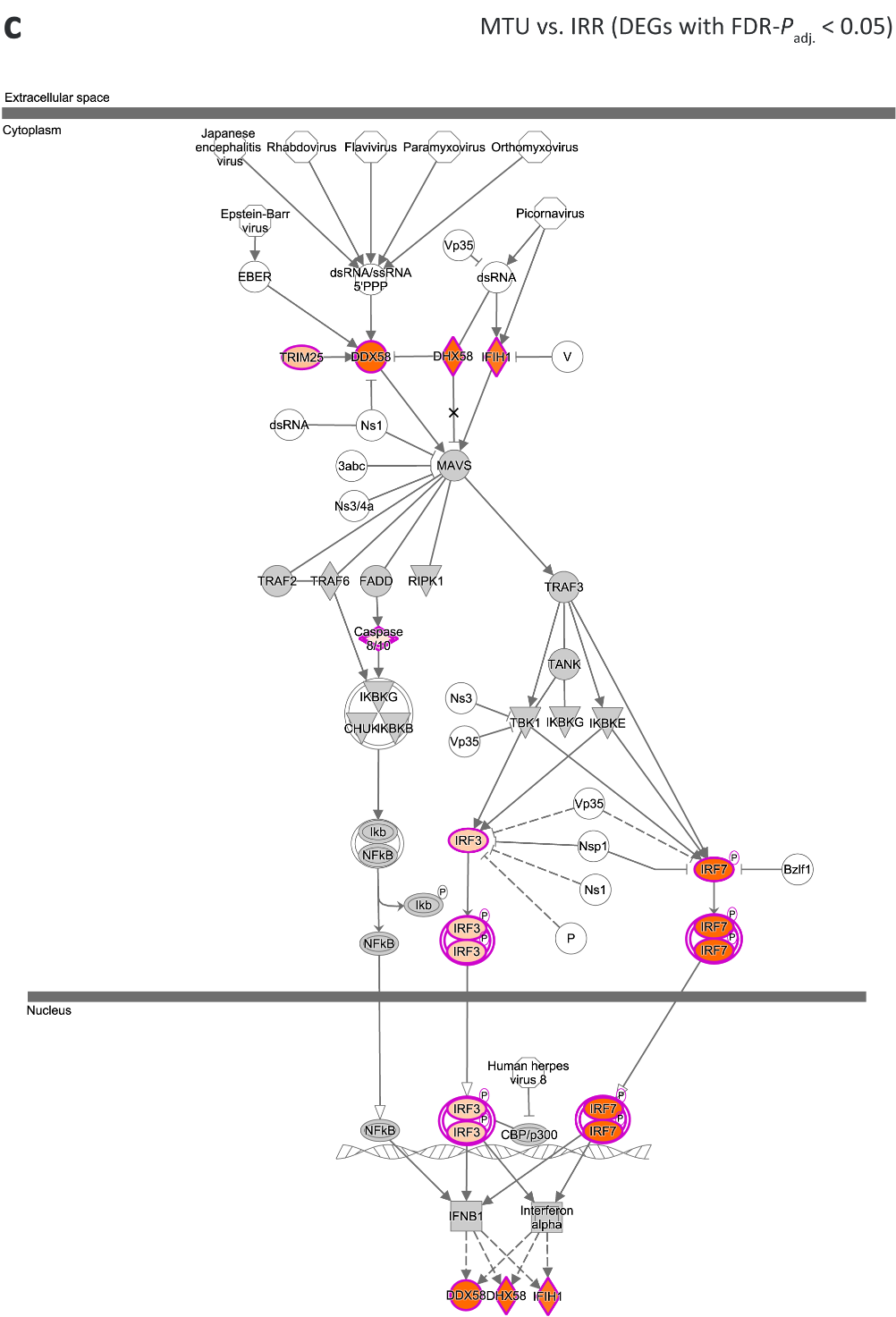

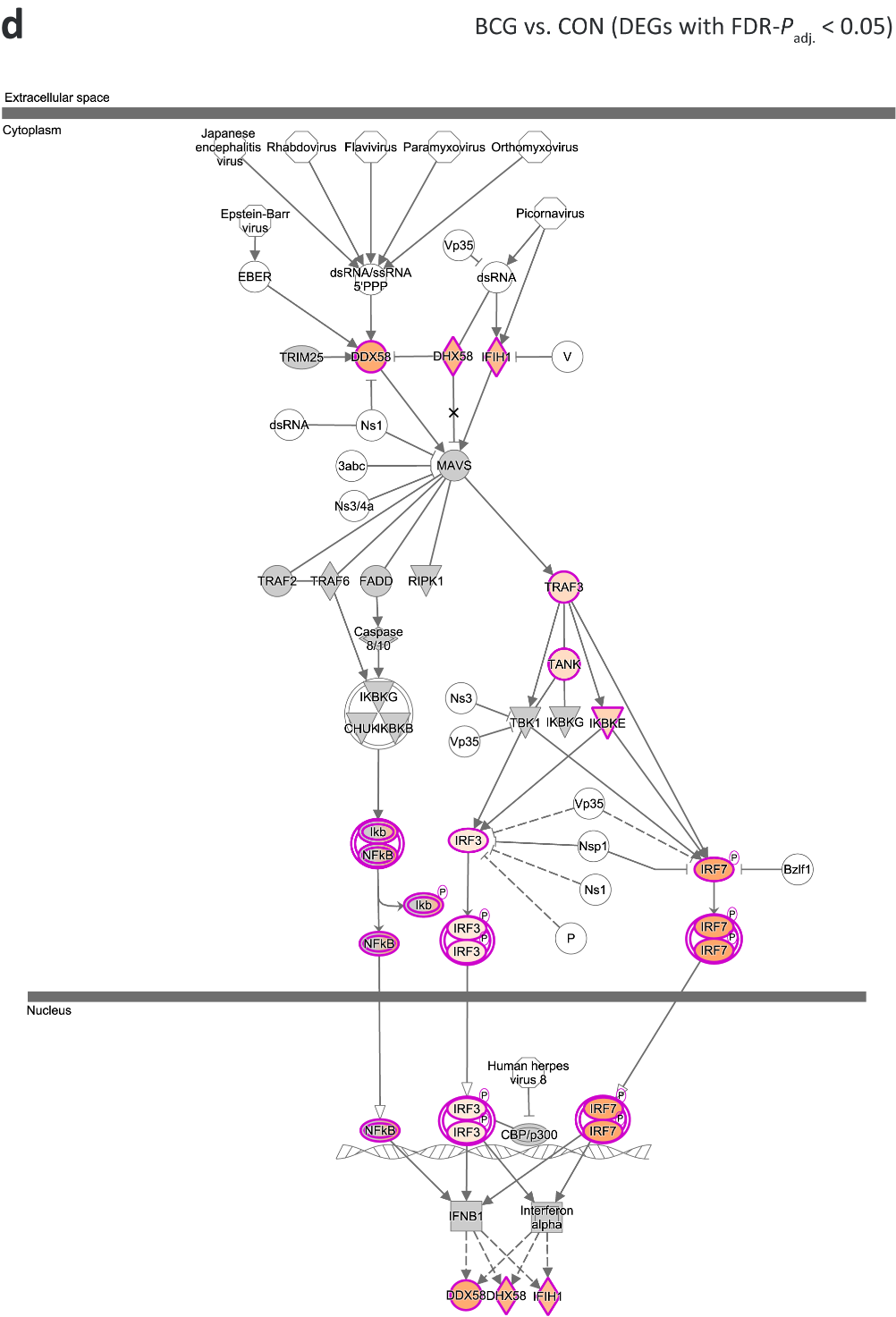

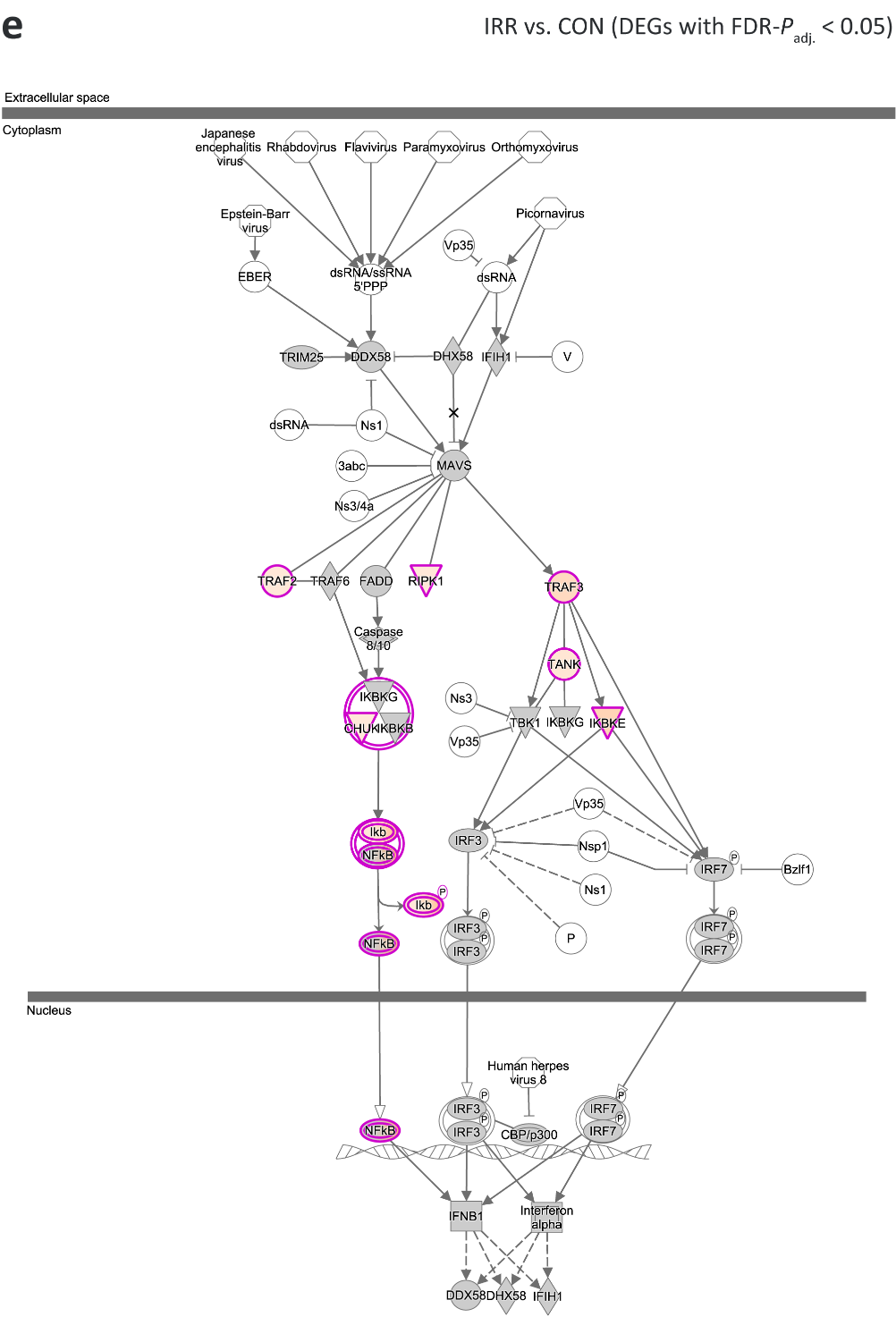
